# Long-term editing of brain circuits in mice using an engineered electrical synapse

**DOI:** 10.1101/2025.03.25.645291

**Authors:** Elizabeth Ransey, Gwenaëlle E. Thomas, Elias Wisdom, Agustin Almoril-Porras, Ryan Bowman, Elise Adamson, Kathryn K. Walder-Christensen, Jesse A. White, Dalton N. Hughes, Hannah Schwennesen, Caly Ferguson, Kay M. Tye, Stephen D. Mague, Longgang Niu, Zhao-Wen Wang, Daniel Colón-Ramos, Rainbo Hultman, Nenad Bursac, Kafui Dzirasa

## Abstract

Electrical signaling across distinct populations of brain cells underpins cognitive and emotional function; however, approaches that selectively regulate electrical signaling between two cellular components of a mammalian neural circuit remain sparse. Here, we engineered an electrical synapse composed of two connexin proteins found in *Morone americana* (white perch fish) – connexin34.7 and connexin35 – to accomplish mammalian circuit modulation. By exploiting protein mutagenesis, devising a new *in vitro* system for assaying connexin hemichannel docking, and performing computational modeling of hemichannel interactions, we uncovered a structural motif that contributes to electrical synapse formation. Targeting these motifs, we designed connexin34.7 and connexin35 hemichannels that dock with each other to form an electrical synapse, but not with other major connexins expressed in the mammalian central nervous system. We validated this electrical synapse *in vivo* using *C. elegans* and mice, demonstrating that it can strengthen communication across neural circuits composed of pairs of distinct cell types and modify behavior accordingly. Thus, we establish ‘<u>L</u>ong-term <u>in</u>tegration of <u>C</u>ircuits using conne<u>x</u>ins’ (LinCx) for precision circuit-editing in mammals.

---

Electrical synapses enable direct flow of ions and small molecules between two cells and play a prominent role in broadly coupling electrical activity in multiple organs including the brain^1–3^. They comprise multiple gap junction channels, each composed of two docked hemichannels embedded in the membranes of two touching cells. Each hemichannel is an oligomer consisting of six monomeric proteins called connexins (Cxs), of which there are 21 isoforms in humans ^4,5^. Most Cxs can form single-isoform hemichannels that dock with themselves to create homotypic gap junctions (Fig. 1A, left). Neural circuit-editing using gap junctions is well-established in *C. elegans*^6–9^. *C. elegans* do not express Cxs; thus, heterologous expression of the vertebrate connexin36 (Cx36) across two connected *C. elegans* neurons results in the formation of an electrical synapse that does not interact with endogenous *C. elegans* gap junction proteins (i.e., innexins). Prior work has successfully implemented this editing approach to modify circuit physiology in multiple behavioral contexts including migration of *C. elegans* in response to various chemicals and temperatures^7–10^.

**Figure 1:**
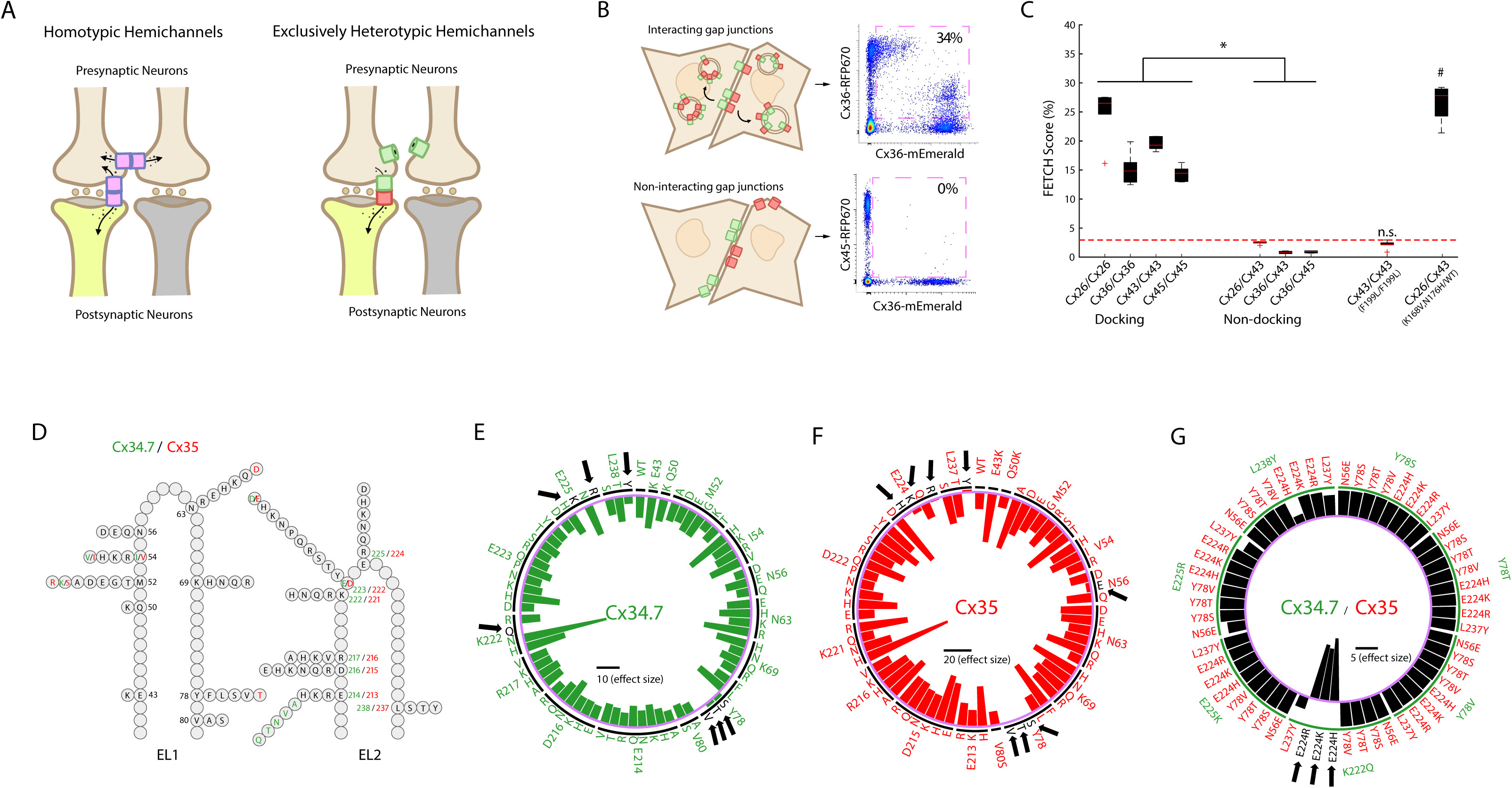
Screen to identify a mutant connexin hemichannel pair that exhibits exclusively heterotypic docking. A) (left) Schematic outlining the limitation of introducing heterologous wild type connexin (Cx) hemichannels (pink rectangles) as a method for modulating precise neural circuits composed by brown and yellow neurons. Note that Cx hemichannels yield an off-target electrical synapse between the pre-synaptic neurons and, thus, off-target modulation of other circuits. (right) Putative strategy for deploying exclusively heterotypic docking hemichannels (green and red rectangles) to selectively modulate precise neural circuits. Note the rectification of this putative gap junction. B) Depiction of red and green fluorescence exchange profiles (left) and representative flow cytometry plots (right) for hemichannel pairs with (Cx36/Cx36; top) and without (Cx36/Cx45; bottom) docking compatibility. Pink dashed squares in flow cytometry plots highlight the portion of transfected cells labeled by two distinct fluorescent proteins. **C)** Portion of dual fluorescent-labeled cells for Cx pairs with known docking compatibility profiles (* = P<0.05 for docking vs. non-docking pairs ; #=P<0.05 using two tailed t-test compared to Cx43/Cx43). The red line corresponds with a distribution of Cx pairs with well-established docking incompatibility. For the box and whisker plots, the central mark is the median, the edges of the box are the 25th and 75th percentiles, the whiskers extend to the most extreme datapoints the algorithm considers to be not outliers, and the outliers are plotted individually as “+.” D) Schematic of *Morone Americana* Cx34.7 and Cx35 extracellular loop mutations used to screen for heterotypic-exclusive hemichannels. Positions and mutations unique to Cx34.7 and Cx35 are shown in green and red, respectively. Positions and mutations common to both proteins are shown in black. **E-F)** Circular plot showing homotypic FETCH results for E) Cx34.7 and F) Cx35 mutations. Circular bar graphs show the Cohen d’s effect size (portion of dual-labeled cells) of homotypic mutant combinations relative to the heterotypic pairing of human Cx36 and Cx45 which fails to dock. Black line in the center of the circles corresponds to scale bar for effect sizes. The names of targeted residues are listed around the rim of the circle. The substituted amino acids are listed just interior. The intermittent black circle segregates each targeted residue, and the light purple circle corresponds to zero effect size. Mutations (in black letters) that disrupted docking are also highlighted by black arrows. For example, the black arrow on the left of panel E highlights a mutation from lysine (K) to glutamine (Q) at the 222nd residue that disrupted docking of Cx34.7. **G)** Heterotypic FETCH results for Cx34.7 (green) and Cx35 (red) mutant protein combinations. Bar graphs show the effect size of homotypic mutant combinations relative to the wild type Cx34.7 and Cx35 pair. The purple circle corresponds to an effect size of zero. The green intermittent circle corresponds to the Cx34.7 mutations identified in green around the rim of the plot. Black line in the center of the circle corresponds to scale bar for effect size.

The potential for using gap junctions to repair dysfunctional circuits has been advanced in *C. elegans* as well^10,11^, as shown in experiments that utilized circuit-editing to restore normal behavior in animals with induced circuit disruptions. Nevertheless, this work also highlighted a significant challenge to employing gap junctions to edit select circuits in higher-complexity organisms. Specifically, when Cx36 was expressed in two sensory neurons of the same cell-type and formed homotypic gap junctions, otherwise normal *C. elegans* showed disrupted behavior in response to olfactory cues^10^. Because vertebrate brains are composed of many more cells of the same cell-type than *C. elegans*, the connexins’ ability to form homotypic gap junctions across more cells has the potential to dramatically diminish the precision of this circuit-editing approach for mammals (e.g., off-target modulation, Fig. 1A, left), yielding even greater behavioral disruption. Moreover, heterologous expression of Cxs from other species in mammals might lead to gap junctions composed of both exogenous and endogenous Cxs, producing undesired connections that may impair normal neural circuit functions.

While nearly all Cxs can form homotypic channels, several Cx isoforms are capable of docking with other Cx isoforms to generate heterotypic channels (Fig. 1A, right)^3,12^. We reasoned that by identifying the mechanisms underlying Cx docking interactions^13^, we could design a hemichannel pair biased towards heterotypic gap junction formation. We also reasoned that we could engineer this pair to be docking-incompetent with the major Cxs endogenous to the mammalian CNS, yielding a precise approach for regulating electrical flow between distinct cell types.

*Morone americana* (white perch fish) expresses two homologs of mammalian neuronal Cx36 – connexin 34.7 (Cx34.7) and connexin 35 (Cx35) – that create a heterotypic gap junction^14^. Interestingly, this electrical synapse exhibits channel-level rectification in the Cx34.7 to Cx35 direction when expressed in *Xenopus* oocytes^14^. The orthologues of Cx34.7 and Cx35 in the Goldfish (*Carassius auratus*) CNS also create a heterotypic gap junction that shows circuit-level rectification in the Cx34.7 to Cx35 direction^15^. Given that Cx34.7 and Cx35 form heterotypic gap junctions with inherent directionality^15^, conduct currents capable of triggering action potential firing^14^, and are potentially amenable to modification of biophysical properties through amino acid sequence mutations, we chose them as building blocks to engineer a desired new electrical synapse for editing mammalian circuits.

## Results

### *In vitro* assay of connexin protein docking

To establish a method for evaluating Cx docking specificity and ultimately engineering our electrical synapse, we exploited the natural cell processes for trafficking docked Cxs. In mammalian cells, Cx hemichannels can be removed from the membrane via a coordinated endo- and exocytic process, resulting in internalization of fully docked gap junctions within double bi-layer vesicles (Fig. 1B, top left)^16–21^. Indeed, it has previously been shown that by tagging Cx hemichannels expressed by one cell with a fluorescent protein, internalization into adjacent, non-fluorescence expressing cells can be visualized (Fig. 1B, top)^21,22^.

Our approach utilized populations of HEK293FT cells that expressed individual Cx as either mEmerald-, or RFP670 fluorescent fusion proteins (Fig. 1B). We then co-plated and incubated the HEK293FT cells that express Cx counterparts. Finally, we evaluate their fluorescence exchange via flow cytometry (see Supplemental Fig. S1). Since docking is prerequisite for internalizing fluorescently tagged connexins expressed by other cells, docking can be quantified as the portion of transfected cells that are labeled by dual fluorescence in the co-plated sample (Fig 1B-C). We first established the utility of our assay (termed FETCH - flow <u>e</u>nabled tracking of <u>c</u>onnexosomes in <u>H</u>EK cells) by testing the well-characterized Cx proteins Cx26, Cx36, Cx43, and Cx45. Given that each these proteins is known to be capable of homotypic docking ^21–24^, we tested them under homotypic pairings (FETCH = 24.8±1.8%, 15.2±1.1%, 19.5±0.4%, and 14.4±0.5% dual-labeled cells for Cx26/Cx26, Cx36/Cx36, Cx43/Cx43, and Cx45/Cx45, respectively; mean±s.e.m). We also tested them in paired combinations for which was there was prior evidence of heterotypic docking-incompatibility (e.g., Cx26/Cx43, Cx36/Cx43, and Cx36/Cx45; FETCH = 2.5±0.1%, 0.8±0.1%, and 0.9±0.1% for Cx26/Cx43, Cx36/Cx43, and Cx36/Cx45, respectively; see Fig. 1C)^12^. Importantly, the proportion of dual-labeled cells in the population of docking-compatible vs. docking-incompatible pairs was statistically different (T_40_ =14.5; P=1.6×10^-17^ using unpaired t-test), establishing that FETCH could be used to broadly assess Cx hemichannel docking compatibility.

Second, we probed the utility of FETCH by testing two Cx mutations that impact gap junction formation. Specifically, we tested a Cx43_F199L_ mutation previously shown to disrupt trafficking to the cell membrane^25^ in homotypic configuration. We also evaluated a Cx26_K168V,N176H_ mutant known to confer heterotypic docking compatibility with wild type Cx43^26^. The Cx43_F199L_ mutant did not exhibit a level of fluorescence exchange that was higher than our non-docking heterotypic pairs (FETCH=2.1±0.3% for Cx43_F199L_/Cx43_F199L_; T_22_ =2.0; P=0.06 compared to pooled distribution of Cx26/Cx43, Cx36/Cx43, and Cx36/Cx45 using a two-tailed unpaired t-test; FETCH=4.4±0.6%% for the pooled distribution; see Fig. 1C, red line), while the Cx26_K168V, N176H_ mutant showed fluorescence exchange that was significantly higher when assayed against Cx43 (FETCH=26.6±1.3% for Cx26_K168V, N176H_/Cx43; T_22_ =32.7; P=3.9×10^-20^; see Fig. 1C). Thus, we established that our FETCH assay could be used to identify Cx mutations that disrupt or enable docking compatibility.

### *In vitro* assessment of Cx34.7 and Cx35 mutant docking

We used FETCH to assay a library of Cx34.7 and Cx35 mutations for their impact on hemichannel docking. Though the precise interactions that guide hemichannel docking are incompletely characterized for the majority of Cxs, structure–function and sequence analyses indicate that both extracellular loops (EL1 and EL2) play a role in hemichannel docking^13,27,28^. To identify Cx34.7 and Cx35 variants that are unable to form homotypic gap junctions, we introduced ∼70 individual mutations at sixteen positions on both extracellular loops (ELs) of each Cx (see methods for mutant library design; Fig. 1D and Supplemental Fig. S2). We then compared the homotypic pairing FETCH scores for the mutants to a docking-incompatible heterotypic pair (e.g., Cx36 paired with Cx45, see Fig. 1E-F)^29^. We identified several homotypic non-docking mutant proteins for Cx34.7 (Y78S, Y78T, Y78V, E225K, E225R, L238Y and K222Q) and Cx35 (N56E, Y78V, Y78S, Y78T, E224H, E224K, E224R, and L237Y; see Fig. 1E-F). Next, to identify mutant protein pairs that exhibit exclusively heterotypic docking, we tested these Cx34.7 and Cx35 mutants against each other and compared their FETCH scores to wild type Cx34.7/Cx35. Strikingly, we discovered three Cx mutant pairs whose FETCH scores were higher than the scores observed for wild type Cx34.7_WT_/Cx35_WT_ gap junctions. These results provided evidence of mutant pairs (Cx34.7_K222Q_ with either Cx35_E224H_, Cx35_E224K_, or Cx35_E224R_) with intact heterotypic, but reduced homotypic, docking (Fig. 1G).

Since our long-term objective was to develop a precise modulation approach that would be amenable for use in the mammalian nervous system, we also probed whether the four identified mutant proteins could dock with the major connexins expressed by mammalian neurons and astrocytes – specifically Cx36 and connexin43 (Cx43), respectively^30,31^. We again used FETCH, and we compared the scores against a broad population of non-docking pair replicates (see methods).

None of the mutant proteins interacted with human Cx43 (FETCH=1.3±0.1%, 0.4±0.1%, 0.5±0.1%, and 0.5±0.1%; T_96_= 0.29, 1.41, 1.36, and 1.28; P=0.61, 0.92, 0.91 and 0.90 for Cx34.7_K222Q_/Cx43, Cx35_E224H_,/Cx43, Cx35_E224K_/Cx43, and Cx35_E224R_/Cx43, respectively; N=6 replicates for all experimental Cx pairs). Cx35_E224K_ and Cx35_E224R_ also failed to interact with human Cx36; however, Cx34.7_K222Q_ and Cx35_E224H_ formed heterotypic gap junctions with human Cx36 (FETCH=22.8±1.9%; T_91_=-24.4; P=3.4×10^-43^ for Cx34.7_K222Q_/Cx36; FETCH=5.9±1.1%, 0.8±0.1%, and 0.6±0.1%; T_96_=-5.50, 0.89, and 1.24; P=1.6×10^-7^, 0.81, and 0.89 for Cx35_E224H_/Cx36, Cx35_E224K_/Cx36, Cx35_E224R_/Cx36, respectively). Thus, while Cx35_E224K_ and Cx35_E224R_ both showed docking incompatibility with Cx36 and Cx43, and neither showed homotypic docking, we failed to identify an effective Cx34.7 partner that did not dock with Cx36 using single-point mutagenesis.

### Integrating *in vitro* analysis and *in silico* modeling to create a putatively selective Cx34.7 and Cx35 pair

We utilized homology modeling and FETCH analysis to design a new Cx34.7 mutant that does not dock with endogenous Cx43 or Cx36, and to design its Cx35 heterotypic docking partner. Briefly, we first developed computational models of wild type and mutant Cx34.7 and Cx35 hemichannels under homotypic and heterotypic pairings. We then validated the computational model by comparing the key residues predicted to underlie hemichannel docking against the docking characteristics we measured for these mutants using FETCH. We also modeled their docking interactions with Cx36. Next, we utilized the insights from all our residue-wise interaction models to computationally design Cx34.7 and Cx35 hemichannels that would dock heterotypically only with each other. Finally, we generated these proteins and confirmed their docking characteristics *in vitro* using FETCH (Supplemental Fig. S3).

First, to model the docking interactions between Cx34.7 and Cx35 hemichannels, we ran molecular dynamics simulations of homotypic and heterotypic pairs of wild type and mutant Cx34.7 and Cx35 proteins^32,33^. We found large negative interaction energies involving residues E214, K222, E223, and E225 in wild type Cx34.7 and residues E213, K221, D222, E224 in wild type Cx35 for both the homotypic and heterotypic docking simulations. These large negative interaction energies were suggestive of salt bridges that stabilize both homotypic and heterotypic docking interactions, consistent with our FETCH screening in which charge-swapping mutations (i.e., positive charge to neutral and negative charge to positive at positions Cx34.7-K222 and Cx35-E224, respectively) disrupted docking. Integrating these results, we identified a common interaction motif for both Cx34.7 and Cx35 consisting of three negative residues (E214/E213, E223/D222, and E225/E224 for Cx34.7 and Cx35, respectively), and a positive residue (K222/K221 for Cx34.7 and Cx35, respectively, see Fig. 2A-C). This interaction motif was consistent with a previously proposed theoretical framework where four residues underlie the docking specificity of most Cx hemichannels^13^.

**Figure 2:**
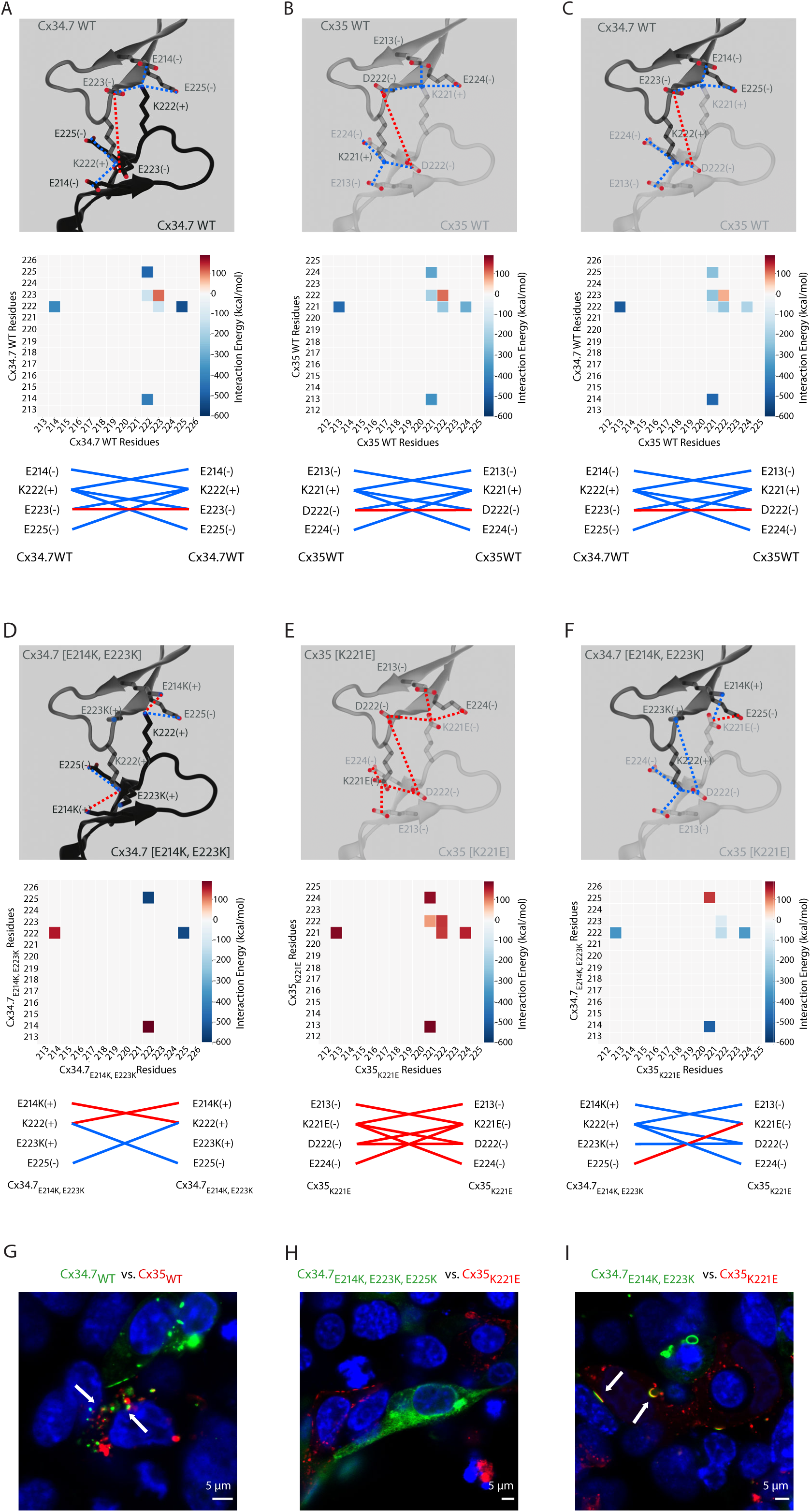
Engineering Cx34.7 and Cx35 mutants to show heterotypic, but not homotypic, hemichannel docking in mammalian cells. **A-C)** EL2-to-EL2 interactions predicted between wild type Cx34.7 and Cx35 using homology modeling. Residues predicted to form strong attractive/repulsive interactions are highlighted in blue/red dashed lines, respectively (top). Contact plots for EL2-to-EL2 interactions produced by molecular dynamics simulation (middle), and summary of interactions predicted to stabilize hemichannels pairs (bottom). Blue in the middle and bottom rows indicates residues that show attractive interactions, and red indicates residues that show repulsive interactions. The same color schemes from panels A-C apply here. Plots are shown for A) homotypic Cx34.7, B) homotypic Cx35, and C) heterotypic Cx34.7 and Cx35 interactions. **D-F)** Homology models predicting EL2-to-EL2 residue interactions for Cx34.7 and Cx35 mutant hemichannels. Plots are shown for D) homotypic Cx34.7_E214K, E223K_, E) homotypic Cx35_K221E_, and F) heterotypic Cx34.7_E214K, E223K_ and Cx35_K221E_ interactions (Cx34.7 residues are shown along the y-axis and Cx35 residues are shown along the x axis). **G-I)** Confocal images of heterotypic Cx pairs expressed in HEK 293FT cells: G) Cx34.7_WT_/Cx35_WT_, H) Cx34.7_E214K, E223K, E225K_/Cx35_K221E_, and I) Cx34.7_E214K, E223K_/Cx35_K221E_. All Cx34.7 and Cx35 proteins are expressed as mEmerald and RFP670 fusion proteins, respectively. White arrows highlight dual fluorescent labeled vesicles. Note the cytoplasmic localization of Cx34.7_E214K, E223K, E225K_ in panel H (images correspond to N = 49, 6, and 6 replicates subjected to FETCH analysis for Cx34.7_WT_/Cx35_WT_, Cx34.7_E214K, E223K, E225K_/Cx35_K221E_, and Cx34.7_E214K, E223K_/Cx35_K221E_, respectively).

Next, we introduced Cx36 into our computational model. Both wild type Cx34.7 and Cx35 showed strong interactions with Cx36, paralleling the significant FETCH scores we observed (FETCH=11.9±1.2%; T_96_ =-12.93; P=4.7×10^-23^ for Cx34.7/Cx36; FETCH=18.0±2.0%; T_96_=-18.69; P =4.7×10^-34^ for Cx35/Cx36). We then modeled the four non-docking Cx mutants identified in our initial FETCH analysis (Cx34.7_K222Q_, Cx35_E224H_, Cx35_E224K_, Cx35_E224R_) against Cx36. Though the K222Q mutation disrupted the large negative interaction energies we observed in the homotypic wild type Cx34.7 model, the three remaining negative residues within the motif that contribute to docking compatibility in Cx34.7_K222Q_ continued to show large negative interaction energies with the positive central lysine residue of Cx36, providing a potential mechanism for the heterotypic docking between Cx34.7_K222Q_ and Cx36 we observed via FETCH. On the other hand, the three candidate Cx35 mutants we tested against Cx36 using FETCH (Cx35_E224H_, Cx35_E224K_, and Cx35_E224R_) maintained the positive K221 residue that formed strong interactions with the negative residues of Cx36; however, the Cx35_E224K_, and Cx35_E224R_ mutations induced strong repulsion with the positive K238 residue of Cx36, providing insight into why these two mutants failed to heterotypically dock with Cx36 in our FETCH analyses. Additionally, introducing a smaller positively charged residue at the E224 position, as observed in the Cx35_E224H_ mutant, was sufficient to restore the interaction with Cx36 in the computational model – again mirroring the heterotypic docking profile we observed from our FETCH analyses.

Having modeled the putative interaction principles underlying the docking specificity between Cx34.7, Cx35, and Cx36 and validated our models using FETCH, we set out to design a Cx34.7/Cx35 pair that would exhibit heterotypic docking only with each other. Our strategy was to mutate residues at the four positions of our identified docking motif such that one Cx isoform contained all negatively charged interactors (Cx35), and the other all positive (Cx34.7). Our Cx35 mutant, Cx35_K221E_, showed strong repulsions in our homotypic model (Fig. 2E), did not exhibit homotypic docking upon FETCH analysis (FETCH=1.2±0.4%; T_96_=0.35; P=0.64), and failed to dock with Cx36 and Cx43 (FETCH=1.5±0.1%, T_91_=0.02, P=0.51 and 1.7±0.2, T_96_=-0.32, P=0.37 for Cx35_K221E_/Cx36 and Cx35_K221E_/Cx43, respectively). Similarly, the positively charged motif mutant, Cx34.7_E214K/E223K/E225K_, showed strong repulsions in our homotypic computation model and did not exhibit homotypic docking in FETCH analysis (FETCH=0.2±0.0%; T_96_=1.76; P=0.96). However, when we assayed Cx35_K221E_ against Cx34.7_E214K, E223K, E225K_ using FETCH, we failed to observe significant fluorescence exchange (FETCH=1.2±0.3; T_96_=0.37; P=0.64). Follow-up confocal imaging analysis of HEK 293FT cells expressing the constructs revealed that Cx34.7_E214K, E223K, E225K_ failed to properly localize to the cell membrane (compare Fig. 2G and 2H, see also Supplemental Fig. S4). Moreover Cx34.7_E214K, E223K, E225K_ predicted weak attractive interactions with Cx35_K221E_ in our homotypic gap junction computational model (Supplemental Fig. S4). Thus, we evaluated an intermediate Cx34.7 mutant protein that exhibited positively charged residues at three of the four critical interacting positions, Cx34.7_E214K/E223K_. This mutant showed repulsive interactions in our homotypic gap junction computational model (Fig. 2D), and it showed strong attractive interactions with Cx35_K221E_ (see Fig. 2F). This mutant docked with Cx35_K221E_ as confirmed via FETCH analysis and confocal microscopy (FETCH=35.7±4.1%; T_96_=-28.11; P=2.0×10^-48^; see Fig. 2I). Notably, Cx34.7_E214K/E223K_ did not show homotypic docking in our FETCH analysis (FETCH=1.1±0.2%, T_96_=0.46, P=0.68), nor did it dock with Cx36 or Cx43 (FETCH=1.0±0.2, T_96_=0.58, P=0.72 and FETCH=0.9±0.1%, T_96_=0.73, P=0.77 for Cx34.7_E214K/E223K_/Cx36 and Cx34.7_E214K/E223K_/Cx43, respectively). Curiously, the Cx34.7_E214K/E223K_ and Cx35_K221E_ mutant pair showed a higher heterotypic FETCH score than Cx36 under homotypic docking conditions (FETCH=15.2±1.1%) and the wild type Cx34.7/Cx35 pair (FETCH=12.0±0.9%), as measured using our *in vitro* assay (T_10_=4.9, P=6.4×10^-4^; T_10_=5.7, P=1.9×10^-4^ for comparisons against Cx36/Cx36 and wild type Cx34.7/Cx35, respectively, using t-test with FDR correction; N=6 replicates/group). From here on, we refer to this Cx pair, Cx34.7_E214K/E223K_ and Cx35_K221E_, as Cx34.7_M1_/Cx35_M1_ (designer Cxs version 1.0, from *Morone americana*).

### Cx34.7_M1_ and Cx35_M1_ dock heterotypically to constitute a functional electrical synapse

To determine whether our mutant Cx34.7 and Cx35 hemichannels could form a functional electrical synapse, we employed *Xenopus* oocytes as a heterologous expression system^34,35^. We also tested wild type Cx34.7 and Cx35 hemichannels as controls. Cxs were expressed in separate populations of oocytes. Oocytes expressing either two different Cxs or the same Cx, were then paired to form heterotypic and homotypic gap junctions, respectively (Fig. 3A, see methods). In our analyses of heterotypic gap junctions, we detected junctional current (*I_j_*) in paired oocytes expressing Cx34.7_M1_ and Cx35_M1_ (Fig. 3B). As expected, we also detected current in pairs expressing the wild type protein pair (Fig. 3B). In response to symmetric transjunctional voltage (*V_j_*) steps (−120mV to +120mV), the mutant heterotypic gap junction exhibited significantly lower instantaneous *I_j_* (*I_j inst_*) and steady-state *I_j_* (*I_j ss_*) currents than the wild-type heterotypic gap junction (Fig. 3C). The *I_j_* traces for both mutant and wild type pairs appeared to be asymmetric between the positive and negative *V_j_* ranges, and the *I_j_* traces recorded from one oocyte in a pair looked like a mirror image of those recorded from the other oocyte of the pair (Fig. 3B). Since these findings were indicative of a rectification property of the mutant and wild type gap junctions, we normalized the *I_j inst_* and *I_j ss_* and compared the rectification index (Fig. 3D-E). We found that *I_j ss_* was rectified in the Cx34.7 to Cx35 direction for both the mutant and wild type pairs, consistent with a prior report for the wild type proteins^14^. On the other hand, *I_j inst_*was rectified in the opposite direction for both pairs (Fig. 3E).

**Figure 3.**
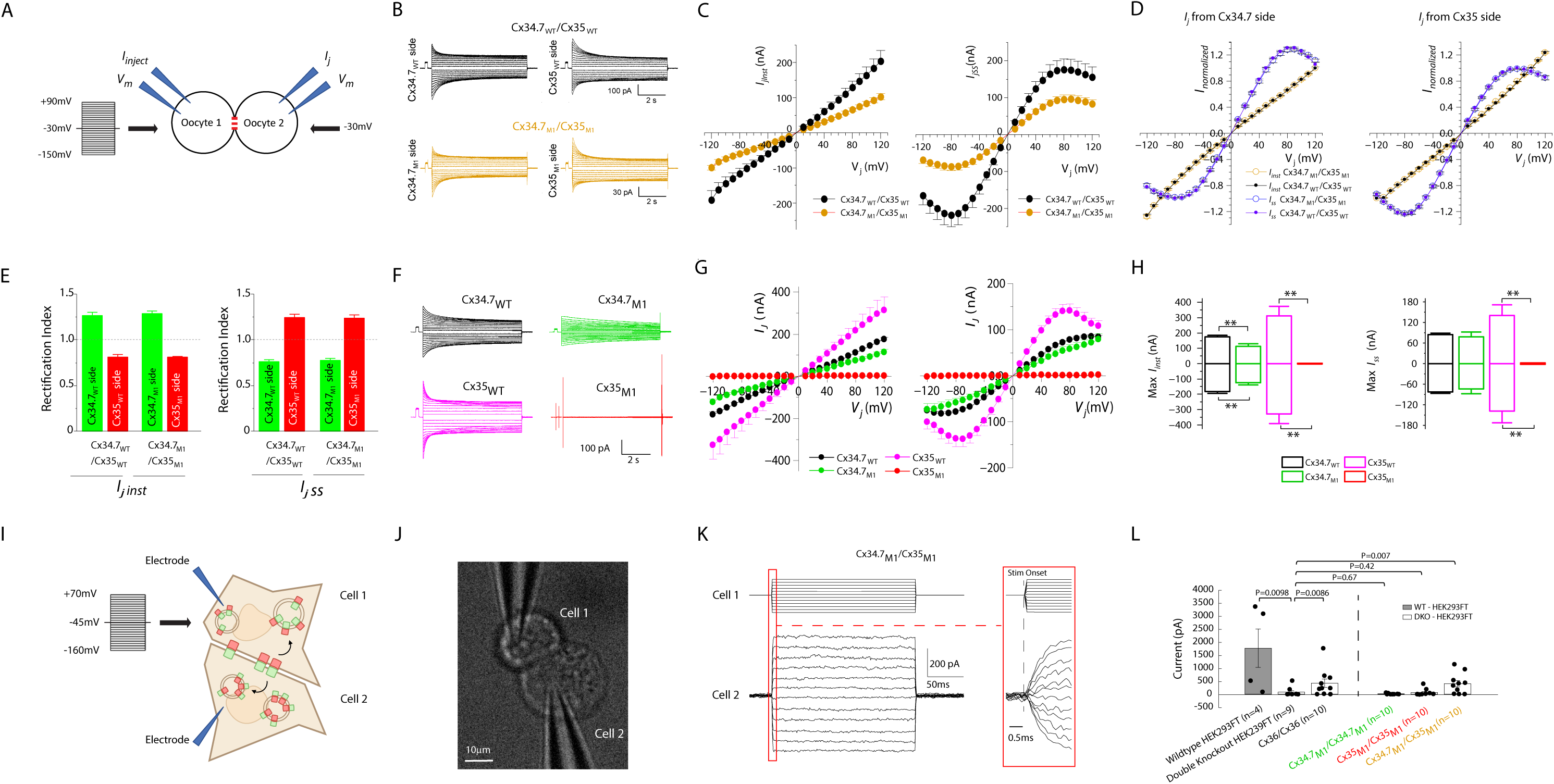
Biophysical properties of gap junctions formed by heterologous expression of wild type and mutant Cx34.7 and Cx35 in *Xenopus* oocytes. **A)** Diagram showing the characterization of connexin (Cx) gap junctions using oocyte pairs. The membrane voltage (*V_m_*) of Oocyte 1 was stepped to a series of voltages (−150 mV to +90 mV at 10-mV intervals) from a holding voltage of -30 mV, while that of Oocyte 2 was held constant at -30 mV to record junctional currents (*I_j_*). **B)** Representative *I_j_* traces from both oocytes of a pair expressing either Cx34.7_WT_ and Cx35_WT_ or Cx34.7_M1_ and Cx35_M1_. **C)** Relationships between instantaneous *I_j_* (*I_j inst_*, left) and transjunctional voltage (*V_j_*), as well as between steady-state *I_j_* (*I_j ss_*, right) and *V_j_*. Transjunctional voltage (*V_j_*) is defined as *V_m_* of Oocyte 2 – *V_m_* of Oocyte 1. **D)** Relationships between normalized *I_j inst_*and *V_j_*, as well as between steady-state *I_j ss_* and *V_j_*. **E)** Comparison of *I_j inst_*and *I_j ss_*rectification indices between Cx34.7_WT_/Cx35_WT_ and Cx34.7_M1_/Cx35_M1_ gap junctions (no significant difference was detected, unpaired *t*-test). The rectification index is defined as the ratio of the maximal *I_j_* from the negative and positive *V_j_*sides. **F)** Representative *I_j_* traces of gap junctions formed by Cx34.7_WT_, Cx34.7_M1_, Cx35_WT_, and Cx35_M1_. **G)** Relationships between *I_j inst_* and *V_j_*, as well as between *I_j ss_* and *V_j_*. **H)** Comparison of the maximal *I_j inst_* and *I_j ss_* at positive and negative *V_j_* among the different gap junctions. **I)** Diagram showing functional characterization of Cx34.7_M1_ and Cx35_M1_ using HEK293FT pairs. **J)** Image showing cell pair characterized in Fig. 3I. **K)** Representative injected and recorded current traces (top and bottom). Current traces are depicted at a 20mV step 1ms resolution, and the stimulation onset is shown to the right at 0.1ms resolution. **L)** Current at the maximum transjunctional voltage is shown for wild type and Cx double knockout HEK293FT cells, and cell pairs transfected with Cx36, Cx34.7_M1_, or Cx35_M1_ under conditions of homotypic and heterotypic pairing.

In our analyses of homotypic gap junctions, we detected *I_j_*from oocytes expressing Cx34.7_WT_, Cx35_WT_, and Cx34.7_M1_, but not Cx35_M1_ (Fig. 3F). The homotypic gap junctions differed in *I_j_* amplitude (Fig. 3G-H), the relationship between the steady-state junctional conductance (*G_ss_*), and the *V_j_*, and the deactivation rate (see Supplemental Fig. S5). Overall, these findings confirmed the formation of functional Cx34.7_M1_/Cx35_M1_ gap junctions and the disruption of Cx35_M1_ homotypic docking. These observations were consistent with our initial screening analysis using FETCH. Contradictory results were observed for Cx34_M1_ under homotypic conditions, where we observed functional gap junctions using *Xenopus* oocytes but disrupted docking using FETCH in mammalian cells.

To address these conflicting findings observed for Cx34_M1_ under homotypic conditions, we next tested whether Cx34_M1_ could form a functional homotypic electrical synapse in HEK293FT cells (Fig. 3I). Here, we used CRISPR-Cas9 to generate a new HEK293FT cell line for which the endogenous Cx43 and Cx45 proteins were disrupted (Cx double knockout HEK293FT cells, see Supplemental Fig. S6 and methods). This approach diminishes the electrical synapses that naturally form between HEK293FT cells^36^, enabling us to assess whether functional connectivity is restored by Cx34.7_M1_ expression. While we found that heterotypic expression of Cx34.7_M1_ and Cx35_M1_ increased electrical connectivity between pairs of Cx double knockout HEK293FT cells (U=60 and P=0.0066 compared to non-transfected Cx double knockout cell pairs using one-tailed Wilcoxon rank sum test followed by FDR correction, Fig. 3J-L; see also Supplemental Figure S7), neither homotypic expression of Cx34.7_M1_ or Cx35_M1_ had this impact (U=95 and P=0.67; U=87 and P=0.42 for Cx34.7_M1_ and Cx35_M1_, respectively, compared to non-transfected Cx double knockout cell using one-tailed Wilcoxon rank sum test followed by FDR correction; see Fig. 3L). Thus, Cx34.7_M1_ failed to show homotypic docking capacity in the mammalian cell line. Thus, in addition to our finding that Cx34.7M1 failed to dock in a homotypic configuration in our FETCH analysis, we found that Cx34.7_M1_ also failed to form a functional homotypic electrical synapse when expressed in a mammalian cell line.

### Cx34.7_M1_/Cx35_M1_ electrical synapse regulates circuit function and behavior in *C. elegans*

Next, we set out to determine the *in vivo* docking selectivity and functionality of our Cx34.7_M1_ and Cx35_M1_ hemichannels by testing whether distinct hemichannel pairs could regulate the activity of two neurons that compose a circuit and their output behavior. Specifically, we evaluated hemichannels under homotypic and heterotypic conditions against the other major Cx proteins expressed in the mammalian CNS (Cx36 and Cx43). We anticipated that these experiments would further clarify whether the inconsistent homotypic interaction observed for Cx34.7_M1_ hemichannels between our *Xenopus* oocyte and HEK293 FETCH experiments could potentially limit their application for precision circuit editing in mammals. Additionally, we tested our Cx34.7_M1_ and Cx35_M1_ channels under heterotypic conditions to confirm their *in vivo* functionality.

Here, we capitalized on *C. elegans* since multiple groups had established that selectively expressing Cx36 is sufficient to reconstitute a functional electrical synapse between two connected neurons. The presence and function of this resultant Cx36/Cx36 synapse has been confirmed via microscopy^7^, measurements of synaptic physiology^7^, calcium imaging^9^, and behavior^6–11^. Thus, we assessed whether we could induce changes in calcium imaging and behavior with Cx34.7_M1_/Cx35_M1_ in a manner that mirrored Cx36/Cx36.

*C. elegans* do not have an innate temperature preference and can thrive in a broad range of temperatures^37^. However, *C. elegans* trained at a particular temperature in the presence of food will migrate towards that temperature when they are subsequently placed on a temperature gradient^37^. This learned preference is in part mediated by plasticity of the synapse between a thermosensory neuron (AFD, presynaptic) and an interneuron (AIY, postsynaptic)^38^. Critically, plasticity in AFD can be genetically manipulated to affect transmission to AIY and to predictably encode the behavioral preference that must otherwise be learned^9^.

We have previously shown that heterologous expression of Cx36 could be used to edit this circuit, via bypassing the presynaptic plasticity mechanisms between AFD and AIY that contribute to the learned temperature preference^9^. As such, circuit-edited worms show a persistent preference for warmer temperatures (Fig. 4A). We therefore used this circuit to validate the functionality of our engineered gap junction proteins (as assessed by calcium imaging and quantitative behavior testing).

**Figure 4.**
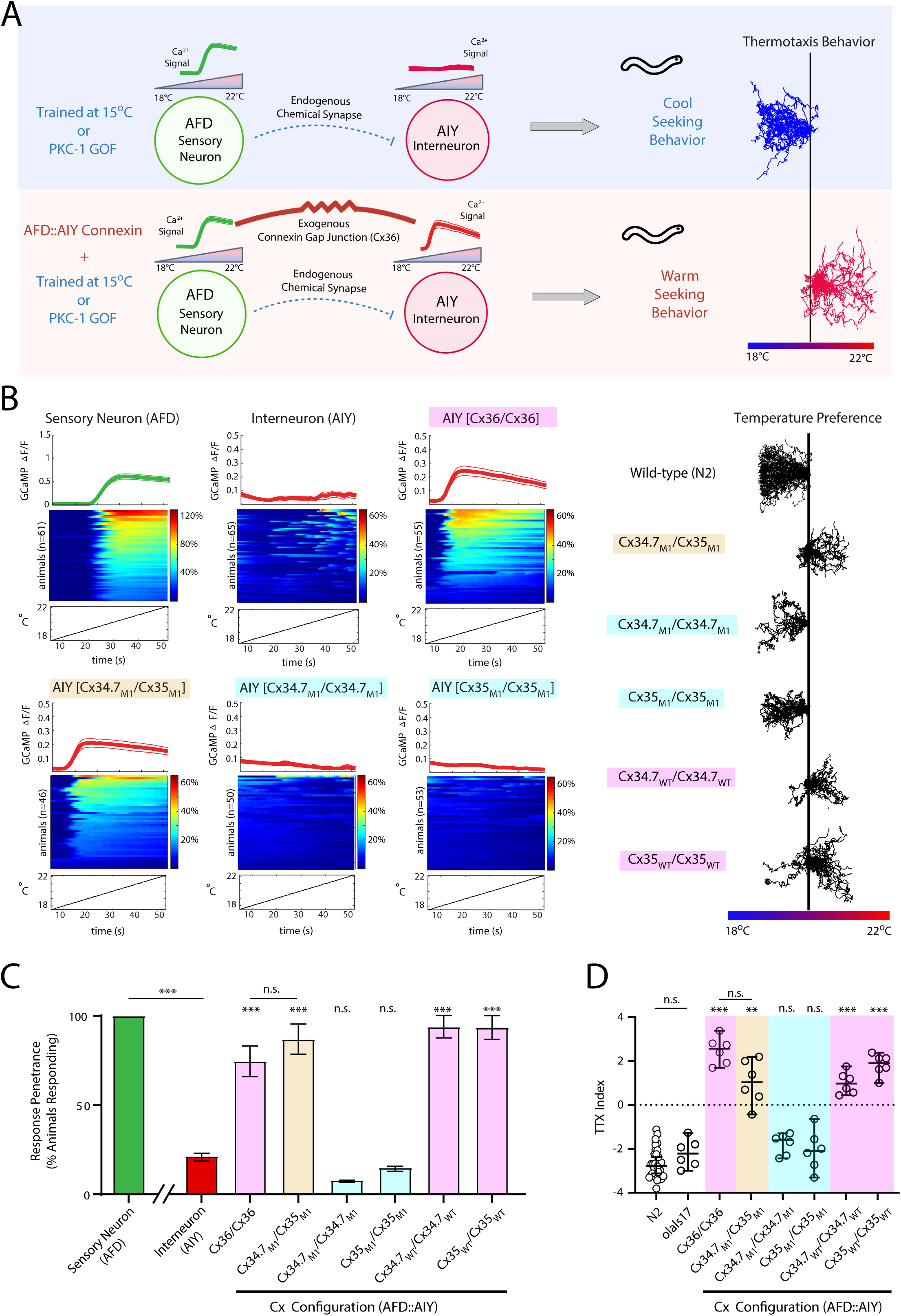
Heterologous connexin hemichannels couple *C. elegans* neurons and recode thermal preference. **A)** Schematic of the AFD◊AIY synaptic communication and expressed temperature preference. The AFD thermosensory neuron has a robust calcium response to warming stimuli. *C. elegans* raised in the presence of food at 15°C, or animals with a Protein Kinase C*-1* gain-of-function (PKC-1 GOF) mutation, move towards cooler temperatures when placed on a thermal gradient from 18-22° (top). Heterologous expression of Cx36 hemichannels between AFD and AIY results in synchronization of the signal to AIY and promotes warm-seeking behavior (bottom). **B)** Calcium traces of neurons expressing heterologous connexin (Cx) hemichannel pairs (left). Baseline AFD and AIY responses are also shown. Each panel depicts the average group trace (top, data shown as mean±SEM), heatmaps of individual animals (middle), and the temperature stimulus (bottom). Behavioral traces for each group are shown on the right. Traces are shown for C. *elegans* homotypically expressing wild type Cx hemichannels (pink highlight), heterotypically expressing the mutant pair (tan highlight), and homotypically expressing mutant Cx hemichannels (cyan highlight). **C)** Portion of animals showing neuronal calcium responses based on the traces shown in B; ***p<0.0005 using Fisher’s exact test for penetrance. Error bars denote 95% C.I. **D)** Thermotaxis indices corresponding to experimental groups. Each data point represents the thermotaxis preference index of a separate assay (12-15 animals/assay), with the median for each group plotted denoted by a black horizontal. **p<0.005; ***p<0.0005 vs. N2; Error bars denote 95% C.I.

We first expressed Cx34.7_M1_ in AFD cells and expressed Cx35_M1_ in AIY cells (see Supplemental Fig. S8 and Supplemental Table S2). Similar to Cx36/Cx36, expression of Cx34.7_M1_/Cx35_M1_ in the AFD/AIY pair resulted in functional coupling between AFD and AIY, as assessed via calcium imaging (Fig. 4B left and 4C; P<0.0005 using Fisher exact test with an FDR correction). These *C. elegans* constitutively migrated towards warmer temperatures when placed on a thermal gradient, mirroring the animals expressing heterologous Cx36/Cx36 (F_7,17.91_= 84.99; P<0.0001 using Welch one-way ANOVA followed by Dunnett’s T3 multiple comparisons; p<0.005 vs. the N2 animals; Fig. 4B right and 4D). Expression of Cx34.7_WT_ or Cx35_WT_, but not Cx34.7_M1_ or Cx35_M1_, in both AFD and AIY neurons synchronized the two cells and modulated behavior (Fig. 4B-D; P<0.0005). We also evaluated Cx34.7_M1_ and Cx35_M1_ hemichannels against Cx36 and Cx43. *C. elegans* expressing Cx34.7_M1_/Cx36, Cx34.7_M1_/Cx43, Cx36/Cx35_M1_ or Cx43/Cx35_M1_ in AFD/AIY pairs all continued to migrate towards cold temperatures (F_7,10.67_= 19.29; P<0.0001 using Welch one-way ANOVA followed by Dunnett’s T3 multiple comparisons; p<0.001; see Supplemental Fig. S9).

Taken together, these findings confirmed that our Cx34.7 _M1_/Cx35_M1_ electrical synapse modified *C. elegans* behavior and physiology in a manner that was statistically indistinguishable from the Cx36/Cx36 electrical synapse. Our findings also supported the docking properties we predicted for the mutants using our *in vitro* screen and *in silico* studies, since both Cx34.7_M1_ and Cx35_M1_ failed to alter behavior/physiology when expressed in homotypic configurations or in heterotypic configurations against Cx36 and Cx43.

### Cx34.7_M1_/Cx35_M1_ electrical synapse enhances synchrony within a mouse neural circuit

Having established the *in vivo* docking selectivity and functionality of our Cx34.7_M1_/Cx35 _M1_ pair, we set out to determine whether these proteins could modulate mesoscale neural circuitry in mammals. After verifying their expression and trafficking (see Supplemental Fig. S10), we chose to edit a circuit composed of two distinct cell types. Mice are an ideal species in which to test cell-type specificity, because they are highly amenable to cell-type specific access via selective promoters and Cre-recombinase targeting. Excitatory pyramidal neurons (PYR) and parvalbumin expressing fast-spiking (PV+) interneurons can form microcircuits whereby PYR neurons excite PV+ interneurons, which in turn inhibit PYR neurons (Fig. 5A). This PYR↔PV+ neural circuit has been well-characterized in the hippocampus, where medial prefrontal PYR neurons show activity coupled to the phase of hippocampal theta frequency (4-10Hz) oscillations during spatial exploration^39^ and PV+ interneuron activity is coupled to hippocampal gamma frequency oscillations (30-80Hz)^40^. Notably, the activity of this PYR–PV+ microcircuit is reflected in the millisecond-resolved synchrony between the phase of theta oscillations and the amplitude of gamma oscillations in rodents^41^.

**Figure 5.**
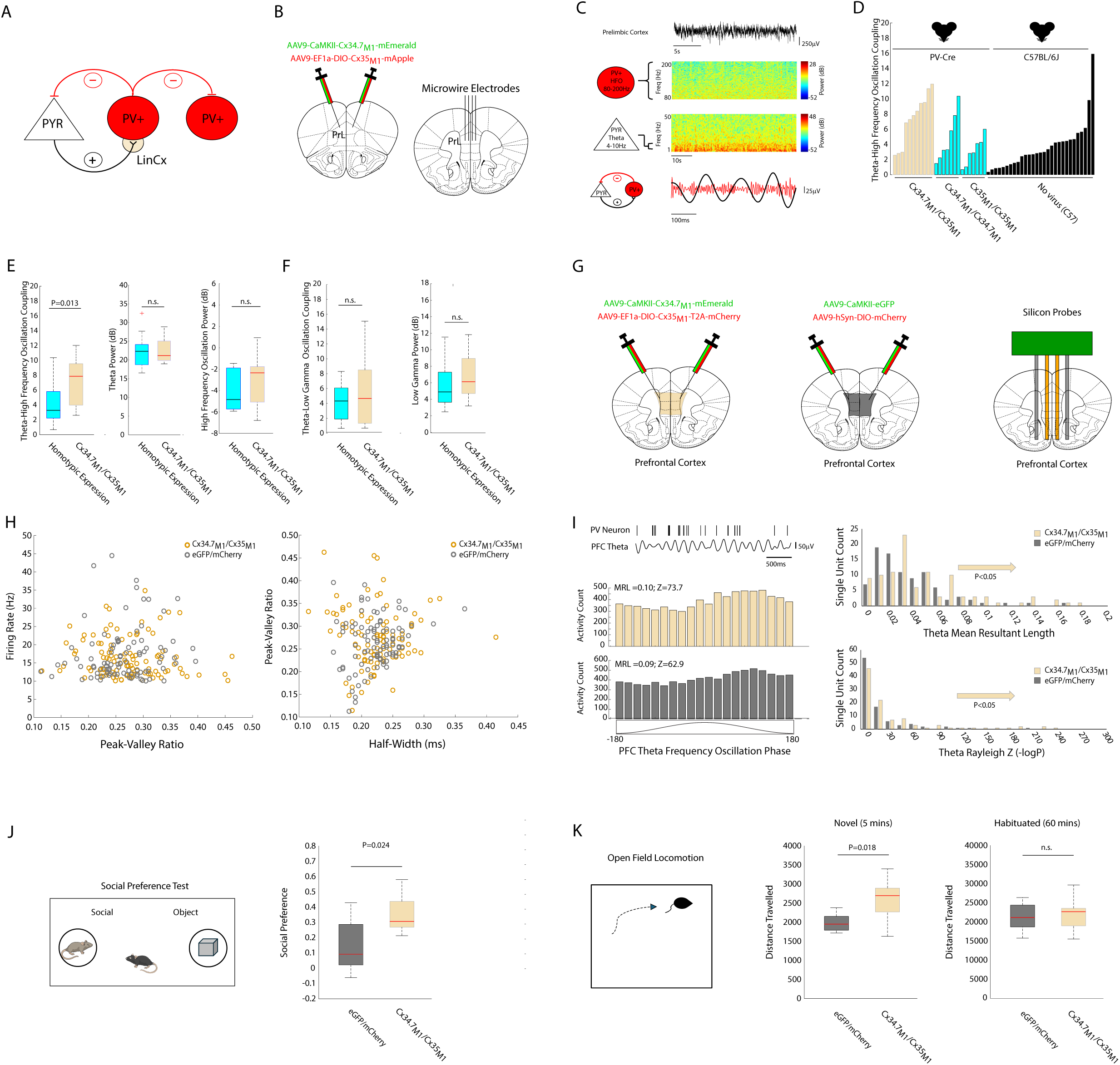
LinCx edits microcircuit dynamics at the millisecond timescale in mice. **A)** Prefrontal cortex microcircuit comprising an excitatory pyramidal neuron (PYR) and 2 parvalbumin expressing fast-spiking (PV+) interneurons. The tan circle highlights the PV+ target for synaptic editing. **B)** PV-Cre mice were bilaterally co-injected with AAV-CaMKII-Cx34.7_M1_ (green) and AAV-DIO-Cx35_M1_ (red). Control mice were injected with a Cx34.7_M1_ or Cx35_M1_ pair of viruses that expressed the same hemichannel in both cell types. Mice were subsequently implanted with microwires in prefrontal cortex. **C)** Representative local field potential recorded from prefrontal cortex (top). Power spectrograms show theta (4-10Hz) and high frequency oscillation activity (80-200Hz), which corresponds to PYR and PV+ interneuron firing, respectively (middle). Microcircuit function is represented by the coupling between the phase of theta oscillations (black) and the amplitude of high frequency oscillations (red, bottom). **D)** Distribution of theta–high frequency coupling scores (Modulation index, z-score) observed across un-injected C57BL/6j mice (N=29, black bars), PV-Cre mice co-injected with a Cx34.7_M1_ (N=7) or Cx35_M1_ (N=7) pair of viruses (light blue bars), or mice co-injected with a docking Cx34.7_M1_/Cx35_M1_ pair (N=11, tan bars). **E)** Mice co-injected with the Cx34.7_M1_/Cx35_M1_ pair showed significantly higher theta-high frequency oscillation coupling (left), but not theta power or high frequency oscillation power (middle, right), compared to mice expressing Cx34.7_M1_ and Cx35_M1_ under homotypic conditions expressing. **F)** Circuit editing had no impact on theta-low gamma oscillatory coupling (left) or low gamma power (right) for these mice. **G)** PV-Cre mice were bilaterally co-injected with AAV-CaMKII-Cx34.7_M1_ and AAV-DIO-Cx35_M1_. Control mice were injected with a Cx34.7_M1_ or Cx35_M1_ pair of viruses which expressed the same hemichannel in both cell types. Mice were subsequently implanted with silicon probes in medial prefrontal cortex. **H)** Waveform properties of PV+ interneurons recorded from experimental (tan circles) and control (dark grey circles) mice. **I)** Activity of PV+ interneuron and theta oscillations recorded concurrently from the same channel (top left). Theta oscillation phase firing distributions of a PV+ interneuron recorded from an experimental mouse (middle left) and a control mouse (bottom left). Distribution of mean resultant lengths (MRLs) across the population of PV+ interneurons recorded from experimental and control mice (top right). Distribution of Rayleigh test statistics across the population of PV+ interneurons recorded from experiment and control mice (bottom right), where Z = -log P. **J)** Social preference assay used to assess impact of PYR and PV+ circuit editing on social behavior (left). LinCx-edited mice exhibited an increase in social preference relative to control mice (right). **K)** LinCx-edited mice exhibited higher exploratory drive in a novel environment compared to controls (left, middle). No group difference in gross locomotor behavior was observed following habituation (right). For box and whisker plots, the central mark is the median, the edges of the box are the 25th and 75th percentiles, the whiskers extend to the most extreme datapoints the algorithm considers to be not outliers, and the outliers are plotted individually as a “+”.

PYR–PV+ microcircuits are also observed in the prefrontal cortex with slightly different neurophysiological properties^42^. Like the cellular dynamics observed in the hippocampus, prefrontal cortex PYR neurons phase-couple to locally recorded theta oscillations^43^. On the other hand, within prefrontal cortex, PV+ interneurons best couple to the phase and amplitude of local high frequency oscillations (HFO, 80-200Hz)^44^. Thus, to determine the effect of our electrical synapse, we quantified the coupling between the phase of prefrontal cortex theta oscillations and the amplitude of prefrontal cortex HFO as a proxy for prefrontal cortex PYR–PV+ microcircuit activity. Specifically, we expressed our Cx34.7_M1_/Cx35_M1_ synapse at the PYR–PV+ interface, hypothesizing that this manipulation would enhance the millisecond-timed coupling between theta and HFO in the prefrontal cortex.

We developed an AAV virus (AAV9-Ef1α-DIO-Cx35_M1_) to selectively target Cx35_M1_ to cells expressing Cre-recombinase, and another virus (AAV9-CaMKII-Cx34.7_M1_) to express Cx34.7_M1_ non-selectively across all neurons. We then co-injected PV-Cre mice with both viruses bilaterally in the prelimbic cortex (N=11; Fig. 5A-B). Two groups of PV-Cre control mice were injected with viruses to express Cx34.7_M1_ or Cx35_M1_ non-selectively across all neurons (N=7 per group, see methods). Finally, we also tested a third group of control C57BL/6J mice that had not been injected with virus (N=29). Neural oscillatory activity was recorded from prelimbic cortex while mice explored an open field (see Supplemental Fig. S11A).

To determine the coupling between theta (4-10Hz) and high frequency oscillations (80-200Hz), we isolated local field potential activity in these two frequency bands (Fig. 5C). We then determined their phase–amplitude coupling relationships using the established Modulation index (z-score), which quantifies the statistical likelihood that measured relationships between two oscillations would be observed by chance (see methods)^45^. Using this approach, we found significant theta–HFO coupling from the majority of implanted mice (80%, 43/54; Fig. 5D). Moreover, theta–HFO coupling was significantly higher in the mice expressing our electrical synapse, compared to the pooled group of control mice expressing the channels under homotypic configurations (U=141; P<0.013 using one-tailed rank-sum test; Fig. 5E). Thus, expression of the synapse was sufficient to enhance millisecond-timed coupling within a circuit defined by two precise cell types in mammals. Interestingly, our post-hoc analysis found no differences in theta–HFO coupling between mice expressing Cx34.7_M1_/Cx34.7_M1_ or Cx35_M1_/Cx35_M1_ in the homotypic configuration compared to uninfected C57BL/6J control mice (U=509 and 542; P=0.28 and 0.84 using two tailed rank-sum test, for Cx34.7_M1_/Cx34.7_M1_ or Cx35_M1_/Cx35_M1_, respectively, see Fig. 5D). These findings support the heterotypic selectivity of the two mutant proteins *in vivo*.

Importantly, we found no differences in theta or HFO power between mice expressing the synapse and the control mice expressing mutant Cxs in homotypic configurations across the PYR–PV+ circuit (U= 172 and 167; P=0.60 and 0.43 using two tailed rank-sum test, for theta and HFO power, respectively; Fig. 5E). Similarly, no difference in theta–low gamma oscillation cross-frequency phase coupling was observed across these groups (U= 176, P=0.76 using two tailed rank-sum test; Fig. 5F). Thus, our electrical synapse selectively increased the synchrony between theta and high frequency oscillation activity in the medial prefrontal cortex.

Next, we tested whether this increased synchrony could be observed at the level of single neurons. Three new experimental PV-Cre mice were infected with AAV9-hsyn-DIO-Cx35_M1_-T2A-mCherry to selectively target Cx35_M1_ to cells expressing Cre-recombinase, and the non-selective AAV9-CaMKII-Cx34.7_M1_-mEmerald virus. Three control mice were infected with AAV9-hsyn-DIO-mCherry and AAV9-CaMKII-eGFP. Two weeks later, these mice were implanted with high density silicon recording probes (Fig. 5G and Supplemental Fig. S11C). Following recovery, neural activity was recorded for 10 minutes while mice were in their home cage. PV+ interneurons were identified based on previously validated waveform criteria (e.g., peak to valley ratio < 1.1 and mean firing rate >10Hz)^46^. Notably, no differences in waveform properties were observed between the experimental and control group (t_190_=1.07 and P=0.29 for peak valley ratio; t_190_=0.15 and P=0.88 for half-width using unpaired two tailed t-tests; N = 91 and 101 total medial prefrontal cortex PV+ single neurons for the experimental and control groups, respectively; Fig. 5H).

After establishing that the activity of prefrontal cortex PV+ interneurons was better coupled to the phase of HFOs than gamma oscillations (see Supplemental Fig. S12), we compared the activity profiles of PV+ interneurons across the two groups. We observed no difference in the mean firing rate of PV+ single units between the experimental and control groups (t_190_=0.90 and P=0.37 for comparison of firing rate using unpaired two tailed t-test; Fig 5H), consistent with our finding that the electrical synapse did not impact the amplitude of HFO activity in the prefrontal cortex (see Fig 5E, right). Next, we quantified the coupling of each PV+ interneuron’s activity to local theta oscillations by determining its mean resultant length (MRL) and with the Rayleigh test of circular uniformity (Fig. 5I). In contrast to the prior study^44^, we found that most PV+ interneurons showed phase coupling to theta oscillations in both groups (69/91 and 83/101 neurons for experimental and control mice, respectively). Strikingly, when we compared activity across the groups, we found that the experimental mice showed stronger PV+ phase coupling to theta oscillations than the control group (t_190_=2.34 and P=0.01 for MRL; t_190_=2.31 and P=0.01 for Rayleigh Z using unpaired one tailed t-tests; Fig. 5I). Thus, the engineered electrical synapse increased coupling of PV+ interneurons to theta oscillations, again consistent with our observations for the coupling of HFO activity.

Finally, prefrontal cortical microcircuit dysfunction has been implicated in mediating social deficits in autism. Thus, we tested whether increasing coupling within this circuit would enhance social behavior. LinCx-edited mice exhibited higher preference for the social stimulus in a social preference task (t_13_=2.55, P=0.024 for LinCx-edited vs. control mice using unpaired t-test followed by false-discovery rate correction, N=7-8 per group; Fig. 5J). Interestingly, LinCx-edited mice also exhibited higher exploratory drive when placed in a novel open field (t_13_=2.69, P=0.018 for LinCx-edited vs. control mice using unpaired t-test; see also Fig. 5K, left). No differences in gross locomotor behavior were observed when animals were previously habituated to the testing arena (t_13_=0.37, P=0.72 for LinCx-edited vs. control mice using unpaired t-test; see also Fig. 5K, right). Overall, these findings showed that expression of the LinCx electrical synapse causally enhanced coupling of single unit activity within a local mammalian circuit and modified behavior accordingly.

### Cx34.7_M1_/Cx35_M1_ synapse potentiates a multi-regional circuit in mice

Having established that our electrical synapse could enhance electrical coupling within a local microcircuit, we next tested whether it could potentiate a long-range circuit consisting of cells in two different brain regions. The infralimbic cortex (IL, another anatomical subdivision of mouse medial prefrontal cortex) sends a monosynaptic projection to medial dorsal thalamus (MD) in mice. We selected this circuit to test the functionality of Cx34.7_M1_/ Cx35_M1_, given our prior experience in quantifying its physiological properties and role in stress behavior ^47,48^. Specifically, the tail suspension test is a classic assay that measures the behavioral response of mice to an inescapable negative experience in which they are suspended upside down by their tail^49^. Exposure to stress diminishes behavioral responses during the assay^50^, and the assay induces a robust stress response^51^. As such, repeated exposure to the tail suspension test increases immobility during subsequent testing^48^ (Fig. 6A, top). This behavioral adaptation is specific to the stress context, as decreased mobility is not observed when locomotor behavior is quantified in an open field immediately prior to each tail suspension stress session^48^.

**Figure 6.**
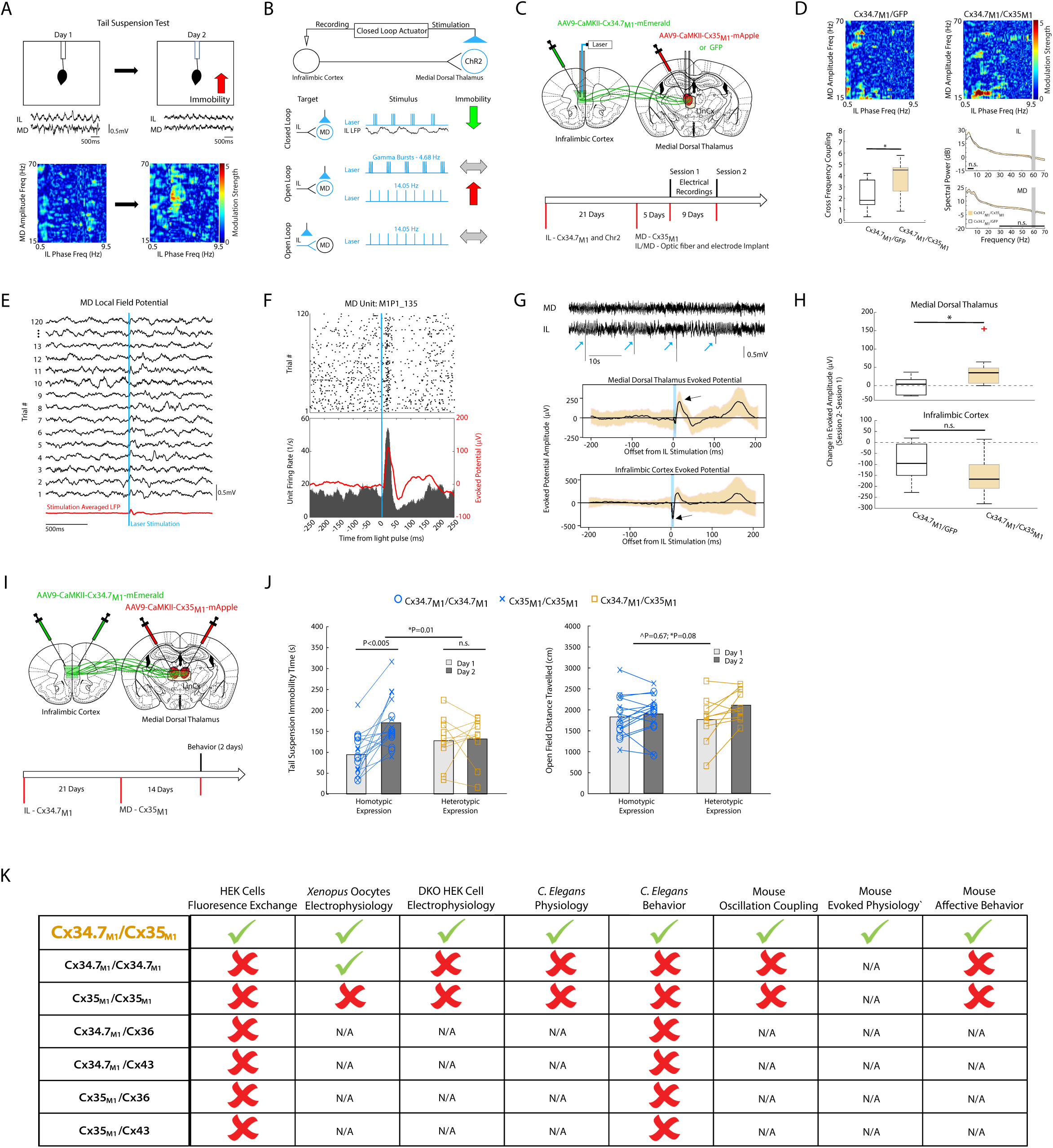
LinCx edits microcircuit dynamics at the millisecond timescale in mice. **A)** The behavioral and neurophysiological impacts of repeated tail suspension stress (data from ^48^). Plots on the bottom show increased in coupling between infralimbic cortex (IL) 2-7Hz oscillations and medial dorsal thalamus (MD) 30-70Hz oscillations due to repeat tail suspension test exposure. **B**) Schematic showing a closed-loop optogenetic approach to synchronizing neuronal firing in MD to IL oscillations as performed in our previous work^48^ (top). Next row depicts behavioral impact of causally coupling MD activity to IL 3-7Hz oscillations using closed-loop optogenetic stimulation. Middle and bottom rows depict behavioral outcomes of stimulating MD (middle row) or IL terminals (bottom row) in MD with a pattern that was uncoupled to ongoing IL activity (open loop). **C)** Schematic of viral and optical fiber/electrode targeting approach (top), and experimental timeline for optogenetic interrogation of the IL◊MD circuit (bottom). D) Representative plots showing coupling between IL and MD oscillations for Cx34.7_M1_/GFP controls mice (top left) and Cx34.7_M1_/Cx35_M1_ (top right). Mice co-injected with the Cx34.7_M1_/Cx35_M1_ pair showed significantly higher coupling between IL 3-7Hz and MD 30-70Hz oscillations (bottom left), but there were no group differences in IL 3-7Hz or MD 30-70Hz oscillatory power (bottom right, data shown as mean±s.e.m; note that the distributions overlap). **E)** Response of representative LFP oscillations from MD in response to an optogenetic light pulse in IL (light blue vertical line). LFP activity averaged across 120 light pulses (MD evoked potential) is shown below in red (light stimulation was delivered at 1mW, 10ms pulse width). **F)** Response of MD neuron to an optogenetic light pulse in IL (light blue vertical line). Cellular activity across 120 light pulses is shown below, with the evoked potential recorded from the same channel overlaid in red. The positive deflection in the evoked potential reflects firing of the neuron. **G)** (top) Representative LFP oscillations recorded from IL and MD during light stimulation. Note the four highlighted supraphysiological IL responses induced by light stimulation (light blue arrows). Representative mean light evoked potential recorded concurrently from an MD (middle) and IL microwire (bottom). Note the large negative instantaneous deflection in the IL channel, and the positive deflection the MD channel (highlighted by arrows). Data shown as mean ± std across ∼30 light pulses. **H)** Box and whisker plots show change in amplitude of evoked potential across sessions in MD (top) and IL (bottom) in mice expressing the Cx34.7_M1_/Cx35_M1_ vs. control mice; *P<0.05 using one tailed t-test. **I)** Schematic of viral injection strategy and experimental timeline for quantifying impact of IL◊MD LinCx editing on behavior. **J)** Immobility time and distance travelled during repeat tail suspension (left) and repeat open field testing (right). Mice co-injected with Cx34.7_M1_ (N=8) or Cx35_M1_ (N=8) in homotypic non-docking configurations showed stress-induced behavioral adaptation during repeat tail suspension testing (blue), while mice co-injected with the functional Cx34.7_M1_/Cx35_M1_ pair did not (N=10; tan). No significant behavioral differences were observed between viral groups in the open field test; ^ denotes Group effect; * denotes Group × Day interaction effect using mixed-effects model ANOVA. For box and whisker plots, the central mark is the median, the edges of the box are the 25th and 75th percentiles, the whiskers extend to the most extreme datapoints the algorithm considers to be not outliers, and the outliers are plotted individually as a “+”. **K)** Summary of biophysical and electrophysiological properties of Cx34.7_M1_ and Cx35_M1_.

In our prior work, we found that exposure to the tail suspension test induced coupling between low frequency oscillations in the IL and low-gamma oscillations in the MD^48^ (see Fig. 6A). Furthermore, when we exogenously recapitulated coupling between IL and MD using a brain-machine interface, mice showed reduced behavioral adaptation in the assay^48^. Stimulation of MD in a manner that was uncoupled to IL activity failed to yield this outcome^48^ (see Fig. 6B for summary). Together, these findings established the IL→MD circuit’s role in stress compensatory behavior^48^.

Here, we co-injected BALB/cJ mice with AAV9-CaMKII-Cx34.7_M1_ and AAV9-CamKII-Chr2 in left IL. Three weeks later, we injected these mice with AAV9-CaMKII-Cx35_M1_ in left MD and implanted microwire recording electrodes in IL and MD (Fig. 6C). After another 5 days of surgical recovery, we recorded baseline LFP activity and activity in response to 10ms pulses of optogenetic stimulation to IL. This timeline ensured expression of Chr2 and Cx34.7_M1_ in IL, but minimal trafficking of Cx34.7_M1_ to the IL axonal terminals in MD (see Supplemental Fig. S10), and minimal local expression of Cx35_M1_ in MD. We acquired additional recording and stimulation data nine days later (Session 2: 5 weeks after the initial IL injection/ 2 weeks after the MD injection), enabling strong trafficking of Cx34.7_M1_ to IL and substantial local Cx35_M1_ expression. A control group was infected with AAV9-CaMKII-GFP in MD instead of Cx35_M1_.

We first compared coupling across the IL→MD circuit between the two groups, two weeks after the MD injection (i.e., second baseline recording session). Mice injected with Cx35_M1_ showed stronger coupling between IL 2-7Hz oscillations and MD 30-70Hz oscillations, compared to the mice injected with GFP (U=88; P=0.033 using one-tailed rank-sum test; Fig. 6D, top and bottom left). No group differences were observed in MD 30-70Hz power (U= 80; P= 0.19 using two tailed rank-sum test; see Fig. 6D, bottom right) or in IL 2-7Hz power (U= 81; P=0.16 using two tailed rank-sum test). Thus, expression of Cx34.7_M1_/ Cx35_M1_ enhanced oscillatory coupling across the IL→MD circuit as it had for the prelimbic cortex PYR– PV+ microcircuit (Fig. 5).

Next, we directly interrogated the IL→MD circuit in the mice expressing Cx34.7_M1_/ Cx35_M1_. In our prior study, we observed a positive evoked potential in MD within 25ms of IL stimulation^47^. We first confirmed that this evoked response directly reflects the activation of single units within MD (see Fig. 6E-F and Supplemental Fig. S13), establishing its local relevance. During our first recording session, we again observed a positive evoked potential in MD within 25ms of IL stimulation with 1mW of blue light (Fig. 6G, see also Supplemental Fig. S14). When we repeated our stimulation experiment nine days later, almost all the mice expressing Cx35_M1_ showed an increase in the amplitude of their evoked MD activity (N=8/9 mice, 41±16mV). This increase was significantly higher than what we observed from the control group across sessions (N=6, 1±10mV; t_13_=1.9, P= 0.043 using one tailed t-test; see Fig. 6H, top). There was no statistical difference between the change in the evoked IL response across groups (−154±30mV and -93±38mV for the Cx35M1 and GFP groups, respectively; t_13_=-1.3, P= 0.11 using one tailed t-test; see Fig. 6H, bottom). Taken together, these findings provided causal evidence that expression of our synapse potentiated the IL→MD circuit.

### Paired Cx34.7_M1_ and Cx35_M1_ hemichannels in a long-range circuit modifies behavior in mice

Finally, we set out to determine whether expressing our engineered electrical synapses across a long-range circuit could modify behavior. Since we had previously shown that exogenous stimulation of the IL→MD circuit using closed-loop optogenetic stimulation enhances stress compensation^48^, we hypothesized that expression of the Cx34.7_M1_/Cx35_M1_ electrical synapse across the IL→MD circuit would also enhance stress compensation and diminish the stress adaptation observed between the two sessions of the tail suspension test (i.e., increased immobilization). Thus, we injected mice with AAV9-CaMKII-Cx34.7_M1_ bilaterally into IL, followed by a second injection of AAV9-CaMKII-Cx35_M1_ bilaterally in MD, three weeks later (Fig. 6I; N=10 mice; see Supplemental Fig. S11B). A negative control group of mice was injected with either AAV9-CaMKII-Cx34.7_M1_ (N=8) or AAV9-CaMKII-Cx35_M1_ (N=8) in both regions. All mice were subjected to two days of testing in an open field and with tail suspension after two weeks of recovery.

Mice expressing the Cx34.7_M1_/Cx35_M1_ hemichannel pair across the IL→MD circuit did not show behavioral adaptation in response to repeat tail suspension testing (F_1,24_=7.85, P=0.01 for Group × Day interaction effect using mixed-effects model ANOVA; t_9_=0.19; P=0.85 for post-hoc testing using two-tailed paired t-test for Cx34.7_M1_/Cx35_M1_ mice across days; Fig. 6J, left). Increases in immobility were observed in the negative control group (t_18_=4.9; P=1.7×10^-4^ for post-hoc testing using two-tailed paired t-test for pooled group of Cx34.7_M1_/Cx34.7_M1_ and Cx35_M1_/Cx35_M1_ mice across days), and post-hoc analysis revealed increases in immobility in both the Cx34.7_M1_/Cx34.7_M1_ and Cx35_M1_/Cx35_M1_ control groups expressing the mutant hemichannels in non-docking configurations, independently (t_7_=3.3; P=0.01, and t_7_=5.5; P=9.5×10^-4^, respectively using two-tailed paired t-test). Moreover, these control mice showed increases in tail suspension immobility that were statistically indistinguishable from that observed in uninfected BALB/cJ mice (Supplemental Fig. S15). No differences in open field exploration were observed across groups between the mice that expressed Cx34.7_M1_/Cx35_M1_ and control mice expressing the hemichannels in non-docking configurations (F_1,24_=0.19, P=0.67 for Group effect; F_1,24_=7.69, P=0.01 for day effect; F_1,24_=3.4, P=0.08 for Group × Day interaction effect using mixed effects model ANOVA) (Fig. 6J, right). Thus, expression of Cx34.7_M1_/Cx35_M1_ in a long-range circuit selectively impacted behavior in mice.

## Discussion

To edit brain circuits in mammals, we created an electrical synapse based on two Cx36 homologues. All our preparations supported the formation of heterotypic gap junctions between Cx34.7 _M1_, and Cx35_M1_, and the physiological and behavioral outcomes of circuit editing were only observed in mice when we expressed these hemichannels under heterotypic conditions (Fig. 6). We also verified that these hemichannels did not modify behavior when expressed heterotypically against Cx36 and Cx43 (Supplemental Fig. S9). Thus, we believe that our results support the use of our newly engineered electrical synapse for precision circuit editing in mammals (Fig. 5 and 6).

Like established protein-based modulation tools such as optogenetics and DREADDs, LinCx can be targeted to precise cell types in mammals. However, LinCx builds upon these technologies by enabling each hemichannel to be expressed in a different cell type. The hemichannels expressed by these two distinct cell types then integrate *in vivo* to form an electrical synapse. As such, LinCx offers unprecedented spatial precision compared to optogenetics and DREADDs in that it enables targeting of one of the specific spatial features that constrain circuits (e.g., the structural integration of two distinct cell types). Moreover, LinCx does not require an exogenous actuator such as light, electricity, or a pharmacological compound. Rather, LinCx utilizes endogenous brain activity to modulate target neurons, yielding a tool for precise circuit editing. Of critical importance, the impact of LinCx editing on circuit function is dependent on the physiological properties of the cell types composing a circuit, including their resting membrane potentials, input resistance, and basal firing rates. Thus, LinCx editing is unlikely to yield identical physiological outcomes across all neural circuits.

The integrated engineering approach we utilized to develop LinCx (see Fig. 2) can likely be deployed to develop a toolbox of Cx protein pairs that exhibit selective docking properties. Future work may also yield novel hemichannel pairs with customized conductance properties, mirroring approaches applied to modify the conductance of invertebrate electrical synapses^34^. Thus, it may be possible to use multiple LinCx pairs in the same animal to simultaneously edit distinct circuits components and regulate behavior. LinCx can also be deployed alongside other well-established preclinical modulation approaches including DREADDs and optogenetics, enabling broad manipulation of brain networks across multiple scales of spatial, temporal, and context resolution concurrently. Finally, LinCx could potentially be used to edit neural circuits outside of the CNS for therapeutic purposes, or to enhance emerging cellular-based therapies that utilize or target specific Cx-expressing cells.

## Limitations

There are several important limitations of our LinCx approach. First, like prior circuit-editing approaches based on inserting gap junctions between specific cell types, our approach is only suitable for editing circuits composed of cells that make physical contact. Second, LinCx has the potential to yield mixed synapses (in neurons) for which chemical and electrical synapses operate in parallel. Since our Cx channels are chronically expressed, we anticipate that LinCx also induces activity-dependent changes in local chemical synapses. Indeed, this plasticity at chemical synapses may contribute to the physiological and behavioral changes we observed in our mouse assays.

We engineered Cx34.7 and Cx35 mutant proteins to be docking-incompatible with Cx43 and Cx36. Since there are other Cx proteins expressed by mammals, we also used FETCH to screen our Cx mutants for heterotypic docking with other human connexin proteins. We observed FETCH scores for Cx31.3 and Cx37 that were higher than the docking-incompatible pairs we used for our initial analysis (Supplemental Fig. S16), but lower than the docking-compatible pairs. Cx31.3 is expressed in the parenchyma of the mammalian central nervous system^52^, while Cx37 is not. Finally, we were unable to quantify docking for all human isoforms using FETCH analysis (Supplemental Fig. S16). Thus, future work is warranted to assess the docking compatibility of our mutant pairs with Cx31.3 in vivo, and the functional significance of any additional putative docking interactions, for CNS applications of LinCx.

Our engineered hemichannels failed to modify circuit function and behavior when expressed homotypically. For Cx35_M1_, these findings were consistent across our FETCH, paired oocyte, and mouse experimental findings. Yet, our paired oocyte electrical recordings indicated that Cx34.7_M1_ formed functional homotypic channels. To better understand the potential impact of this discrepancy on the function of our mutated proteins in mammals, we also quantified the function of Cx34.7_M1_ in an engineered HEK293FT cell line. Here, we found that homotypic expression of Cx34.7_M1_ failed to induce functional channels (Fig. 3L), clarifying that the discrepancy was solely observed in the *Xenopus* oocytes system. This discrepancy may be due to protein interactions that restored the docking capacity of Cx34.7_M1_ in *Xenopus* oocytes system. Future experiments may help to clarify these mechanisms, and aid in further optimization of Cx34.7_M1_.

Finally, we cannot exclude the possibility that our mutants oligomerize with endogenous Cx36 in mammals, yielding heteromeric hemichannels. Indeed, any such heteromeric channels may exhibit docking properties that are distinct from Cx36, Cx34.7_M1_, or Cx35_M1_ exclusive hemichannels (homomeric), ultimately limiting the functionality of LinCx across some neural circuits. Future work to assess and optimize the oligomerization specificity for Cx34.7_M1_ and Cx35_M1_ may further enhance the applicability and utility of LinCx to mammalian neural circuit editing. Future work may also yield new features to regulate the cellular level directionality of Cx34.7/Cx35 *in vivo*^15^.

## Acknowledgements

The authors would like to acknowledge Dr. Terry Oas, Dr. Mike Tadross, Dr. Nicole Calakos, Dr. Marc Caron, Dr. David Carlson, and Dr. Fan Wang for helpful discussion and/or guidance; Tatiana Rodriguez, Dr. Kayla Lemons, Dr. Gary Sutherland, Hannah Soliman, Amanda Benton, and Reagan Portelance for technical assistance; Dr. Julia Derk, Dr. Lin Shan, and Dr. Giorgia Boero for validating the Cx double knockout cell line, preparing materials used in physiological and histological analyses, and preparing accompanying figures/legends; Dr. John O’Brien for supplying reagents; and Dr. Jean Mary Zarate for comments/feedback on the manuscript and editorial guidance All other technical contributions are acknowledged in prior preprints of this work^53^. The authors would like to acknowledge the generous funding that has supported this work: Ernest E. Just Life Science Institute Postdoctoral Research Fellowship; Burroughs Wellcome Fund Postdoctoral Enrichment Program Grant (E.R.); Hartwell Postdoctoral Research Fellowship (E.R.); SOM Core Facility Voucher Program (E.R. and K.D.); NIH grants R01NS076558 and DP1NS111778 (D.C.R.); HHMI Scholar Award (D.C.R.); NIH grants U01HL134764, R01HL132389, and R01HL126524 (N.B.); Duke University School of Medicine MedX Grant (K.D. and N.B.); Duke University Chancellor’s Discovery award (K.D. and N.B.); Duke University School of Medicine Kahn Neurotechnology Grant (E.R., N.B., and K.D.); NIH BRAIN Initiative Grant R21EY029451 (K.D. and N.B.); Hope for Depression Research Grant, and NIH Grants R01MH120158, R01MH125430 and 1DP1MH132709 (K.D.). A special thanks to Freeman A. Hrabowski, Robert and Jane Meyerhoff, and the Meyerhoff Scholarship Program.

## Author contributions

Conceptualization and Methodology – E.R., E.W., A.A.P., D.C.R., R.H., and K.D.; Formal Analysis – E.R., G.E.T., E.W., T.R., K.K.C.W., D.H., A.A.P., S.D.M., L.N., Z.W., and D.C.R., K.D.; Investigation – E.R., G.E.T., E. W., A.A.P., R.B., T.R., E.A., K.K.C.W., D.H., H.S., S.D.M., R.H.; Resources – E.R., Z.,W., N.B., D.C.R., K.D.; Writing – Original Draft, E.R., E.W., A.A.P., D.C.R., N.B., K.D. Writing – Review & Editing, E.R., G.E.T., E.W., T.R., A.A.P., K.K.C.W., D.C.R., L.N., Z.W., S.D.M., R.H., N.B, K.D. Visualization – E.R., G.E.T., E.W., T.R., A.A.P., L.N., Z.W., D.C.R., K.D.; Supervision – S.D.M., L.N., D.C.R., N.B. and K.D.; Project Administration and Funding Acquisition – E.R., Z..W., D.C.R., R.H., N.B., and K.D. See Supplemental materials for detailed author contributions.

## Materials and Methods

### Design of Cx34.7 and Cx35 mutant library

A semi-rational design approach was used to design the mutant library. Sequence alignments between the *Morone americana* connexins (Cxs) and the Cxs for which the most structure-function data existed (Cx26, Cx32, Cx36, Cx40, and Cx43) were performed in ClustalW. Sites identified by previous studies as conferring specificity for docking were used as well as those identified by homology modeling from the structures of Cx26 ^12^. Specifically, we primarily focused on the extracellular loops: four residues at the interface in loop two, KEVE/KDVE (*M. americana* Cx34.7/Cx35) and one residue of EL1. The homologous residues in other Cxs had been demonstrated to be highly tolerant to mutation and critical for docking specificity ^54^. Mutations were modeled in Swiss PDB Viewer using homology models of Cx34.7 and Cx35 from a Cx26 and Cx32 interface structure so as not to create mutations with obvious steric hindrance. A wide range of substitutions were made for these five residues of interest, including both those intended to introduce compatible electrostatic interactions as well as less likely candidates. Mutations were also created targeting other residues nearby and/or adjacent to these five for which there was some published evidence that they contributed to docking specificity. However, our semi-rational approach was such that not as many variants were tried for these more distal site mutations; mutations made in those sites were more conservative with regards to the steric and electrostatic properties of the change.

### Construct cloning and preparation

The initially acquired *Morone Americana* Cx34.7 and Cx35 cDNA constructs failed to express efficiently in HEK 293FT cells. Thus, Cx gene information was procured from the National Center for Biotechnology Information (NCBI, ncbi.nlm.nih.gov) and the Ensembl genome browser (ensembl.org). The human codon-optimized genes were ordered from Integrated DNA Technology as gBlocks Gene Fragments (IDT, idtdna.com). To generate constructs for transient transfection of HEK 293FT cells, genes were subcloned into BamHI and SacI digested mEmerald-N1 (addgene:53976) and piRFP670-N1 (addgene: 45457) vectors using In-Fusion cloning (takarabio.com), resulting in Cx fluorescent fusion proteins, specifically with the fluorescent proteins being adjoined to the Cx carboxy-terminus. Mutant constructs were generated by employing overlapping primers within standard Phusion polymerase PCR reactions to facilitate site-directed mutagenesis.

The Gateway recombination (Invitrogen) system was used to generate all Cx36, Cx34.7, Cx35, wild type and mutant protein *C. elegans* expression plasmids. For PCR-based cloning and subcloning of components into the Gateway system, either Phusion or Q5 High-Fidelity DNA-polymerase (NEB) was used for amplification. All components were sequenced within the respective Gateway entry vector prior to combining components into expression plasmids via a four-component Gateway system ^55^. The different Cx versions were introduced into pDONR221a using a similar PCR-based strategy from plasmid sources ^7,56,57^. Cell-specific promoters were introduced using the pENTR 50 -TOPO vector (Invitrogen) after amplification from genomic DNA or precursor plasmids. Transgenic lines were created by microinjection into the distal gonad syncytium ^58^ and selected based on expression of one or more co-injection markers: Punc-122::GFP, Pelt-7::mCherry::NLS.

### Cell Culture

HEK293FT cells were purchased from Thermo Fisher Scientific (cat# R70007) and were maintained according to manufacturer instructions. Briefly, cultures were grown in 10-cm tissue culture-treated dishes in high-glucose DMEM (Sigma Aldrich, D5796) supplemented with 6mM L-glutamine, 0.1 mM MEM non-essential amino acids, and 1 mM MEM sodium pyruvate in a 5% CO_2_, 37°C incubator. Cells were passaged via trypsinization every 2-3 days or until 60-80% confluency was reached.

### Transient Transfection

HEK293FT cells were plated in 10ug/ml Fibronectin coated multi-well dishes to achieve ∼75% confluency after overnight incubation. For transfection, 250ng DNA was combined with polyethylenimine (PEI) diluted in Opti-MEM in a 1:3 ratio (µg of DNA: µL of PEI) and incubated at room temperature for 10 minutes. Following incubation, PEI-DNA complexes were added dropwise to wells of plated cells. Treated cells were then incubated at 37°C for 16-18 hrs, followed by media change. Expression of the Cx-FP constructs were evaluated at 24 and 48 hrs post-transfection via widefield or confocal microscopy and western blotting.

### Flow Enabled Tracking of Connexosomes in HEK 293FT cells (FETCH)

FETCH analysis is fundamentally a two-component system (Supplemental Figure S1). To complete FETCH analysis, replica multi-well plates with HEK 293FT cells were transfected with either of the two components being evaluated. The media of transfected wells was changed 16-18 hrs post transfection and cells were trypsinized. Next, HEK 293FT cells expressing experimental Cx counterparts were combined. The entirety of combined samples was then plated onto new, 10ug/ml Fibronectin coated wells of the same size, resulting in hyperdensity and overconfluency. Following co-plating, samples were incubated for ∼20-24 hrs, allowing cells to make contacts and potentially generate and internalize dually labeled gap junctions. Samples were then trypsinized, resuspended in phosphate buffered saline with 10U/ml DNAse and fixed with paraformaldehyde (f/c of 1.5%). Co-plated samples in 96-well plates were resuspended to a final volume of ∼150 µL, while samples from 24-well plates were resuspended to a final volume of ∼600 µL.

Flow cytometry data was collected on a BD FACSCanto II (2-color experiments and high-throughput 96-well plates; 488nm and 633nm lasers), which utilizes the BD FACSDiVa software. Samples were analyzed in two selection gates prior to their fluorescence evaluation. First, presumable HEK cells were identified by evaluating sample forward vs. side scatter area. Next, single cells were selected by evaluating cells that maintained a linear correlation of forward scatter height to forward scatter area. Finally, the fluorescence profiles of each sample were generated.

### Automated FETCH output processing pipeline

Each FETCH experiment produces “*.fcs” files that contain all the channel data for fluorescence in the sample. Our automated pipeline loads these files, extracts FSC-A, SSC-A, and FSC-H. Depending on the machine used, we either load green channel as 1-A or as FITC-A. For the red channel, we can either have two [APC-A (RFP670) and PE-A (mApple)], or just one: 5-A(RFP670). Next, our code produces two matrices containing SSC-A with FSC-A, and FSC-A with FSC-H respectively.

Our first gate is drawn on the FSC-A vs SSC-A axes to exclude cellular debris that clusters in the lower left corner and the cells that are saturating the laser (at the max of both axes). On an FSC-A vs SSC-A plot, the cellular debris usually is smoothly transitioning into the population of intact cells; therefore, we use a Gaussian kernel density estimator with the estimator bandwidth selection defined by the Scott’s Rule to draw contours around the data in SSC-A vs. FSC-A matrix. We next use a set of heuristics to determine which of the contour lines should be used to define the first gate. Specifically, cellular debris usually clusters below 25000 on both axes, so any contour that includes values at or below this value is excluded. Similarly, any contour within 1000 of the maximum value of each axis is also excluded. Of the remaining contours, the largest one is selected, and an oval equation is fitted to the points defining that contour to attenuate occasional protrusions that tap into the cellular debris population in rare cases. The fitted oval becomes the first gate.

For all the elements inside of the first gate, a second gate is drawn in the axis of FSC-A and FSC-H to exclude non-single cells. For the second gate, first we fit a line to all the points. Next, for each point we find a norm to the fit line and find a standard deviation of all such norms. Using this standard deviation, we define a second gate 4 standard deviations away from the fitted line on both sides and exclude all the points outside of this gate (supplemental Fig. 1C).

Upon applying the first two gates, we finally plot the data with the red fluorophore on the y-axis and the green fluorophore on the x-axis. If a sample contains more than two fluorescent signals the last gate is drawn for each possible combination. Since some readings are below zero due to fluorescence compensation, we shift all the data points by the smallest value along both axes and then take a natural log of fluorescence levels. To achieve the optimal bandwidth for the kernel density estimation, we run a cross-validation grid search algorithm on the points in the log space. Then we fit a Gaussian kernel density with the bandwidth estimated to obtain density contours. For properly expressing samples, we expect a large population of non-transfected cells in the bottom left quadrant of the plot, a population of cells strongly-expressing red fluorophore along the y-axis, and a population of cells strongly expressing green fluorophore along the x-axis (supplemental Fig. S1D). We expect autofluorescence to not exceed 500 on either axis, so the non-transfected population is defined to be below this value along both axes. To draw a tighter bound on the non-transfected population, we choose the first contour whose mean kernel density estimate (kde) value is at or above the 60th quantile (identified as a generalizable heuristic value) of the distribution of kde values within the largest contour (at or below the autofluorescence cutoff). The top-most point of the tight contour defines the horizontal gate and the right-most point is the vertical gate, separating the plot into four quadrants.

The upper left quadrant Q1 corresponds to the cells expressing just the red fluorophore, the upper right quadrant Q2 represents dual-colored cells, the lower right quadrant Q3 -- the cells expressing just the green fluorophore, and the lower left quadrant Q4 represents non-transfected population. The FETCH score is defined as the portion of transfected cells that were dual-colored: 100*[Q2/(Q1 + Q2 + Q3)].

Expecting approximately equal expression levels of each fluorophore, if the number of cells in Q1 is two or more times larger/smaller than the number of cells in Q3, the FETCH score is classified as “dubious” and marked accordingly in the output table. The “dubious” label is also given to samples that have less than 500 cells total after the application of the second gate and to the samples that failed at any of the steps in the pipeline (usually due to poor expression or the absence of cells in the sample). Code is available at: https://github.com/carlson-lab/FETCH.

### *In vitro* screening of Cx34.7 and Cx35 mutants for docking selectivity

For homotypic docking screening analysis, five FETCH replicates were obtained for each mutation. These scores were benchmarked against scores for Cx36 with Cx45 (FETCH=1.2±0.1%, N=54 replicates). For our heterotypic docking screening analysis, five replicates were obtained for each mutant pair. These scores were then benchmarked against scores for wild type Cx34.7 with Cx35 (FETCH=14.7±0.4%, N=49 replicates).

To quantitatively determine whether a connexin pair docked, we determined FETCH scores for the dual fluorescence of cells under conditions where docking was not anticipated. These conditions included pairs of connexins previously established to not show docking: Cx36 and Cx45 (FETCH=0.7±0.0%, N=59 replicates), homotypic Cx23 (FETCH=0.9±0.4%, N=6), Cx36 and Cx43 (FETCH=1.2±0.2%, N=10), and under conditions for which cells were transfected with cytoplasmic fluorophores rather than tagged connexins (FETCH=4.4±0.6%, N=17). These 92 FETCH scores were used as the ‘known-negative’ distribution. FETCH scores from each experimental condition were then compared against the established negative score distribution using a one-tailed t-test, with an alpha threshold normalized to the total number of experimental conditions tested. These FETCH replicates were independent of the replicates utilized for our screening analysis. Statistics are reported as mean±s.e.m, and only uncorrected P values are reported through the text.

### Confocal Imaging Analysis of Gap Junction Partners

For imaging of putative gap junction partners, different populations of HEK 293FT cells were transfected with counterpart connexin proteins, incubated, and combined as described for FETCH analysis.

Combined samples of HEK 293FT cells were co-plated onto 10 ug/ml Fibronectin coated 35-mm, glass-bottom Mattek dishes (cat# P35GC-1.5-14-C). Cells were imaged at ∼20 hrs post-co-plating. Images were acquired on a Leica SP5 laser point scanning inverted confocal microscope using Argon/2, HeNe 594nm and HeNe633nm lasers, conventional fluorescence filters and a 63X, HCX PL APO W Corr CS, NA: 1.2, Water, DIC, WD: 0.22mm, objective. Images were taken with 1024 x 1024 pixel resolution at 200Hz frame rate.

For assessing Cx34.7_M1_::Cx35M1 expression *in vivo* in *C. elegans*, we imaged strain DCR8669 *olaEx5214 [Pgcy-8::CX34.7(E214K, E223K)::GFP; Pttx-3::CX35(K221E)::mCherry; Punc-122::GFP]*. L4 animals were mounted in 2% agarose in M9 buffer pads and anaesthetized with 10mM levamisol (Sigma). Confocal images were acquired with dual Hamamatsu ORCA-FUSIONBT SCMOS cameras on a Nikon Ti2-E Inverted Microscope using a confocal spinning disk CSU-W1 System, 488nm and 561nm laser lines and a CFI SR HP PLAN APO LAMBDA S 100xC SIL objective. Images were captured using the NIS-ELEMENTS software, with 2048px x 2048px, 16-bit depth, 300nm step size, 200ms of exposure time and enough sections to cover the whole worm depth.

### Protein modeling pipeline

Our protein modeling pipeline is based on previously published methodology ^33^ and integrates five components: 1) homology model generation, 2) embedding of proteins in a lipid bilayer and aqueous solution, 3) protein mutagenesis, 4) system minimization, equilibration, and molecular dynamics simulation production run, and 5) residue-wise energy calculation.

#### Homology Modeling

We initially tested five homology modeling software suites: Robetta, SwissModel, Molecular Operating Environment (MOE; Chemical Computing Group ULC, Montreal, QC, Canada, H3A 2R7, 2021), I-Tasser, Phyre2 ^59–67^. A quality assessment suite, — MolProbity ^68–70^ revealed that Robetta models outperformed the rest, based on a set of standard metrics (Ramachandran plot outliers, clashscore, poor rotamers, bad bonds/angles, etc).

Since our aim was to model the extracellular loops responsible for Cx hemichannel docking, we picked all the resolved Cx structures that possessed a high degree of extracellular loop homology to our Cx of interest as the inputs for Robetta. The top homolog hits were generally the same for the three connexins of interest: Cx26 Bound to Calcium (5er7.1) ^71^; Human Cx26 (Calcium-free) (5era) ^71^; Structure of Cx46 intercellular gap junction channel at 3.4 angstrom resolution by cryoEM (6mhq) ^32^; Structure of Cx50 intercellular gap junction channel at 3.4 angstrom resolution by cryoEM (6mhy) ^32^; Structure of the Cx26 gap junction channel at 3.5 angstrom resolution (2zw3) ^72^. Cx34.7 and Cx35 wild type sequences had the greatest homology degree with 6mhq, while Cx36 was most homologous to 5er7.1. We generated three wild type hemichannels for Cx34.7, Cx35, and Cx36.

#### System Assembly

Next, we assembled hemichannels into homotypic and heterotypic gap junctions, embedded them in two double bilayers, dissolved them in water, and added appropriate ion concentrations for the extracellular and two intracellular compartments. The primary software suite used for this modeling step was VMD ^73,74^. We also utilized CHARMM GUI to generate the naturalistic model of a region of a double bilayer ^75–79^. Membrane components were then selected in appropriate proportions to resemble experimentally derived data from a neuronal axonal membrane.

Specifically, since Robetta was unable to model the full gap junction, we merged hemichannels into full homotypic/heterotypic gap junctions in a semi-automated way. First, to make homotypic gap junctions, we loaded the two-homology models for a hemichannel. We then aligned them using the center of mass of the extracellular loops. A slight rotation along the z-axis was implemented for several pairs to optimize their fit. To make heterotypic gap junctions, we created a homotypic gap junction for each hemichannel, aligned the extracellular loops for the two homotypic gap junctions, then removed an opposing hemichannel from each homotypic gap junction (leaving the two different hemichannels aligned). Next, using the constructed gap junction, we aligned two pre-made membrane bilayers with the center of mass assigned as each embedded hemichannel. We then removed membrane molecules that overlapped with the hemichannel or the hemichannel pore. Next, we dissolved the system in water and removed water that overlapped with the lipid bilayer. Extracellular water was then separated to a new file, where Na+, K+, Cl-, and Ca2+ ions were added to yield concentrations mirroring the extracellular environment of mammalian neurons ^80^. Finally, Na+, K+, Cl-, and Ca2+ ions were added to the intracellular space to mirror the intracellular environment of mammalian neurons, and the files containing the embedded Cx hemichannels and extracellular water were merged. Notably, these stages were automated yielding a streamlined progression from a protein-only hemichannel model to a fully embedded gap junction model ready for subsequent simulation and/or mutagenesis.

#### Mutagenesis

We developed a Python command-line tool that utilizes VMD to generate mutation configuration files for subsequent molecular dynamics simulation. Here, we simply specified the Cx hemichannels of interest and the position at which a specific mutation should be introduced.

#### System minimization, equilibration, and molecular dynamics simulation production run

Next, we minimized atomic energies, equilibrated the system, and ran the stable system in a production simulation run. Specifically, molecular dynamics simulation was performed using NAMD ^81^ and was divided into five steps:

1. *Melt lipid tails while keeping remaining atoms fixed (simulate for 0.5 ns)*
2. *Minimize the system, then allow the bilayers and solutions to take natural conformation while keeping gap junction fixed (split in two stages to accommodate reduction in volume of relaxing system; simulate 0.5 ns total)*
3. *Release the gap junction and equilibrate the whole system (simulate 0.5 ns)*
4. *Run minimized and equilibrated system in a production run (simulate 0.5 ns)*

Though molecular dynamics simulation (step 4) is highly reliant on the input file provided by the *System Assembly* process, these steps render the simulation much more robust to modeling imperfections. For example, the membrane model developed though System Assembly is very rigid and has the potential to behave like a solid rather than like a liquid. Thus, melting the lipid tails encourages the model to embody a liquid. Similarly, many atoms in the input file may have unnatural initial energies, such that if they are all released at once, they would start moving at high velocities and the simulation would fail. Therefore, bringing the system to a local energy minimum increases stability. Removing constraints on the water and lipids enables them to surround the gap junction in a naturalistic form. Finally, releasing the constraints on the gap junction enables it to take the most energetically stable conformation given the environment.

#### Energy Calculation

To predict the residues that play a prominent role in docking, we quantified all non-bonding interactions between the two Cx hemichannels at key residues on the extracellular loops. Output from the molecular dynamics simulation was loaded into the VMD “NAMD Energy” plugin. We then calculated nonbonding energies for all residues on each hemichannel that were within 12 angstroms of at least one residue on the other hemichannel. For each residue pair, we then averaged energies across the 250 simulation frames.

### Characterization of gap junction biophysical properties using *Xenopus* oocytes

DNA sequences encoding Cx34.7_WT_-mEmerald, Cx34.7_M1_-mEmerald, Cx35_WT_-mApple, and Cx35_M1_-mApple were cut out from the plasmids utilized for our mouse studies (see below), by BamH1 and ECoR1, and cloned into the pXMX_T3(+) *Xenopus* oocyte expression vector at identical restriction sites. The newly generated plasmids were linearized by cutting with NgoMIV to serve as templates for *in vitro* synthesis of complementary RNAs (cRNAs). Capped cRNAs were synthesized using a mMessage mMachine^TM^ T3 Transcription kit (AM1348, ThermoFisher Scientific). Each cRNA injection solution was a mixture of a specific cRNA and a Cx38 antisense oligonucleotide (5ʹ-GCTTTAGTAATTCCCATCCTGCCATGTTTC-3ʹ, 100 ng/µl).

In preliminary experiments with Cx34.7_WT_ and Cx35_WT_ cRNAs injected at an identical concentration (900 ng/µl), we observed much larger *I_j_* in homotypic gap junction of Cx35_WT_ than those of Cx34.7_WT_ under identical experimental conditions, and as well as cytotoxic effect (oocytes dying) of Cx35_WT_ but not Cx34.7_WT_. Therefore, in the final injection solutions used for the main experiments presented in this paper, the concentrations of Cx34.7_WT_-mEmerald and Cx34.7_M1_-mEmerald cRNAs were 900 ng/µl, whereas those of Cx35_WT_-mApple and Cx35_M1_-mApple cRNAs were 90 ng/µl. Approximately 50.6 nl of the mixture was injected per oocyte using a Drummond Nanoject II injector (Drummond Scientific, Broomall, PA). Injected oocytes were incubated in ND96 solution (96 mM NaCl, 2 mM KCl, 1.8 mM CaCl_2_, 1 mM MgCl_2,_ and 5 mM HEPES [pH 7.5]) inside an environmental chamber (15°C) before being paired.

Oocytes were paired 20-24 hours after Cx35_WT_ and Cx35_M1_ cRNA injections, but 68-72 hours after Cx34.7_WT_ and Cx34.7_M1_ cRNA injections. To closely match experimental conditions between related groups, oocytes from one frog were utilized to analyze homotypic gap junctions of Cx34.7_WT_ and Cx34.7_M1_, while oocytes from another frog were employed to examine homotypic gap junctions of Cx35_WT_ and Cx35_M1_. Furthermore, an identical batch of oocytes from a third frog was used to investigate heterotypic gap junctions of both Cx34.7_WT_/Cx35_WT_ and Cx34.7_M1_/Cx35_M1_. The vitelline membrane of the oocyte was removed using a pair of #5 Dumont Tweezers (500342, Word Precision Instruments, Sarasota, FL) during the pairing process. Paired oocytes were incubated in ND96 solution for 12-24 hours at room temperature (20-22°C) prior to electrophysiological recordings. Voltage-clamp recordings were performed using two Oocyte Clamp amplifiers (OC-725C, Warner Instruments LLC, Hamden, CT) in the high-side current measuring mode. In the electrophysiological experiments, we applied a series of membrane voltage steps (−150 mV to +90 mV at 10-mV intervals, 7 sec in duration) to one oocyte from a holding voltage of -30 mV while holding the other oocyte constant at -30 mV to record transjunctional currents (*I_j_*). A pre-pulse (−30 mV) was applied from the holding voltage before each voltage step so that stability of the recordings could be monitored by the induced *I_j_*. Instantaneous current (*I_inst_*) and steady-state current (*I_ss_*) were defined as the peak *I_j_* at the beginning of each *V_j_*step and as the averaged *I_j_* during the last 2 sec of the *V_j_*step, respectively. Details of the experimental and data-analysis procedures can be found in our previous publications^34,35^.

### Generation and validation of Cx43 and Cx45 double knockout HEK294FT cells

Cx double knockout HEK293FT cells were produced by Duke Functional Genomics Core Facility using CRISPR/Cas9 technology. Specifically, indel (insertion and/or deletion) mutations were introduced via CRISPR-Cas9 in the Cx43 and Cx45 sequences to induce frameshifts within the Cx43 (*GJ1A*) and Cx45 (*GJ1C*) DNA sequence resulting in premature translation termination (Supplemental Fig S6A-C). Specific mutations were identified via DNA sequencing and evaluation of sequence chromatograms (exported as encapsulated postscript (.eps) files) in ApE Plasmid editor^82^. To confirm Cx43 and Cx45 knockout, three double knockout HEK 293FT monoclonal cell lines were isolated and expanded for evaluation via western blotting. Cells were harvested in ice-cold PBS (Gibco #10010031) enriched with protease and phosphatase inhibitors 1:100 (Sigma-Aldrich #P8340 and #P5726). Cells were then centrifugated at 1000xg for 5 min at 4 °C to remove the PBS, and the pellet was sonicated twice for 30 sec (20% output power) in ice-cold lysis buffer (Phosphosolutions #NC1671658) enriched with protease and phosphatase inhibitors 1:100 (Sigma-Aldrich #P8340 and #P5726). Cell lysates were kept on ice for 30 min and then centrifugated at 14,000× g for 30 min at 4 °C. Lysate supernatants were collected, and total protein concentrations were measured via bicinchoninic acid assay (Thermo Fisher Scientific #23228, #1859078). 40μg of total protein were denatured at 95°C for 5 min in LDS sample buffer (7 µL per sample) (Thermo Fisher Scientific #NP0007) and sample reducing agent (3 µL per sample) (Thermo Fisher Scientific #NP0009). The denatured samples were then separated via Novex™ Tris-Glycine Gels 4-20% (Thermo Fisher Scientific #XP04200BOX) electrophoresis at 125V for 10 min and 150V for ∼1 hour, then transferred to polyvinylidene fluoride membranes (PVDF) (Thermo Fisher Scientific #88520) overnight at 0.04A. Membranes were blocked with 5% dry non-fat milk (Blotting-Grade Blocker, Bio-Rad #1706404) in PBST (0.1% Tween-20) for at least 2 hours at room temperature and next incubated overnight at 4°C with the following antibodies in blocking solution: anti-Cx43 (Cell Signaling Technologies #3512, 1:1000), anti-Cx45 (Abcam #ab316742, 1:1000). GAPDH was measured after the protein of interest (Cx43 or Cx45) as control (anti-GAPDH, Abcam #ab181602, 1:5000). After incubation with the primary antibody, this was removed and blots were washed in PBST 0.1% (3 washes, 15 min each) at room temperature, following incubation with horseradish peroxidase-labeled secondary anti-rabbit IgG antibody (Cell Signaling Technologies, #7074) for 1h at room temperature. Membranes were washed in PBST 0.1% to remove the secondary antibody (3 washes, 15 min each). Immunoreactive bands were visualized with SuperSignal™ West Pico PLUS Chemiluminescent Substrate (Thermo Fisher Scientific #34578) and detected with ChemiDoc Imaging System (BioRad #12003153). Bands were analyzed with ImageJ. For each protein studied, the ratio was calculated dividing the densitometric value of the protein of interest by the correspondent GAPDH value *100 (Supplemental Fig. S6D).

### PCR confirmation of Cx43 and Cx45 knockouts

Genomic DNA from HEK293FT double knock out cell line clone of #405, #412 and #422 was extracted using KAPA NGS DNA extraction kit (Roche. Cat# 09189823001), and PCR was performed using Invitrogen Platinum super Fi PCR Master Mix (Cat# 12358010), following manufacturer’s instructions. PCR products were prepared by adding a DNA stain (APEBIO #ab743) and then separated in 1% agarose gel (BioRad #1613102) in TAE buffer (BioRad #1610743) at 100V for ∼30 min. DNA fragments were detected with ChemiDoc Imaging System (BioRad #12003153).

### Wild type and Cx double knockout HEK293FT cultures for patching experiments

Wild type HEK293FT cells were purchased from Thermo Fisher Scientific (cat# R70007). Cx double knockout HEK293FT cells were generated as described above. Cells were maintained according to manufacturer instructions. Briefly, cultures were grown in 10-cm tissue culture-treated dishes in high-glucose DMEM (Sigma-Aldrich, D5796) supplemented with 6mM L-glutamine, 0.1 mM MEM non-essential amino acids, and 1 mM MEM sodium pyruvate in a 5% CO2, 37° C incubator. Cells were passaged via trypsinization every 2-3 days or until 60-80% confluency was reached.

### Glass coverslips preparation

12-mm German glass coverslips (NeuVitro cat#GG-12-15-Fibronectin) were first pre-treated with acetone for 20 minutes with shaking. Next, they were washed 3 x 10 minutes in 70% ethanol with shaking and stored in 70% ethanol before use. Once needed, each coverslip was washed 6 times with cell culture-grade deionized water to remove any ethanol remnants. For the patching experiments, coverslips were then coated using 1:100 Fibronectin (Sigma-Aldrich cat# F1141-2MG) (stock at 1mg/mL) in Poly-D-Lysine (Gibco cat# A38904-01) (stock at 0.1 mg/mL in PBS) for one hour at room temperature. After coating, the solution was aspirated and the coverslips were left to dry in a cell culture hood, uncovered, for 1-2 hours.

### Sample preparation for paired cell recordings

For the recording experiments using non-transfected cells, HEK293FT or Cx double knock out HEK293FT cells were grown in 10 cm tissue-culture treated dishes and seeded onto the coverslips, at the same density conditions used for all patching experiments (35,000 cells per well in total), the day before the recording. To evaluate connexin proteins, Cx double knockout HEK293FT cells were seeded with no antibiotic cell culture medium, and the next day they were transfected using Lipofectamine 2000 transfection reagent (Thermo Fisher cat# 11668027) at a Lipofectamine 2000 to DNA ratio 2.5-1.9:1. (cDNA 0.8-1 *u*g; Lipofectamine 1.9–2µl). Opti-MEM (Gibco cat# 31985070) medium was used following the Lipofectamine 2000 protocol. Media was changed after 4 hours to remove the transfection reagent. For each patching condition, 4 or 20 hours after transfection, cells were detached, counted and re-seeded in equal amounts of cells from each fluorescent condition (35,000 cells per well in total) onto a precoated glass coverslip in 24-well plate (see section above for coverslip preparation). The cells were left to form gap junctions overnight and recordings were conducted the following day. These patching experiments were performed using the construct cloning and preparation procedures described above, with minor changes. Briefly, for the Cx34.7_M1_/Cx35_M1_ or Cx36/Cx36 patching conditions tested, two fluorescent tags were used (mEmerald or mCherry). For two of the four Cx double knockout HEK293FT negative control experiments, we used cell pairs that expressed mEmerald vs. mCherry. Finally, we used only one fluorescent tag for the Cx34.7_M1_/Cx34.7_M1_ or Cx35_M1_/Cx35_M1_ experiments (mEmerald, and mCherry respectively).

### Paired HEK293FT cell recordings

Glass coverslips with adhered wild type or Cx double knockout HEK293FT cells were placed into a recording chamber on an upright fluorescent microscope (Scientifica). The chamber was filled with extracellular solution (ECS) at room temperature (21–24°C). The ECS composition was as follows (in mM): 5.4 KCl, 1 CaCl2, 140 NaCl, 5 HEPES, 1.8 MgCl2, 5 glucose; pH was adjusted to 7.4 with NaOH with an osmolarity of 310-320 mOsm. Patch pipettes were pulled using a micropipette puller (Sutter Instruments P1000) and filled with intracellular solution containing (in mM): 135 CsCl, 0.5 CaCl2, 2 MgCl2, 5.5 EGTA, 5 HEPES, 3 MgATP, 2 Na2ATP; pH was adjusted to 7.2 with CsOH, with an osmolarity of 290 mOsm. Patch pipette resistance was in the range of 2–4 MΩ. Isolated cell pairs were selected for dual whole-cell voltage clamp experiments.

To study transjunctional currents between cell pairs, the following protocol was applied in voltage clamp. A series of square voltage pulses (stepped up from -160 mV to +70 mV in 10-mV increments) was alternated between the two cells. While the first cell received the pulses, the second cell was held at -45 mV to measure the transjunctional current. This procedure was repeated for both cells.

### Analysis of transjunctional current

The baseline current was measured 500 ms before the voltage pulse. The current for each 200ms voltage step was then determined as the difference between the current recorded during the last 50 ms of each step and this baseline current. The relationship between this current and transjunctional voltage was determined for each cell with a Pearson correlation and the values were averaged across cells within a cell pair (Supplemental Fig. S7). The current at the 70mV step was extracted for each cell and averaged across the pair to determine the max transjunctional current used for analysis.

To test Cx36, we co-plated cells expressing red or green fluorescent proteins. Because Cx36 shows homotypic docking, we patched fluorescent cell pairs irrespective of their color. To test Cx34.7_M1_/Cx35 _M1_, we patched pairs for which one of the cells expressed mEmerald-tagged Cx and the other cell expressed mCherry-tagged Cx. We then separated experimental groups for which both cells showed green and red (indicative of connexosome formation) from the pairs in which both cells did not (Supplemental Fig. S7). To test Cx34.7_M1_ or Cx35_M1_, we patched cell pairs expressing a single fluorophore.

### Statistical analysis of patching data

The max transjunctional voltage recorded from wild type HEK293FT cell pairs, or pairs of Cx double knockout HEK293FT cells transfected with connexins, were compared to a Cx double knockout HEK293FT cell pair control group using a Wilcoxon rank-sum test, followed by a false discovery rate correction. Notably, in contrast to the published methodology using HEK293 cells^36^, knocking down Cx43 and Cx45 did not completely abolish electrical coupling between HEK293FT cells. Indeed, in our preliminary experiments, we observed electrical coupling in 1/10 Cx double knockout HEK293FT cell pairs (as indicated by a transjunctional voltage x current relationship > R^2^=0.75 using a Pearson correlation). In our main experiments, we observed coupling in 3/20 of our negative control pairs (2/9 Cx double knockout HEK293FT cell pairs and 1/11 of the pairs that expressed Cx34.7_M1_/Cx35_M1_ but did not exhibit dual fluorescence in both cells; see Supplemental Figure S7B). In our post-hoc testing, we observed a significantly higher portion of coupled cells in our two positive controls groups (7/10 Cx36/Cx36 cell pairs, χ^2^ = 7.8 and P=0.002 compared to the pooled negative controls using a Chi-squared test; 7/10 Cx34.7_M1_/Cx35_M1_ that exhibited dual fluorescence in both cells, χ^2^ = 7.8 and P=0.002, with Yates correction, P<0.05). These findings increased our confidence that this mammalian cell system could be used to test Cx34.7_M1_/Cx34.7_M1_ and Cx35_M1_/Cx35_M1_ coupling. Indeed, the coupling observed in these groups was statistically indistinguishable from the pooled negative control group (0/10 Cx34.7_M1_/ Cx34.7_M1_ cell pairs, χ^2^ = 1.7 and P=0.20, with Yates correction, P>0.05; 3/10 Cx35_M1_/ Cx35_M1_ cell pairs, χ^2^ = 0.9 and P=0.33, with Yates correction, P>0.05).

### *C. elegans* strains and genetics

Nematodes were maintained at 20°C on NGM plates seeded with a lawn of *Escherichia coli* strain OP50 using standard methods ^83^. All worm experiments were performed using one-day-old adult hermaphrodites. The strains used in this study are listed in the Strain Table (Supplemental Table S2). All thermotaxis behavioral assays, calcium imaging experiments and generation of transgenic lines were performed as previously described in Hawk et al. 2018 ^9^, with minor modifications as outlined in the following subsections.

### Thermotaxis Behavioral Assay

Animals were grown and assayed as previously described in Hawk et al. 2018 ^9,84^. Briefly, after being reared at 20°C, animals were trained at 15°C for 4 hours prior to testing. Their migration tracks were analyzed as previously described ^9,84,85^. Each behavioral arena was split in half along the temperature gradient using a thin and clear plastic divider. This allowed for wild type controls to be assayed on one half of the arena and Cx-expressing animals on the other half, simultaneously.

### Calcium Imaging in *C. elegans*

Imaging calcium dynamics for assaying AFD-AIY functional coupling was performed as previously described ^9^, with the following modifications: a Leica DM6B was used instead of Leica DM5500, and image acquisition was performed using MicroManager ^86^. Segmentation into regions of interest (ROIs) and downstream data processing was performed using FIJI ^87^ and custom scripts written in MATLAB (MathWorks) were used as detailed previously ^9^. For analyses of AFD calcium transients, we generated and measured an ROI around a single AFD soma per animal. For analyses of AIY calcium dynamics, we generated and quantified an ROI at the synaptic subcellular region known as Zone 2 ^88^. Responses were scored as the initial rise of the AFD or AIY calcium signal as determined by a blind human observer. The genetic background for the AFD and some of the AIY calcium imaging lines used in this study (control and experimental) contained *olaIs23*, a caPKC-1 GOF mutation. This was done to match prior work ^9^ in which Cx36 was demonstrated to evoke AFD-locked responses in AIY compared to caPKC-1 animals without Cx36.

### Vertebrate Animal Care and Use

Male B6.129P2-*Pvalb^tm^*^1^(cre)*^Arbr^*/J (PV-Cre mice) and female C57BL/6J mice (Stock No: 017320 and 000664, respectively) purchased from Jackson labs were bred to generate the male and female (N=28 total virally injected and implanted mice) PV-Cre heterozygous mice subjected to the prefrontal cortex PYR-PV+ editing experiment quantifying LFP coupling. Male C57BL/6J mice (N=29) purchased from Jackson were utilized for non-edited controls. Mice were housed 3-5/cage on a 12-hour light/dark cycle and maintained in a humidity- and temperature-controlled room with water and food available *ad libitum*.

Neural recordings were conducted during the dark cycle (Zeitgeber time: 13-19), given prior evidence that electrical synapse conductance can be diminished in the retina via circadian regulation ^89^. PV-Cre mice were crossed with B6;129S-*Slc17a6^tm^*^1.1^(flpo)*^Hze^*/J (Sock No: 030212) purchased from Jackson labs to obtain the mice used for PV-Cre/Vglut2-flp mice subjected to the prefrontal cortex PYR-PV+ editing experiment quantifying cellular coupling. Groups were balanced by age and sex. Inbred BALB/cJ male mice (strain: 000651) purchased from the Jackson Labs were used for infralimbic cortex (IL) ◊ medial thalamus (MD) circuit editing experiments. We chose this strain and sex of mice to mirror our previous study which optogenetically targeted the IL→ MD circuit ^48^. Behavioral and physiological experiments were conducted during the dark cycle (Zeitgeber time: 13-22). All vertebrate animal studies were conducted with approved protocols from the Duke University Institutional Animal Care and Use Committees and were in accordance with the NIH guidelines for the Care and Use of Laboratory Animals.

### Generation of mouse viral constructs

The mouse codon-optimized wild type and mutant Cx genes were ordered from Integrated DNA Technology as gBlocks Gene Fragments (IDT, idtdna.com). To generate fluorescently tagged Cx constructs for viral transformation of mouse neurons, gBlocks were ligated into BamHI and EcoRI digested pAAV-CaMKIIa-EGFP (addgene:50469). To generate fluorescently tagged Cx constructs for viral transformation of Cre-expressing neurons, Cx constructs were amplified from aforementioned CaMKIIa constructs and ligated into AscI and NheI digested pAAV-Ef1a-DIO-EYFP (addgene: 27056). Finally, to generate FLAG-tagged Cx constructs with co-expressed cytoplasmic fluorescent proteins, a gblock Gene Fragment of Cx35-FLAG-T2A-mCherry was ordered from IDT, amplified and ligated into AscI and NheI digested pAAV-hSyn-DIO-eGFP (#124; addgene: 50457*). All ligations were accomplished using In-Fusion cloning (takarabio.com). AAV viruses were created through the Duke Universitys viral vector core.

### Mouse viral injection surgeries

For the PYR-PV+ editing experiment using microwires, PV-Cre mice were anesthetized with isoflurane (1%), placed in a stereotaxic device, and injected with a 1:1 solution of AAV9-CaMKII-Cx34.7_M1_-mEmerald (titer: 5.0×10^12^ vg/ml) and AAV9-Ef1α-DIO-Cx35_M1_-mApple (titer: 1.3×10^13^ vg/ml), based on stereotaxic coordinates measured from bregma at the skull to target prelimbic cortex bilaterally (PrL: 2.1mm AP, 0.65mm ML, -1.45mm DV from the dura at a 21⁰ angle for male mice, and PrL: 2.05mm AP, 0.62mm ML, -1.41mm DV from the dura at a 21⁰ angle for female mice). A total of 1µL viral solution was delivered to each hemisphere over 10 minutes using a 5µL Hamilton syringe. This strategy yielded expression of Cx35_M1_ solely by PV+ interneurons, and non-selective expression of Cx34.7_M1_. Control mice non-selectively expressed Cx34.7_M1_ or Cx35_M1_. In these mice, we injected with a 1:1 solution of AAV9-CaMKII-Cx34.7_M1_-mEmerald and AAV9-Ef1α-DIO-Cx34.7_M1_-mApple (titer: 1.1×10^13^ vg/ml) or a 1:1 solution of AAV9-CaMKII-Cx35_M1_-mEmerald (titer: 6.9×10^12^ vg/ml) and AAV9-Ef1α-DIO-Cx35_M1_-mApple, respectively, to mirror the injection conditions of the experimental group. All viruses were created in the Duke Viral Vector Core or purchased from Addgene. Viral injections were performed in male and female mice at age 2.5-5 months, and viral manipulations were balanced across cages and sex.

For the PYR-PV+ editing experiment using silicon probes, PV-Cre /VGLUT2-flp mice were anesthetized with isoflurane (1%), placed in a stereotaxic device, and injected with a 1:1 solution of AAV9-CaMKII-Cx34.7_M1_-mEmerald (titer: 1.4×10^13^ vg/ml) and AAV9-hsyn-DIO-Cx35_M1_-T2A-mCherry (titer: 1.3×10^14^ vg/ml), based on stereotaxic coordinates measured from bregma at the skull to target medial prefrontal cortex bilaterally (medial prefrontal cortex: 1.8mm AP, 0.3mm ML, -2.85mm and -2.5mm DV from the skull for male mice, and 1.75mm AP, 0.29mm ML, -2.78mm and -2.44mm DV from the skull for female mice). A total of 0.3µL viral solution was delivered to each DV target for each hemisphere over 10 minutes using a 5µL Hamilton syringe. This strategy yielded expression of Cx35_M1_ solely by PV+ interneurons, and non-selective expression of Cx34.7_M1_. Control mice solely expressed fluorophores. In these mice, we injected with a 1:1 solution of AAV9-CaMKII-eGFP (titer: 2.8×10^13^ vg/ml) and AAV9-hSyn-DIO-mCherry (titer: 1.9×10^13^ vg/ml). All viruses were created in the Duke Viral Vector Core or purchased from Addgene. Viral injections were performed in four female and two male mice at age 2.5-5 months, and viral manipulations were balanced across cages and sex.

For the IL◊MD behavioral experiment, 3 month old male BALB/cJ mice were anesthetized with isoflurane (1%), placed in a stereotaxic device, and injected with AAV9-CaMKII-Cx34.7_M1_-mEmerald (titer: 5.0×10^12^ vg/ml) based on stereotaxic coordinates measured from bregma at the skull to target IL bilaterally (1.7mm AP, ±0.72mm ML, -2.03mm DV from the dura at an angle of 10°). A total of 0.5µL viral solution was delivered to each hemisphere over 5 minutes using a 5µL Hamilton syringe, which was left in place for an additional 10 minutes prior to removal. Three weeks later (see Supplemental Fig. S11), mice were injected with AAV9-CaMKII-Cx35_M1_-mApple (titer: 6.9×10^12^ vg/ml) based on stereotaxic coordinates measured from bregma at the skull to target MD bilaterally (1.58mm AP, 0.5mm ML, - 2.88mm DV from the dura at an angle of 10°). Control mice were injected with AAV9-CaMKII-Cx34.7_M1_-mEmerald in both IL and MD, or AAV9-CaMKII-Cx35_M1_ in both IL and MD, to express the mutant hemichannels in non-docking homotypic configurations. For the IL→MD circuit interrogation study, 3 month old male BALB/cJ mice were injected with a 1:1 solution of AAV9-CaMKII-Cx34.7_M1_-mEmerald (titer: 2.3×10^13^ vg/ml) and AAV9-CamKII-ChR2-EYFP (titer: ≥ 1×10¹³vg/ml) to target left IL unilaterally (1.7mm AP, 0.72mm ML, measured from bregma; - 2.03mm DV from the dura at an angle of 10°). A total of 0.5µL viral solution was delivered at a rate of 100nL/minute. Three weeks after the first surgery, mice were again anesthetized, placed in a stereotaxic device and injected with either AAV9-CaMKII-Cx35_M1_-mApple (titer:3.16×10^13 vg/ml) or AAV9-CaMKII-EGFP (titer: 2.3×10^13^ vg/ml) to target left MD unilaterally. A total of 0.5µL viral solution was delivered. Viruses were infused at a rate of 100nL/minute.

### Electrode Implantation Surgery

For the PYR-PV+ experiment, PV-Cre mice were anesthetized with isoflurane (1.0%), placed in a stereotaxic device, and metal ground screws were secured to the cranium. A total of 8 tungsten microwires were implanted in prelimbic cortex (centered at 1.8mm AP, ±0.25mm ML, -1.75mm DV from the dura for male mice; centered at 1.76mm AP, ±0.25mm ML, -1.71mm DV from the dura for female mice). C57BL/6J control mice were implanted at 2 months. A total of 32 tungsten microwires were arranged in our previously described multi-limbic circuit recording design^90^. Briefly, bundles were implanted to target basolateral and central amygdala (Amy), medial dorsal thalamus (MD), nucleus accumbens core and shell (NAc), VTA, medial prefrontal cortex (mPFC), and VHip were centered based on stereotaxic coordinates measured from bregma (Amy: -1.4mm AP, 2.9 mm ML, -3.85 mm DV from the dura; MD: -1.58mm AP, 0.3 mm ML, -2.88 mm DV from the dura; VTA: -3.5mm AP, ±0.25 mm ML, - 4.25 mm DV from the dura; VHip: -3.3mm AP, 3.0mm ML, -3.75mm DV from the dura; mPFC: 1.62mm AP, ±0.25mm ML, 2.25mm DV from the dura; NAc: 1.3mm AP, 2.25mm ML, -4.1 mm DV from the dura, implanted at an angle of 22.1°). We targeted cingulate cortex, prelimbic cortex (PrL), and infralimbic cortex (IL) using the mPFC bundle by building a 0.5mm and 1.1mm DV stagger into our electrode bundle microwires. Animals were implanted bilaterally in mPFC and VTA. All other bundles were implanted in the left hemisphere. The NAc bundle included a 0.6mm DV stagger such that wires were distributed across NAc core and shell. We targeted basolateral amygdala (BLA) and central amygdala (CeA) by building a 0.5mm ML stagger and 0.3mm DV stagger into our AMY electrode bundle^90^.

For the IL◊MD circuit interrogation study, 16 tungsten microwires were arranged into two bundles to target IL (8 wires) and MD (8 wires). The IL bundle was also built with an optical fiber (MFC_100/125-0.22_8.0mm_MF2.5_FLT, Doric Lenses) 0.5mm above the tip of the wires as previously described^48^. An optic fiber was also built into the MD microwire bundle for two-thirds of the animals, although it was not utilized for this study. Mice were anesthetized as described above, and metal ground screws were secured to the cranium. Bundles were implanted in left IL (1.7mm AP, 0.15mm ML, from bregma; - 2.25mm DV from the dura) and left MD (centered at 1.58mm AP, 0.35mm ML, -2.88mm DV from the dura).

### Data acquisition for PrL microcircuit editing

Neural recording experiments were performed in PV-Cre mice one week after implantation surgery and blind to viral group. Mice were habituated to the recording room for at least 60 minutes prior to testing. PV-Cre mice were connected to a headstage (Blackrock Microsystems, UT, USA) without anesthesia, and given a single saline injection (10mL/kg mouse, intraperitoneally). Notably, these saline injections were performed to facilitate comparison of acquired neural data with future drug studies. Twenty-five minutes later, mice were placed in a 17.5in × 17.5in × 11.75in (L×W×H) chamber for 60 minutes.

Recordings were conducted under low illumination conditions (1-2 lux), and only data from the first 10 minutes of exposure to the open field were used for neurophysiological analysis. For C57BL/6J control mice, experiments were performed at least two weeks following implantation surgery. Mice were habituated to the recording room for at least 60 minutes prior to testing, and headstages were connected without anesthesia. Twenty-six male mice were placed in a 17.5in × 17.5in × 11.75in (L×W×H) chamber for 10 minutes, and three mice were recorded in a 19.5in × 12in (D×H) circular chamber. Six of these mice were injected with saline (10mL/kg mouse, intraperitoneally) 30 minutes prior to recordings, and all recordings were conducted under an illumination of 125 lux.

Neuronal activity was sampled at 30kHz using the Cerebus acquisition system (Blackrock Microsystems Inc., UT). Local field potentials (LFPs) were bandpass filtered at 0.5–250Hz and stored at 1000Hz. All neurophysiological recordings were referenced to a ground wire connected to both ground screws, and an online noise cancellation algorithm was applied to reduce 60Hz artifact.

### Determination of LFP cross-frequency phase coupling and spectral power

Signals recorded from all viable implanted microwires were used for analysis. LFPs were filtered using 4^th^-order Butterworth bandpass filters designed to isolate theta (4-10Hz) prefrontal cortex oscillations and high frequency oscillations (HFOs, 80-200Hz). The instantaneous amplitude and phase of the filtered LFPs were then determined using the Hilbert transform, and the Modulation index was calculated for each LFP channel using the MATLAB (The MathWorks, Inc., Natick, MA) code provided by Canolty et al^45^. Briefly, a continuous variable z(t) is defined as a function of the instantaneous theta phase and instantaneous gamma amplitude such that z(t) = A_G_(t)*e^iϕ^_TH_(t), where A_G_ is the instantaneous gamma oscillatory amplitude, and e^iϕ^_TH_ is a function of the instantaneous theta oscillatory phase. A time lag is then introduced between the instantaneous HFO amplitude and theta phase values such that z_surr_ is parameterized by both time and the offset between the two variables, z_surr_ = A_HG_(t+τ)*e^iϕ^_TH_(t). The modulus of the first moment of z(t), compared to the distribution of moduli for the surrogates, provides a measure of coupling strength. The normalized z-scored length, or Modulation index, is then defined as M_NORM_ = (M_RAW_-µ)/σ, where M_RAW_ is the modulus of the first moment of z(t), µ is the mean of the surrogate lengths, and σ is their standard deviation ^43,45,48^. The Modulation index scores were averaged across all implanted channels for each mouse (∼7.3 channels for each PV-Cre mouse, and 2 channels implanted bilaterally for each C57BL/BJ control mouse), yielding a single score per animal.

To quantify LFP oscillatory power, a sliding window Fourier transform with Hamming window was applied to the LFP signal using MATLAB. Data was analyzed with a 1-second window, 1-second step, and a frequency resolution of 1Hz. Signals were averaged across time windows and frequencies used for cross-frequency phase coupling analysis, and then across all microwires, yielding a single measure per animal.

### Determination of medial prefrontal cortex PV+ interneuron phase locking to local oscillations

Nine days following viral surgery, PV-Cre/VGLUT2-flp mice were again anesthetized with isoflurane (1%), placed in a stereotaxic device, and metal ground screws were secured to the cranium. A 1024-channel silicon probe (NeuroNexus, SINAPS_4S_1024) was implanted to target medial prefrontal cortex (−1.7mm AP in male mice/1.65mm AP in female mice, centered at the midline, measured from bregma; - 4mm DV from the dura). Individual shanks were spaced by 500µm on the probe such that the medial two targeted medial prefrontal cortex bilaterally at ±0.25mm ML. All mice were pair-housed with one animal from the other experimental group. Following a 2-week recovery period, mice were placed in a new cage that contained bedding from their home cage, connected to a mezzanine board/headstage, and this new cage was placed within the recording arena. Mice were individually habituated to this arena during their light cycle, and each mouse was habituated to its own new home cage. After habituation, mice were returned to their pair housing in their original home cage. This habituation procedure was repeated during the dark cycle at least 3 times over the subsequent 2-4 days. On the recording day, mice were habituated to their second home cage for 10 minutes, connected to the connected to a mezzanine board/headstage, and their second home cage was placed within the recording arena. Recordings began after animals were habituated for an additional 10-15 minutes. Neural data was collected for at least 10 minutes. Several mice showed increased activity profiles during this period; thus, neural recordings were extended for 5-10 additional minutes. For each mouse, the last 10 minutes of neural data, which corresponded to low activity periods, were used for subsequent electrophysiological analysis. We selected these periods since our prior work demonstrated that phase coupling is diminished during novelty exposure and exploration^43,91^.

Neural data was sampled at 20kHz using a SmartBox Pro acquisition system (NeuroNexus). To extract single unit activity, neural activity was converted to the *Neurodata Without Borders (NWB)* format, high-pass filtered at 300Hz, and the medial 512 channels (two medial shanks) were automatically sorted with Kilisort4^92^. PV+ single units were identified based on ISI violations < 0.5, presence ratio > 0.9, an amplitude cutoff < 0.1^93,94^, a peak-valley ratio < 1.1, and a mean firing rate > 10Hz^46^. Only single units that mapped to the medial prefrontal cortex (cingulate cortex, prelimbic cortex, infralimbic cortex) targeted channels (192 total PV+ interneurons, 32±5.4 across 6 mice) were used for analysis.

Only neuronal activity that occurred within the last 10 minutes of the recording was used to determine phase locking. This approach was taken to control for small differences in recording length across animals. LFP activity was high-pass filtered at 0.5Hz using a first-order Butterworth filter and then filtered again using a 4^th^-order Butterworth bandpass filter to isolate theta (4-10Hz), gamma (30-80Hz), or high-frequency oscillations (HFOs, 80-200Hz). The instantaneous phase of the filtered LFPs were then determined using the Hilbert transform, and phase locking was calculated using the MATLAB (MathWorks) circular statistics tool box (i.e., the ‘circ_r’ function for the mean resultant length and the ‘circ_rtest’ function for the Rayleigh test of uniformity analyses).

### Social preference and gross locomotor testing in mice subjected to prefrontal cortex microcircuit editing

Eight PV-Cre/VGLUT2-flp mice (12-13 weeks old, balanced across sex) were injected with AAV9-CamKII-Cx34.7_M1_-mEmerald (titer: 1.4×10^13^ vg/ml) and AAV9-hsyn-DIO-Cx35_M1_-T2A-mCherry (titer: 1.3×10^14^ vg/ml), as described above. Seven PV-Cre/VGLUT2-flp control mice were injected with a 1:1 solution of AAV9-CaMKII-eGFP (titer: 2.8×10^13^ vg/ml) and AAV9-hSyn-DIO-mCherry (titer: 1.9×10^13^ vg/ml) to solely express fluorophores. Two weeks after surgical recovery, mice were subjected to behavioral testing.

The social preference assay was conducted as previously reported^95^. In brief, mice were habituated to the experimental room (125 lux) for at least 1 hour prior to behavioral testing. Mice were allowed to explore a rectangular arena (61cm × 42.5cm × 22cm, L×W×H) for 10 minutes. Clear plexiglass walls divided the area into two equal chambers with an opening at the center to allow for free exploration. Each chamber contained a circular holding cage (8.3cm diameter and 12cm tall) containing either a novel object or a C3H target mouse matched for sex and age. Video data was tracked using Ethovision XT17 (Noldus) where interaction time was identified based on proximity (∼5cm) to each chamber. Social preference ratios were calculated as (Social Interaction Time - Object Interaction Time) / (Social Interaction Time + Object Interaction Time). For open field testing, mice were placed in a square arena (45.72cm × 45.72cm × 40.64cm, L×W×H) for 60 minutes. The arena was lit at 50 lux, and the location of the mice was tracked using Ethovision XT17 (Noldus). Several days later, mice were placed back into the area, and their location was tracked for an additional 60 minutes. Data recorded during the first 5 minutes of the first exposure were used to quantify the novel exploratory drive of mice. Data quantified during the second 60-minute exposure were used to quantify gross locomotion function.

### MD single unit responses to direct IL activation

Two mice were injected with AAV9-CamKII-ChR2-EYFP (titer: ≥ 1×10¹³vg/ml) to target IL bilaterally (1.7mm AP, ±0.72mm ML, measured from bregma; - 2.03mm DV from the dura at an angle of 10°). A total of 0.5µL viral solution was delivered at a rate of 100nL/minute. Mice were then implanted bilaterally with two optic fibers (MFC_100/125-0.22_8.0mm_MF2.5_FLT, Doric Lenses) directly above IL (1.7mm AP, ±0.3mm ML, measured from bregma, - 1.75 mm DV, measured from the dura) and with a 1024-channel silicon probe (NeuroNexus, SINAPS_4S_1024) to target MD (−1.58mm AP, centered at 0.13mm ML, measured from bregma; - 4mm DV from the dura). The two optic fibers were constructed onto a single holder such that they were implanted at the same depth.

Following a 2-week recovery period, mice were connected to a mezzanine board/headstage and optic fibers, without anesthesia, and placed in a new cage. Two 473nm lasers (CrystaLaser LC, DL473-025-O, CL-2005 Laser Power Supply and Laser Glow, R47-F-473-nm-DPSS-Laser-System/250mW), calibrated using an optical power meter (ThorLabs PM100D), were used to deliver output at 1mW. Laser stimulation was triggered using analog output from the Cerebus System (Blackrock Microsystems, UT, USA) to deliver 120 light pulses (10ms pulse width), each separated by pseudorandomized inter-stimulus intervals ranging from 8-24 seconds. One mouse was stimulated bilaterally and the other mouse was stimulated unilaterally. Neural data was sampled at 20kHz using a SmartBox Pro acquisition system (NeuroNexus), along with an analog input signal corresponding to the laser trigger. To extract single unit activity, neural activity was converted to *NWB* format, high-pass filtered at 300Hz, and the medial 512 channels (two medial shanks) were automatically sorted with Kilisort4^92^. We also included one lateral shank that was verified histologically to traverse MD in one of the mice. Single units were identified based on ISI violations < 0.5, presence ratio > 0.9, and an amplitude cutoff < 0.1^93,94^ (421/569 total classified cells). Only single units that mapped to the MD-targeted channels (145/421 single units) were used for further analysis.

To determine the response of each MD single unit to IL stimulation, we used our previously described approach^47^. Briefly, neuronal activity relative to the light stimulation was averaged in 20ms bins, shifted by 1ms, and averaged across 120 trials to construct the unit peri-event time histogram. Distributions of the histogram from the [-5000ms, -2000ms] interval were treated as baseline activity. We then determined which 20ms bins, slid in 1ms steps during an epoch spanning from the [0ms, 30ms] interval, met the criteria for modulation by cortical stimulation. A unit was found to be modulated by cortical stimulation if at least 20 bins had firing rates either larger than a threshold of 99% above baseline activity or smaller than a threshold of 94% below baseline activity. This approach was modeled after peri-event analytical approaches utilized in other published studies^96^. To determine the mean LFP evoked responses, neural data recorded from the same channel as each spike was bandpass filtered at 0.5–250Hz and down-sampled to 1000Hz. Activity was then averaged across 120 light pulses. Correlations between peri-stimulus firing rate time histograms and mean LFP evoked responses were calculated for the [0ms, 30ms] interval using a linear regression at α = 0.05.

### Neural data acquisition and analysis in IL-MD circuit-edited mice

Experiments were conducted during two sessions: three weeks and five days after the first viral surgery, and again five weeks after the first viral surgery. The 473nm laser was calibrated to deliver output at 3mW, 1mW, 0.75mW, 0.5mW, and 0.25mW. Laser stimulation was triggered using analog output from the Cerebus System (Blackrock Microsystems, UT, USA). Prior to recordings, mice were connected to a headstage and optic fiber without anesthesia and placed in a new cage.

LFP activity was acquired using a Cerebus recording system, as described above, concurrently with an analog signal corresponding to the laser TTL pulse. After a 10-minute baseline recording period, mice received repeated 10ms pulses of light in IL, delivered with pseudorandomized inter-stimulus intervals ranging between 8 and 24 seconds. During the first session, mice were stimulated with light intensities of 0.25, 0.5, 0.75, and 1mW (30 trials each) in pseudorandomized order. Stimulation protocol was fully automated using a MATLAB script available at https://github.com/carlson-lab/OptoLinCx.

We then determined the portion of mice that showed clear evoked responses at each light threshold. Since our objective was to determine the impact of LinCx expression on IL→MD circuit physiology, our experimental approach required an IL-stimulation intensity that was strong enough to evoke a potential in MD but did not saturate the elicited MD response. When we analyzed data from the first session (as described below), we observed that most animals failed to show an evoked potential in MD greater than 50µV using 0.25, 0.5, or 0.75mW stimulation. Thus, we utilized the 1mW IL stimulation (where 13/21 mice showed mean evoked responses to >50µV; see Supplemental Figure S14), to directly assess the impact of LinCx expression of IL→MD circuit function. We then added 30 trials of 3mW IL stimulation to the second session. Here, we reasoned that mice which failed to show clear evoked responses in MD in responses to 3mW stimulation (>75µW) would be below the detection threshold for determining the impact of LinCx expression at 1mW stimulation. Note that these experimental design criteria were implemented in an initial cohort of eight mice prior to histological confirmation of viral expression and electrode placement. Identical experimental parameters were then utilized for the remainder of the mice in the study.

Custom Python scripts were used to analyze raw recordings (https://github.com/carlson-lab/OptoLinCx). First, the code loads raw NSX files and filters the LFPs with a forward and backward second-order IIR notch filter at 60Hz, with a quality factor of 30, and a Butterworth analog high pass filter of order 5, with a cutoff of 15 Hz using Scipy implementation. After filtering, the average voltage in the -200ms to 1ms pre-stimulus window was subtracted out, normalizing each evoked response to 0mV. Trials were excluded if a given trace had more than 50% of the time points identified as outliers from a 1.5 interquartile range (IQR) between the 25th and 75th quartile. The remaining trials were averaged within each channel and for each stimulus intensity. Channels that showed a pre-stimulus dominant frequency with peaks larger than 25μV were removed from subsequent analysis. Only animals with at least fifteen 1mW trials acquired during both sessions were utilized. Three mice that failed to show a robust response in MD to the 3mW stimulus were removed from subsequent analysis.

The mean of the peak amplitude in the 0-10ms window, averaged across all IL microwires, and in the 10-25ms window, averaged across all MD microwires, was used for comparisons across groups. We reasoned that if an increase in the amplitude of MD-evoked potentials was solely driven by an increase in IL-evoked activity, we would observe a direct correlation between the two variables across animals.

We found no such relationship between the amplitude change of MD- and IL-evoked potentials across the mice that expressed Cx35_M1_ (R = -0.02; P=0.70 using Pearson correlation). Furthermore, we observed a significant stimulation intensity × brain region effect when we compared the magnitude of neural responses across the first session (F_3,60_=34.32, P <5×10^-13^ using a within-subjects two-way ANOVA; see Supplemental Figure S14). Together, these data established that IL and MD responses to light stimulation were not linearly correlated. Thus, our analysis was performed independently for each brain region.

We note that the first neurophysiological testing session was performed 5 days after electrode implantation, which is far shorter than our typical post-operative recovery period of 14 days. While this timeline was necessary to obtain neural recording prior to Cx35_M1_ expression (and thus LinCx formation), this approach of recording neural activity during a period of high post-operative caused concern given that our aim was to quantify the function of a circuit that we knew was impacted by stress exposure. We ultimately concluded that the probative value of measuring the function of the IL→MD circuit in a supraphysiological context (direct optogenetic stimulation of IL), exceeded such concerns. On the other hand, we solely assessed the impact of LinCx expression on the normal physiological function of the IL◊MD after a full postoperative recovery period (14 days). Specifically, the modulation index between IL 2-7Hz and MD 30-70Hz oscillations was calculated for the second session in which we anticipated strong Cx34.7M1 trafficking to MD and strong Cx35M1 expression, as well as full recovery from surgery. Neural activity recorded from a 10-minute baseline period (acquired immediately prior to the light stimulation) was used for analysis. Four to eight LFPs were recorded from each brain region (IL and MD). We then calculated the modulation index for pairs of LFPs recorded from the two regions (6-8 per region), yielding up to 64 measured couplings per mouse. These values were then averaged together, yielding a single score per animal.

### Behavioral testing of IL**◊**MD circuit manipulation

Repeat exposure to a tail suspension test induces behavioral immobility. This behavioral change is specific to the stressful tail suspension test context, as no changes in gross locomotor behavior is observed when mice are also monitored in an open field. For this study, behavioral testing was conducted under low illumination conditions (1-2 lux). Two weeks after the second viral surgery, mice were initially placed in a 17.5in × 17.5in × 11.75in (L×W×H) chamber for five minutes of open field testing. Mice were then suspended 1cm from the tip of their tail for six minutes for the tail suspension assay. Open field and tail suspension behavioral data were acquired during a single testing session, and the entire behavioral testing session was repeated the next day. Testing sessions were video recorded, and open field and tail suspension behavior was analyzed using Ethovision XT 12 (Noldus, Wageniingen, the Netherlands) to quantify immobility in the tail suspension assay, and forward locomotion in the open field. Behavioral experiments and subsequent video analyses were performed blind to group.

### Histology for mouse studies

Mice were perfused transcardially with ice-cold PBS followed by 4% PFA in PBS (EM Sciences, Hatfield, PA). The brains were harvested and coronally sliced in 1X PBS at 35µm using a vibratome (Vibratome Series 3000 Plus, The Vibratome Company, St. Louis, MO) and mounted onto positively charged slides using a mild acetate buffer (82.4mM Sodium Acetate, 17.6mM acetic acid) or 1X PBS solution. Brain slices were covered with DAPI-Mowiol mounting solution [Glycerol, puriss. p.a., Mowiol 4-88 (Sigma-Aldrich, St. Louis, MO), and 0.2M Tris-Cl pH 8.5, DAPI (Sigma-Aldrich, St. Louis, MO)] and cover-slipped prior to imaging. Images were obtained using a Nikon Eclipse fluorescence microscope at 4x magnification with illumination source power and exposure kept consistent between samples.

For the PrL PYR→PV+ study, we only used mice with confirmed bilateral Cx expression for neurophysiological analysis (N=26 out of the 29 brains analyzed).

We also performed immunohistochemistry to assess the specificity of our viral targeting approach for the PrL PYR◊PV+ study. Briefly, we injected three C57BL6/J mice in PrL with AAV-CamKII-Cx34.7_M1_-mEmerald. Following a 4-week expression period, brains were removed, sliced, and stained. We performed immunohistochemistry using anti-PV antibody and AlexaFluor 568 dye. Neurons were identified by DAPI expression and a diameter > 13µm. Green fluorescence (indicating viral expression) and red fluorescence (indicating PV expression) was then quantified across all neurons in PrL. Using this approach, we found that only 4.3% of all cells that expressed Cx34.7_M1_ also expressed PV, and 15.5% of PV+ interneurons expressed Cx35_M1_. Thus, most of the cells that expressed Cx34.7_M1_ were excitatory.

For the IL→MD behavioral study, we used mice that showed bilateral Cx expression in one region, and at least unilateral expression in the other region (N=26 out of the 35 brains analyzed). We chose this strategy since our prior work indicated that unilateral optogenetic stimulation of the IL→MD circuit was sufficient to alter stress-related behavior in the tail suspension assay. For the IL→MD behavioral study, histology was performed as noted above to confirm electrode placement, EYFP/mEmerald trafficking to MD, and mApple expression in MD. In a subset of animals, verification of MD expression required additional staining against mApple. Here, we used Rabbit primary antibody against RFP (Rockland, 600-401-379) and secondary antibody with Alexa Fluor 568 (Abcam, ab175471). ChR2 expression was confirmed via electrophysiology (neural responses to blue light). Only mice with accurate targeting of both viral sites and electrodes were used for analysis.

### Data Visualization

For bar graphs, data is plotted as mean ± s.e.m. Box and whisker plots were created using the MATLAB boxplot function. Here, the central mark is the median, the edges of the box are the 25th and 75th percentiles, the whiskers extend to the most extreme datapoints the algorithm considers to be not outliers, and the outliers are plotted individually as a “+” (MATLAB, The MathWorks, Inc., Natick, MA).

