## Supplemental Materials for "Long-term editing of brain circuits in mice using an engineered electrical synapse"

**Supplemental Table S1, Detailed Author Contributions**

| | |
|---|---|
| Elizabeth Ransey | Conceived FETCH methodology as approach for screening connexin mutant docking; coordinated and performed connexin DNA construct cloning; performed all FETCH experiments, generated flow cytometry data, and co-supervised processing and analysis of FETCH flow-cytometry data. Performed all HEK293FT cell microscopy in the main text and Supplemental Fig. S1; conceived strategy to employ computational models for rational design; coordinated design of computationally guided mutations; designed and cloned AAV vector plasmids; secured funding and resources; prepared figures; wrote original draft of the introduction, mammalian *in vitro* study results and associated materials and methods, and discussion; and revised the paper. |
| Gwenaëlle Thomas | Initiated research on gap junction structural/functional properties across animal species, including target residues to alter docking properties; systematically organized findings, selected candidate proteins as substrates for LinCx, and presented recommendations (to RH and KD) with EA, RB, and HS; performed Cx34.7 expression and trafficking study in mice (viral surgeries and histological analysis); performed viral surgeries for PYR→PV+ mouse editing studies; collected and analyzed neurophysiological data for PYR→PV+ editing study with S.D.M. and K.K.W., performed viral surgeries and collected data for infralimbic cortex→medial dorsal thalamus editing behavioral experiments with KKW; analyzed data for infralimbic cortex→medial dorsal thalamus behavioral experiment; prepared figures. |
| Elias Wisdom | Conceived *C elegans* neurophysiological and behavioral experiments with AAP; Generated Cx34.7 and Cx35 wild-type and mutant expression vectors for *C. elegans*; performed *C. elegans* experiments testing wild-type and mutant Cx34.7 and Cx35 under homotypic and heterotypic expression conditions and analyzed results; prepared figures; wrote original draft of *in vivo* study results and associated methods with A.A.P. and D.C.; and revised the paper. |
| Agustin Almoril-Porras | Conceived *C elegans* neurophysiological and behavioral experiments with EW; generated *C elegans strains*; generated Cx36 and Cx43 expression vectors for *C. elegans;* performed *C. elegans* experiments testing mutant Cx34.7 and Cx35 against Cx36 and Cx43. under heterotypic expression conditions and analyzed results; assisted in calcium imaging analyses; coordinated all C. elegans experiments; prepared figures; wrote original draft of *in vivo* study results and associated methods with E.W. and D.C.R.; and revised the paper. |
| Ryan Bowman | Researched gap junction structural/functional properties across animal species, systematically organized findings, selected candidate proteins as substrates for LinCx, and presented recommendations (to RH and KD) with EA, GET, and HS; Cloned connexin DNA constructs; Tested in vitro methods for evaluating gap function in mammalian cells. |
| Elise Adamson | Researched gap junction structural/functional properties across animal species, systematically organized findings, selected candidate proteins as substrates for LinCx, and presented recommendations (to RH and KD) with GET, HS, and RB; Cloned connexin DNA constructs; |
| Kathryn Katsue Walder-Christensen | Planned PYR→PV+ and IL→MD (behavioral) experiments with G.E.T., S.D.M., K.D., and performed associated mouse viral surgeries in conjunction with S.D.M. and G.E.T.; Assisted in PYR→PV+ behavioral data collection and analysis; |

| | |
|---|---|
| | collected and analyzed behavioral data for mouse IL→MD editing behavioral experiment with GET; Performed TST control experiment in unedited mice; performed mouse histological analysis with K.C. |
| Jesse A. White | Performed the electrophysiological experiments with HEK293FT and HEK293FT Cx43 and Cx35 double knockout cells, analyzed the results, and prepared first draft of results and methods. |
| Dalton N. Hughes | Collected and analyzed C57 neurophysiological control data utilized for mouse PYR→PV+ editing experiment. |
| Hannah Schwennesen | Researched gap junction structural/functional properties across animal species, systematically organized findings, selected candidate proteins as substrates for LinCx, and presented recommendations (to RH and KD) with EA, GET, and RB; |
| Caly Ferguson | Performed all reanalysis of computational models used to assess connexin hemichannel interactions. |
| Kay M. Tye | Supervised the electrophysiological experiments with HEK293FT and HEK293FT Cx43 and Cx35 double knockout cells. |
| Stephen D. Mague | Performed all PYR→PV+ and IL→MD (behavioral) mouse viral surgeries in conjunction with K.K.W. and G.E.T.; Performed electrode implantation surgeries in mice with KD. Oversaw mouse PYR→PV+ editing experiment. Supervised mouse IL→MD editing experiments. |
| Longgang Niu | Performed the electrophysiological experiments with *Xenopus* oocytes, analyzed the results, and made the figures. |
| Zhao-Wen Wang | Supervised the electrophysiological experiments with *Xenopus* oocytes, and wrote the related text (results, methods, and figure legends). |
| Daniel Colón-Ramos | Conceived *C. elegans* based strategy to assess designer gap junction functional properties; supervised all *C. elegans* experiments; wrote original draft of *in vivo* study results and associated methods with E.W. and A.A.P.; and revised the paper. |
| Rainbo Hultman | Coordinated research of gap junction structural/functional properties across animal species; co-selected fish Cx34.7/Cx35 pair for subsequent protein engineering with KD; Designed Cx34.7/Cx35 mutant library for subsequent cloning; Co-conceptualized framework for screening homotypic then heterotypic interactions in Cx mutants with KD; secured funding and resources; revised the paper. |
| Nenad Bursac | Supervised in vitro FETCH studies with KD; secured funding; wrote original draft with E.R., K.C., A.A.P., E.W., D.C.R., and K.D.; and revised the paper. |
| Kafui Dzirasa | Conceived strategy to integrate membrane of two distinct cell types using heterotypically, but not homotypically docking gap junction hemichannels. Co-selected Cx34.7/Cx35 pair for subsequent protein engineering with RCH; Co-conceptualized framework for screening homotypic then heterotypic interactions in Cx mutants with RCH; supervised in vitro FETCH studies with NB; supervised computational studies; analyzed data and oversaw all statistical procedures; conceived all mouse editing experiments; performed electrode implantation surgeries for PYR→PV+ editing experiment and controls with S.D.M. and K.K.W; supervised PYR→PV+ editing experiment and performed all data analysist; Designed experiment testing $Cx35_{WT.}$ Designed and performed circuit interrogation study using optogenetics; analyzed all neurophysiological data, and performed behavioral analysis with GET; Designed and secured |

| | |
|---|---|
| | funding and resources; prepared figures; wrote original draft with E.R., A.A.P., E.W., D.C.R., and N.B.; and revised the paper. |
| **Attribution Process | Each team member outlined their individual contributions across a standard set of domains (conceptualization and methodology, formal analysis, investigation, resources, writing -original draft, writing -review & editing, visualization, Supervision, and Project Administration and Funding Acquisition), and subsequently had the opportunity to edit a summarized attribution description to their satisfaction. Contribution summaries were then shared across all team members. Each team member had the opportunity to raise concerns with regards to any other team member's outlined contributions, and issues that remained unaddressed after additional revisions were subjected to a mediation process led by the lead principal investigator. The assigned authorships and these detailed author contribution descriptions on the initial submitted manuscript deposited in a preprint server reflect the outcome of this process[1]. During revisions, contributions were made by a subset of the authors and several new authors.<br><br>All new attributions were signed off by all authors, and the authorship assignments were revised by the lead principal investigator to reflect each author's composite contributions. Each team member again had the opportunity to raise concerns with regards to any other team member's outlined contributions. Additionally, a subset of the data was reanalyzed to ensure the rigor and reproducibility of the work presented here. The development of all the computational methodology utilized to perform this reanalysis have been acknowledged in a prior preprint [1]. The author list was revised to reflect this process. |

**Supplemental Table S2, *C. elegans* Strain Table**

| | Genotype | Source | Line # |
|---|---|---|---|
| N2 | *Wild-type* | CGC | |
| DCR3056 | *olaIs17 [Pmod-1::GCaMP6s (25ng/ul) Pttx-3::mCherry (25ng/ul) Punc-122::dsRed (40ng/ul)] I* | Hawk et al. 2018 | |
| DCR6604 | *olaIs23 [Pgcy-8(800)::caPKC-1B (30ng/ul), Pgcy-8(800)::tagRFP (10ng/ul), Punc-122::RFP (30ng/ul)] V; wyIs629 [Pgcy-8(2kb)::GCaMP6s (30ng/ul), Pgcy-8(2kb)::mCherry (5ng/ul), Punc-122:GFP (20ng/ul)] X* | Hawk et al. 2018 | |
| DCR5790 | *olaIs17 [Pmod-1::GCaMP6s (25ng/ul) Pttx-3::mCherry (25ng/ul) Punc-122::dsRed (40ng/ul)] I; olaIs72 [Pelt-7::mCherry (25ng/ul) + Pttx-3::CX36::mCherry (25ng/ul)]* | Hawk et al. 2018 | |
| DCR5793 | *olaIs17 [Pmod-1::GCaMP6s (25ng/ul) Pttx-3::mCherry (25ng/ul) Punc-122::dsRed (40ng/ul)] I; olaIs70 [Pelt-7::GFP (15ng/ul) + Pgcy-8::CX36::mCherry (25ng/ul)]; olaIs72 [Pelt-7::mCherry (25ng/ul) + Pttx-3::CX36::mCherry (25ng/ul)]* | Hawk et al. 2018 | |
| DCR5404 | *olaIs17 [Pmod-1::GCaMP6s (25ng/ul) Pttx-3::mCherry (25ng/ul) Punc-122::dsRed (40ng/ul)] I; olaIs23 [Pgcy-8(800)::caPKC-1B (30ng/ul), Pgcy-8(800)::tagRFP (10ng/ul), Punc-122::RFP (30ng/ul)] V; olaEx3219 [Pgcy-8::CX36::mCherry (25ng/ul); Pttx-3::CX36::mCherry (25ng/ul); Pmyo-3::Red(10ng/ul)]* | Hawk et al. 2018 | |
| DCR8225 | *olaIs17 [Pmod-1::GCaMP6s (25ng/ul) Pttx-3::mCherry (25ng/ul) Punc-122::dsRed (40ng/ul)] I; olaIs23 [Pgcy-8(800)::caPKC-1B (30ng/ul), Pgcy-8(800)::tagRFP (10ng/ul), Punc-122::RFP (30ng/ul)] V 13xOC* | This Paper | |
| DCR8678 | *olaEx5223 [Pgcy-8::CX34.7::GFP; Pttx-3::CX34.7::mCherry; Punc-122::GFP (All 25ng/ul)]* | This Paper | Line 1 |
| DCR8717 | *olaEx5255 [Pgcy-8::CX34.7::GFP; Pttx-3::CX34.7::mCherry; Punc-122::GFP (All 25ng/ul)]* | This Paper | Line 1 |
| DCR8716 | *olaEx5254 [Pgcy-8::CX34.7(E214K, E223K)::GFP; Pttx-3::CX34.7(E214K, E223K)::mCherry; Punc-122::GFP (All 25ng/ul)]* | This Paper | Line 1 |
| DCR8673 | *olaEx5218 [Pgcy-8::CX35(K221E)::GFP; Pttx-3::CX35(K221E)::mCherry; Punc-122::GFP (All 25ng/ul)]* | This Paper | Line 1 |
| DCR8669 | *olaEx5214 [Pgcy-8::CX34.7(E214K, E223K)::GFP; Pttx-3::CX35(K221E)::mCherry; Punc-122::GFP (All 25ng/ul)]* | This Paper | Line 1 |
| DCR8684 | *olaEx5230 [Pgcy-8::CX35::GFP; Pttx-3::CX35::mCherry; Punc-122::GFP (All 25ng/ul)]* | This Paper | Line 2 |
| DCR8719 | *olaEx5256 [Pgcy-8::CX35::GFP; Pttx-3::CX35::mCherry; Punc-122::GFP (All 25ng/ul)]* | This Paper | Line 2 |
| DCR8715 | *olaEx5253 [Pgcy-8::CX34.7(E214K, E223K)::GFP; Pttx-3::CX34.7(E214K, E223K)::mCherry; Punc-122::GFP (All 25ng/ul)]* | This Paper | Line 2 |
| DCR8674 | *olaEx5219 [Pgcy-8::CX35(K221E)::GFP; Pttx-3::CX35(K221E)::mCherry; Punc-122::GFP (All 25ng/ul)]* | This Paper | Line 2 |

| | | | |
|---|---|---|---|
| DCR8676 | *olaEx5221 [Pgcy-8::CX34.7(E214K, E223K)::GFP; Pttx-3::CX35(K221E)::mCherry; Punc-122::GFP (All 25ng/ul)]* | This Paper | Line 2 |
| DCR8685 | *olaEx5231 [Pgcy-8::CX34.7::GFP; Pttx-3::CX34.7::mCherry; Punc-122::GFP (All 25ng/ul)]* | This Paper | Line 3 |
| DCR8720 | *olaEx5257 [Pgcy-8::CX35::GFP; Pttx-3::CX35::mCherry; Punc-122::GFP (All 25ng/ul)]* | This Paper | Line 3 |
| DCR8714 | *olaEx5252 [Pgcy-8::CX34.7(E214K, E223K)::GFP; Pttx-3::CX34.7(E214K, E223K)::mCherry; Punc-122::GFP (All 25ng/ul)]* | This Paper | Line 3 |
| DCR8672 | *olaEx5217 [Pgcy-8::CX35(K221E)::GFP; Pttx-3::CX35(K221E)::mCherry; Punc-122::GFP (All 25ng/ul)]* | This Paper | Line 3 |
| DCR8677 | *olaEx5222 [Pgcy-8::CX34.7(E214K, E223K)::GFP; Pttx-3::CX35(K221E)::mCherry; Punc-122::GFP (All 25ng/ul)]* | This Paper | Line 3 |
| DCR8675 | *olaIs17 [Pmod-1::GCaMP6s (25ng/ul) Pttx-3::mCherry (25ng/ul) Punc-122::dsRed (40ng/ul)] I; olaIs23 [Pgcy-8(800)::caPKC-1B (30ng/ul), Pgcy-8(800)::tagRFP (10ng/ul), Punc-122::RFP (30ng/ul)] V; olaEx5220 [Pgcy-8::CX34.7::GFP; Pttx-3::CX34.7::mCherry; Pelt-7::NLS::mCherry (All 25ng/ul)]* | This Paper | Line 1 |
| DCR8776 | *olaIs17 [Pmod-1::GCaMP6s (25ng/ul) Pttx-3::mCherry (25ng/ul) Punc-122::dsRed (40ng/ul)] I; olaIs23 [Pgcy-8(800)::caPKC-1B (30ng/ul), Pgcy-8(800)::tagRFP (10ng/ul), Punc-122::RFP (30ng/ul)] V; olaEx5287 [Pgcy-8::CX35::GFP; Pttx-3::CX35::mCherry; Punc-122::GFP (All 25ng/ul)]* | This Paper | Line 1 |
| DCR8777 | *olaIs17 [Pmod-1::GCaMP6s (25ng/ul) Pttx-3::mCherry (25ng/ul) Punc-122::dsRed (40ng/ul)] I; olaIs23 [Pgcy-8(800)::caPKC-1B (30ng/ul), Pgcy-8(800)::tagRFP (10ng/ul), Punc-122::RFP (30ng/ul)] V; olaEx5288 [Pgcy-8::CX34.7(E214K, E223K)::GFP; Pttx-3::CX34.7(E214K, E223K)::mCherry; Punc-122::GFP (All 25ng/ul)]* | This Paper | Line 1 |
| DCR8671 | *olaIs17 [Pmod-1::GCaMP6s (25ng/ul) Pttx-3::mCherry (25ng/ul) Punc-122::dsRed (40ng/ul)] I; olaIs23 [Pgcy-8(800)::caPKC-1B (30ng/ul), Pgcy-8(800)::tagRFP (10ng/ul), Punc-122::RFP (30ng/ul)] V; olaEx5216 [Pgcy-8::CX35(K221E)::GFP; Pttx-3::CX35(K221E)::mCherry; Pelt-7::NLS::mCherry (All 25ng/ul)]* | This Paper | Line 1 |
| DCR8670 | *olaIs17 [Pmod-1::GCaMP6s (25ng/ul) Pttx-3::mCherry (25ng/ul) Punc-122::dsRed (40ng/ul)] I; olaIs23 [Pgcy-8(800)::caPKC-1B (30ng/ul), Pgcy-8(800)::tagRFP (10ng/ul), Punc-122::RFP (30ng/ul)] V; olaEx5215 [Pgcy-8::CX34.7(E214K, E223K)::GFP; Pttx-3::CX35(K221E)::mCherry; Pelt-7::NLS::mCherry (All 25ng/ul)]* | This Paper | Line 1 |
| DCR9178 | *olaEx5452 [Pgcy-8::CX34.7(E214K, E223K)::GFP; Pttx-3::CX43::mCherry; Pelt-7::NLS::mCherry (All 25ng/ul)]* | This Paper | Line 1 |
| DCR9179 | *olaEx5453 [Pgcy-8::CX34.7(E214K, E223K)::GFP; Pttx-3::CX43::mCherry; Pelt-7::NLS::mCherry (All 25ng/ul)]* | This Paper | Line 2 |

| | | | |
|---|---|---|---|
| DCR9180 | *olaEx5454 [Pgcy-8::CX34.7(E214K, E223K)::GFP; Pttx-3::CX43::mCherry; Pelt-7::NLS::mCherry (All 25ng/ul)]* | This Paper | Line 3 |
| DCR9181 | *olaIs17 [Pmod-1::GCaMP6s (25ng/ul) Pttx-3::mCherry (25ng/ul) Punc-122::dsRed (40ng/ul)] I; olaIs23 [Pgcy-8(800)::caPKC-1B (30ng/ul), Pgcy-8(800)::tagRFP (10ng/ul), Punc-122::RFP (30ng/ul)] V; olaEx5455 [Pgcy-8::CX34.7(E214K, E223K)::GFP; Pttx-3::CX43::mCherry; Pelt-7::NLS::mCherry (All 25ng/ul)]* | This Paper | Line 1 |
| DCR9182 | *olaEx5456 [Pgcy-8::CX43::GFP; Pttx-3::CX35(K221E)::mCherry; Pelt-7::NLS::mCherry (All 25ng/ul)]* | This Paper | Line 1 |
| DCR9183 | *olaIs17 [Pmod-1::GCaMP6s (25ng/ul) Pttx-3::mCherry (25ng/ul) Punc-122::dsRed (40ng/ul)] I; olaIs23 [Pgcy-8(800)::caPKC-1B (30ng/ul), Pgcy-8(800)::tagRFP (10ng/ul), Punc-122::RFP (30ng/ul)] V; olaEx5457 [Pgcy-8::CX43::GFP; Pttx-3::CX35(K221E)::mCherry; Pelt-7::NLS::mCherry (All 25ng/ul)]* | This Paper | Line 1 |
| DCR9184 | *olaIs17 [Pmod-1::GCaMP6s (25ng/ul) Pttx-3::mCherry (25ng/ul) Punc-122::dsRed (40ng/ul)] I; olaIs72 [Pelt-7::mCherry (25ng/ul) + Pttx-3::CX36::mCherry (25ng/ul)]; olaEx5458 [Pgcy-8::CX34.7(E214K, E223K)::GFP; Pelt-7::NLS::GFP (All 25ng/ul)]* | This Paper | Line 1 |
| DCR9185 | *olaIs17 [Pmod-1::GCaMP6s (25ng/ul) Pttx-3::mCherry (25ng/ul) Punc-122::dsRed (40ng/ul)] I; olaIs72 [Pelt-7::mCherry (25ng/ul) + Pttx-3::CX36::mCherry (25ng/ul)]; olaEx5459 [Pgcy-8::CX34.7(E214K, E223K)::GFP; Pelt-7::NLS::GFP (All 25ng/ul)]* | This Paper | Line 2 |
| DCR9186 | *olaIs17 [Pmod-1::GCaMP6s (25ng/ul) Pttx-3::mCherry (25ng/ul) Punc-122::dsRed (40ng/ul)] I; olaIs72 [Pelt-7::mCherry (25ng/ul) + Pttx-3::CX36::mCherry (25ng/ul)]; olaEx5460 [Pgcy-8::CX34.7(E214K, E223K)::GFP; Pelt-7::NLS::GFP (All 25ng/ul)]* | This Paper | Line 3 |
| DCR7790 | *olaIs17 [Pmod-1::GCaMP6s (25ng/ul) Pttx-3::mCherry (25ng/ul) Punc-122::dsRed (40ng/ul)] I; olaIs70 [Pelt-7::GFP (15ng/ul) + Pgcy-8::CX36::mCherry (25ng/ul)]* | This Paper | |
| DCR9187 | *olaIs17 [Pmod-1::GCaMP6s (25ng/ul) Pttx-3::mCherry (25ng/ul) Punc-122::dsRed (40ng/ul)] I; olaIs70 [Pelt-7::GFP (15ng/ul) + Pgcy-8::CX36::mCherry (25ng/ul)]; olaEx5461 [Pttx-3::CX35(K221E)::mCherry; Pelt-7::NLS::mCherry (All 25ng/ul)]* | This Paper | Line 1 |
| DCR9188 | *olaIs17 [Pmod-1::GCaMP6s (25ng/ul) Pttx-3::mCherry (25ng/ul) Punc-122::dsRed (40ng/ul)] I; olaIs70 [Pelt-7::GFP (15ng/ul) + Pgcy-8::CX36::mCherry (25ng/ul)]; olaEx5462 [Pttx-3::CX35(K221E)::mCherry; Pelt-7::NLS::mCherry (All 25ng/ul)]* | This Paper | Line 2 |

| | | | |
|---|---|---|---|
| DCR9189 | *olaIs17 [Pmod-1::GCaMP6s (25ng/ul) Pttx-3::mCherry (25ng/ul) Punc-122::dsRed (40ng/ul)] I; olaIs70 [Pelt-7::GFP (15ng/ul) + Pgcy-8::CX36::mCherry (25ng/ul)]; olaEx5463 [Pttx-3::CX35(K221E)::mCherry; Pelt-7::NLS::mCherry (All 25ng/ul)]* | This Paper | Line 3 |

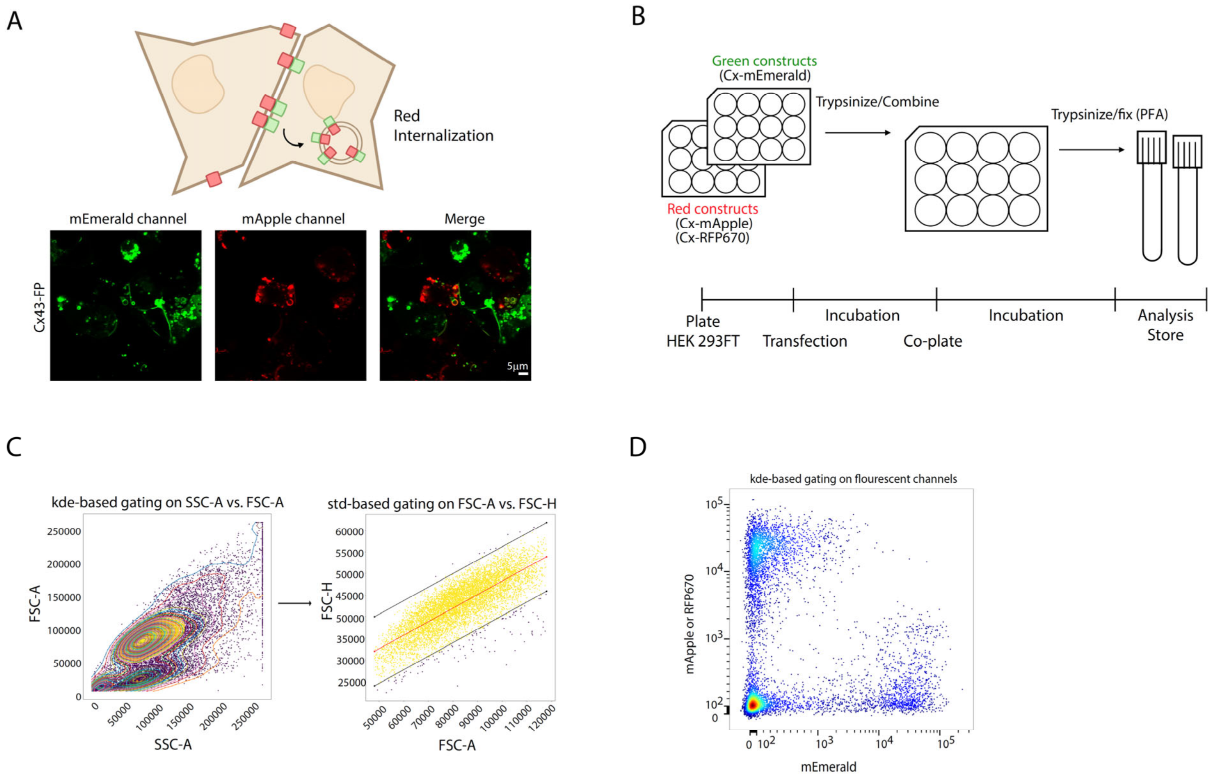


**Supplemental Figure S1: Schematic for Flow-Enabled Tracking of Connexosomes in HEK Cells (FETCH). A)** Depiction of red fluorescence internalization following gap junction formation (top). Image showing co-plated population of cells expressing counterpart Cx43 tagged to mApple or mEmerald. Image corresponds to 42 Cx43/Cx43 samples subjected to the initial FETCH analysis. **B)** FETCH pipeline **C)** Automated gating pipeline that uses sequential kernel density estimate (kde)- and standard deviation-based gating approaches **D)** Flow cytometry plot obtained from automated pipeline for a well containing one connexin pair.

### Cx36 Extracellular Loop 2

| Species | Start | Sequence | End |
|---|---|---|---|
| Homo sapiens | 227– | GLYECNRYPCIKEVECYVSRPTEKTVFLVF | –256 |
| Mus musculus | 227– | GLYECNRYPCIKEVECYVSRPTEKTVFLVF | –256 |
| Macaca mulatta | 227– | GLYECNRYPCIKEVECYVSRPTEKTVFLVF | –256 |
| Callithrix jacchus | 289– | GLYECNRYPCIKEVECYVSRPTEKTVFLVF | –318 |
| Taeniopygia guttata | 196– | AIFECDRYPCVKEVECYVSRPTEKSVFLVF | –225 |
| Danio rerio (Cx34.7) | 209– | GIFECDRYPCLKEVECYVSRPTEKTVFLVF | –238 |
| Danio rerio (Cx35) | 210– | AVYECDRYPCIKDVECYVSRPTEKTVFLVF | –239 |

### Cx43 Extracellular Loop 2

| Species | Start | Sequence | End |
|---|---|---|---|
| Homo sapiens | 171– | LIQWYIYGFSLSAVYTCKRDPCPHQVDCFLSRPTEK | –206 |
| Mus musculus | 171– | LIQWYIYGFSLSAVYTCKRDPCPHQVDCFLSRPTEK | –206 |
| Macaca mulatta | 171– | LIQWYIYGFSLSAVYTCKRDPCPHQVDCFLSRPTEK | –206 |
| Callithrix jacchus | 171– | LIQWYIYGFSLSAVYTCKRDPCPHQVDCFLSRPTEK | –206 |
| Taeniopygia guttata | 171– | LIQWYIYGFSLNAIYTCERDPCPHRVDCFLSRPTEK | –206 |
| Danio rerio | 171– | VIQWYLYGFSLSAVYTCERTPCPHRVDCFLSRPTEK | –206 |

### Cx45 Extracellular Loop 2

| Species | Start | Sequence | End |
|---|---|---|---|
| Homo sapiens | 200– | GFQVHPFYVCSRLPCPHKIDCFI | –222 |
| Mus musculus | 200– | GFQVHPFYVCSRLPCPHKIDCFI | –222 |
| Macaca mulatta | 200– | GFQVHPFYVCSRLPCPHKIDCFI | –222 |
| Callithrix jacchus | 200– | GFQVHPFYVCSRLPCPHKIDCFI | –222 |
| Taeniopygia guttata | 198– | RFEVSPSYVCSRSPCPHTVDCFV | –220 |
| Danio rerio | 196– | GFEVAPSYVCTRSPCPHTVDCFV | –218 |

**Supplemental Figure S2: Sequence alignment of several connexin proteins predicted extracellular loop 2.** Predicted EL2 regions of Cx36 (GJD2), Cx43 (GJA1), and Cx45 (GJC1) for humans and several species broadly utilized in neuroscience research, related to Fig. 6. Identical residues are shown in black, residues that are variable are highlighted with red text. Residues of Cx36 that align to the interaction motif of Cx34.7 and Cx35 are indicated by blue underline. Note zebrafish (Danio rerio) do not have a Cx36 gene, thus the closest homolog, Cx34.7, was used for comparison.

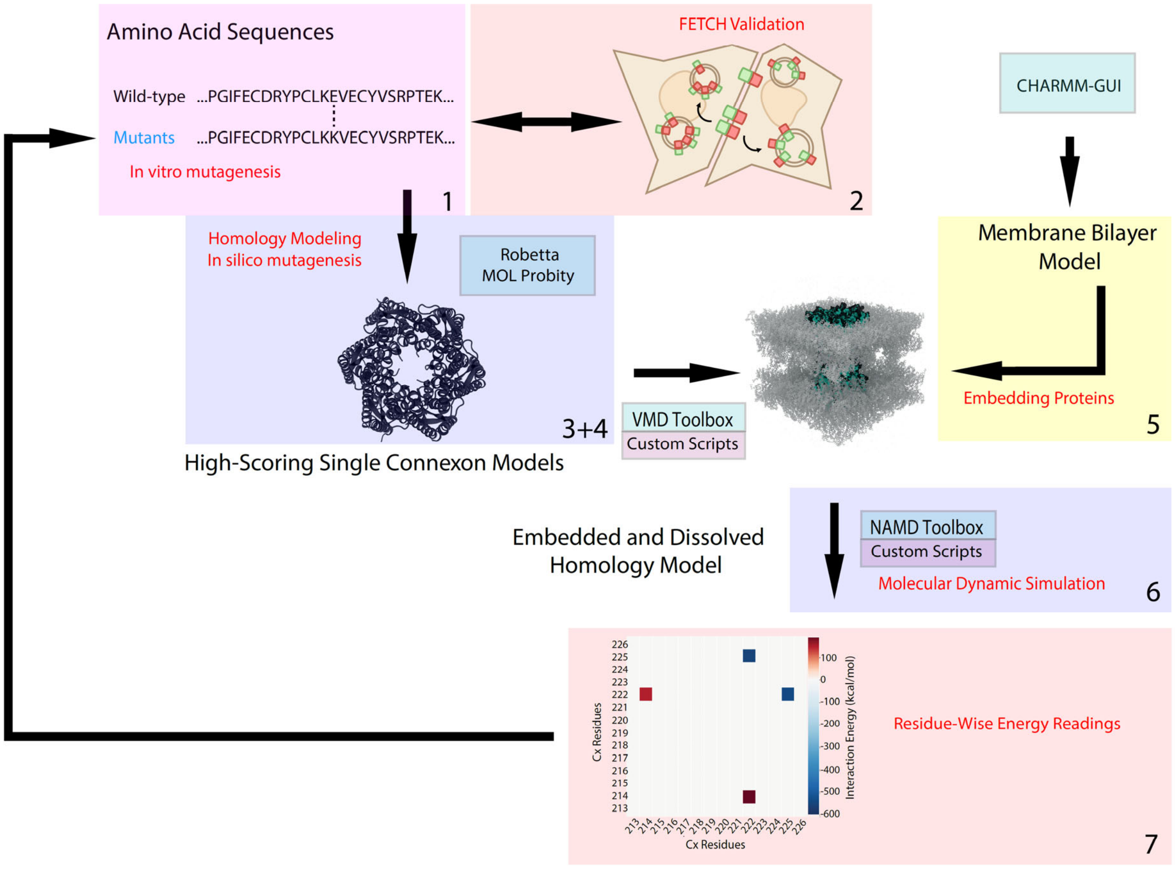


**Supplemental Figure S3: Integrated approach used to engineer Cx34.7 and Cx35 mutants with docking selectively.** Our approach consists of seven integrated components: 1) *in vitro* protein mutagenesis, 2) FETCH screening/validation, 3) homology model generation, 4) *in silico* protein mutagenesis, 5) embedding of proteins in a lipid bilayer and aqueous solution, 6) system minimization, equilibration, and molecular dynamics simulation, 7) and residue-wise energy calculation.

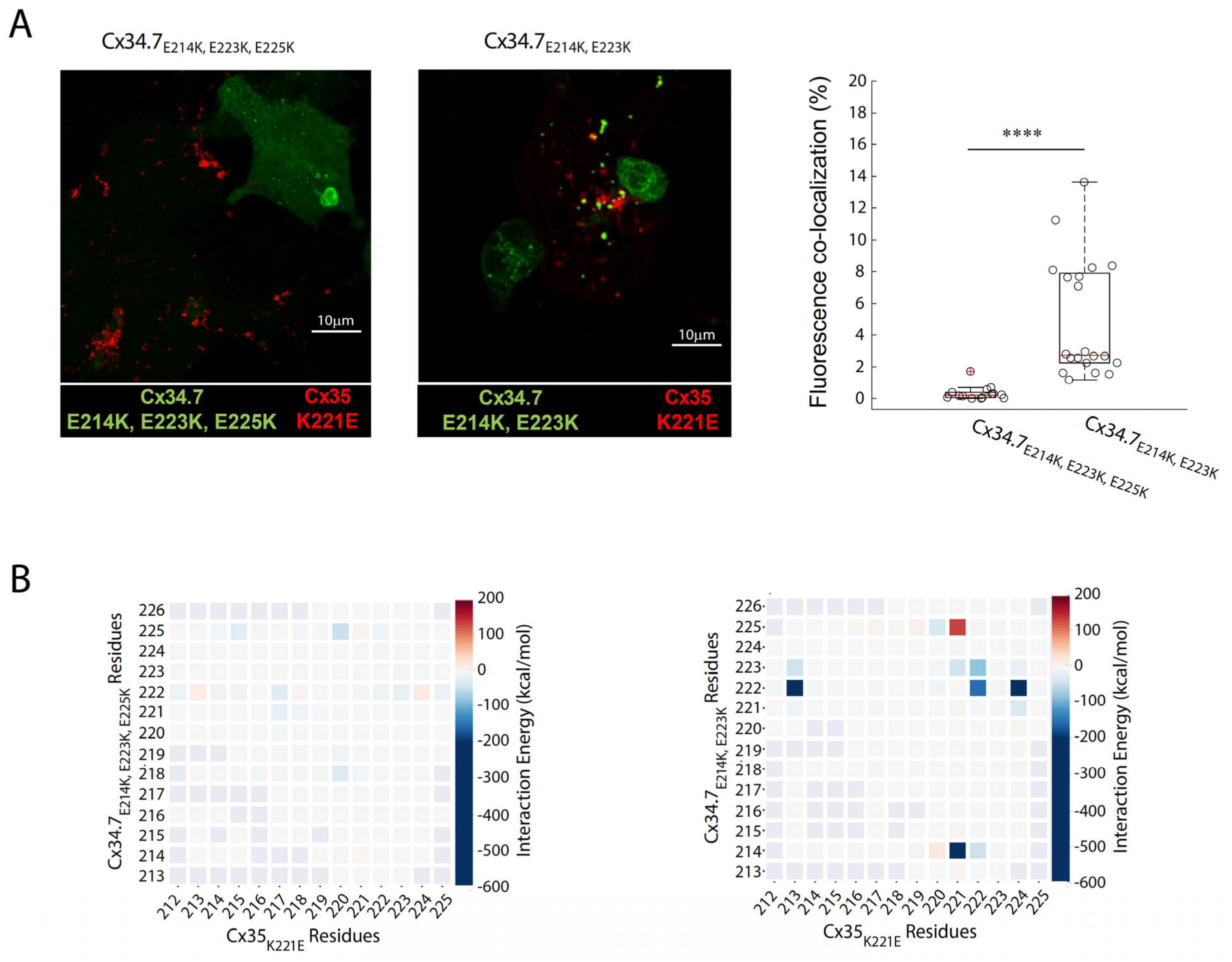


**Supplemental Figure S4: Cx34.7$_{E214K/E223K/E225K}$ shows disrupted docking. A)** Fifteen images were obtained of cell doublets of Cx34.7$_{E214K/E223K/E225K}$ mutants (green) or Cx34.7$_{E214K/E223K/E225K}$ mutants adjacent to Cx35 $_{K221E}$ mutants (red). Each image obtained was a 5.5-micron Z-stack with 12 steps, which was then Z-projected to obtain a 3-D image for analysis. The Colocalization Threshold program on FIJI (ImageJ) was used to determine the percent of Cx34.7$_{E214K/E223K/E225K}$ or Cx34.7$_{E214K/E223K}$ pixels above the threshold that were colocalized to Cx35$_{K221E}$ pixels. We observed a greater percentage of colocalized green and red pixels for the Cx34.7$_{E214K/E223K}$/Cx35$_{K221E}$ pairs than the Cx34.7$_{E214K/E223K/E225K}$/Cx35$_{K221E}$ pairs, indicating a higher rate of putative gap junction formation ($U=109$, $P<10^{-5}$, for comparison across the two groups using a Mann-Whitney U-test). ***$p < 0.0001$. Images were obtained on the Zeiss 800 Confocal Microscope at North Carolina Central University Biomanufacturing Research Institute and Technology Enterprise facility. **B)** Contact plots for EL2-to-EL2 interactions produced by molecular dynamics simulation for Cx34.7$_{E214K/E223K/E225K}$/Cx35$_{K221E}$ (left) and Cx34.7$_{E214K/E223K}$/Cx35$_{K221E}$ (right, similar to Figure 2F).

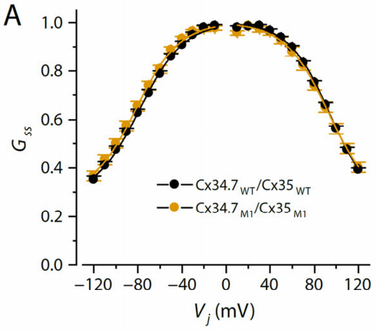


| | $V_0$ (mV) | | $G_{jmin}$ | | $A$ | |
|---|---|---|---|---|---|---|
| | Left | Right | Left | Right | Left | Right |
| $Cx34.7_{WT}/Cx35_{WT}$ | -84.47 ± 2.17 | 84.71 ± 2.88 | 0.28 ± 0.009 | 0.27 ± 0.022 | -0.057 ± 0.001 | 0.051 ± 0.002 ## |
| $Cx34.7_{M1}/Cx35_{M1}$ | -86.69 ± 2.01 | 87.22 ± 2.84 | 0.28 ± 0.031 | 0.28 ± 0.059 | -0.056 ± 0.002 | 0.050 ± 0.002 # |

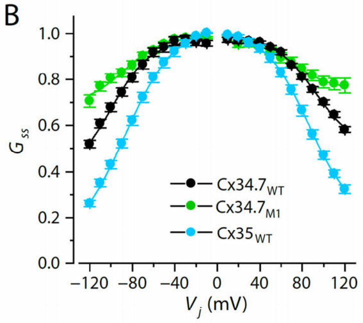


| | $V_0$ (mV) | | $G_{jmin}$ | | $A$ | |
|---|---|---|---|---|---|---|
| | Left | Right | Left | Right | Left | Right |
| $Cx34.7_{WT}$ | -102.33±3.47 | 100.65±2.79 | 0.29±0.035 | 0.39±0.036 | -0.048±0.003 | 0.043±0.003 |
| $Cx34.7_{M1}$ | -95.34± 6.67 | 85.38±7.73 | 0.21±0.003* | 0.64±0.059*# | -0.043±0.004 | 0.047±0.008 |
| $Cx35_{WT}$ | -82.02±2.54* | 81.63±1.94* | 0.19±0.012* | 0.24±0.014*# | -0.054±0.002 | 0.056±0.003* |

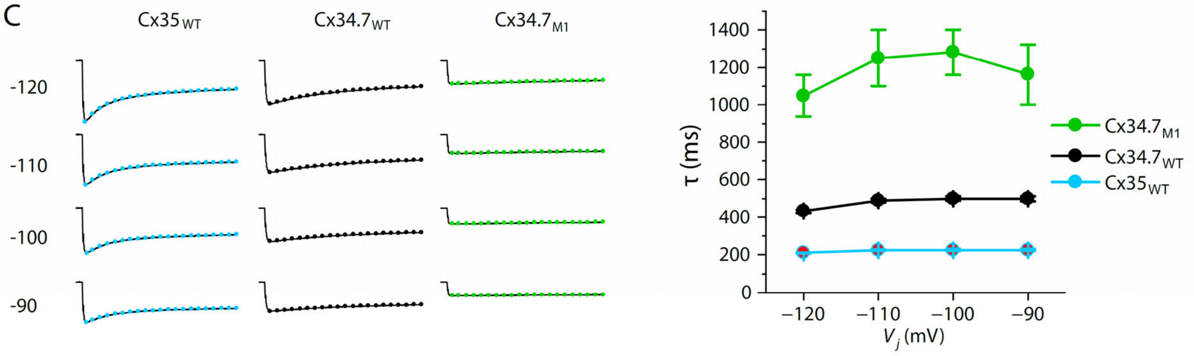


**Supplemental Figure S5: Biophysical properties of wild type and mutant Cx34.7 and Cx35 hemichannels. A)** Relationships between steady-state junctional conductance ($G_{ss}$) and transjunctional voltage ($V_{j,}$, left), and comparison of $V_0$, $G_{jmin}$, $A$, and the left and right sides of the $G_{ss} – V_j$ curves for $Cx34.7_{WT}/Cx35_{WT}$ vs $Cx34.7_{M1}/Cx35_{M1}$ junctions. These variables were fitted to the Boltzmann equation where $V_0$ is the voltage at which the conductance is half-maximal, $G_{min}$ is the normalized voltage-insensitive residual conductance and $A$ is a parameter defining the steepness of voltage sensitivity[2]. The $G_{ss} – V_j$ plot was based on averaged $Ij_{ss}$ recorded from both oocytes of the pair and using the $V_j$ defined by "$V_m$ of $Cx34.7_{WT}$ (or $Cx34.7_{M1}$) – $V_m$ of $Cx35_{WT}$ (or $Cx35_{M1}$)" as the *x*-axis. The pound symbols "#" and "##" indicate significant differences between the left and right $G_{ss} – V_j$ curves at $p < 0.05$ and $p < 0.01$, respectively (paired *t*-test). No significant difference was detected between the two groups. N=6 for all groups. **B)** Relationships between $G_{ss}$ and $V_j$, and comparison *of* $V_0$, $G_{jmin}$, and $A$ values among the different gap junctions and between the left and right sides of the $G_{ss} – V_j$ curves. The asterisk (*) indicates a significant difference compared with $Cx34.7_{WT}$ ($p < 0.01$, one-way ANOVA followed by Tukey's post-hoc test), while the pound symbol (#) indicates a significant difference between the left and right side of the same $G_{ss} - V_j$ curve ($p < 0.01$, paired *t*-test). **C)** Representative $I_j$ traces (solid lines) fitted by a single exponential (dotted lines), and comparison of the deactivation time constant (τ) among the different groups. N=6 for all groups.

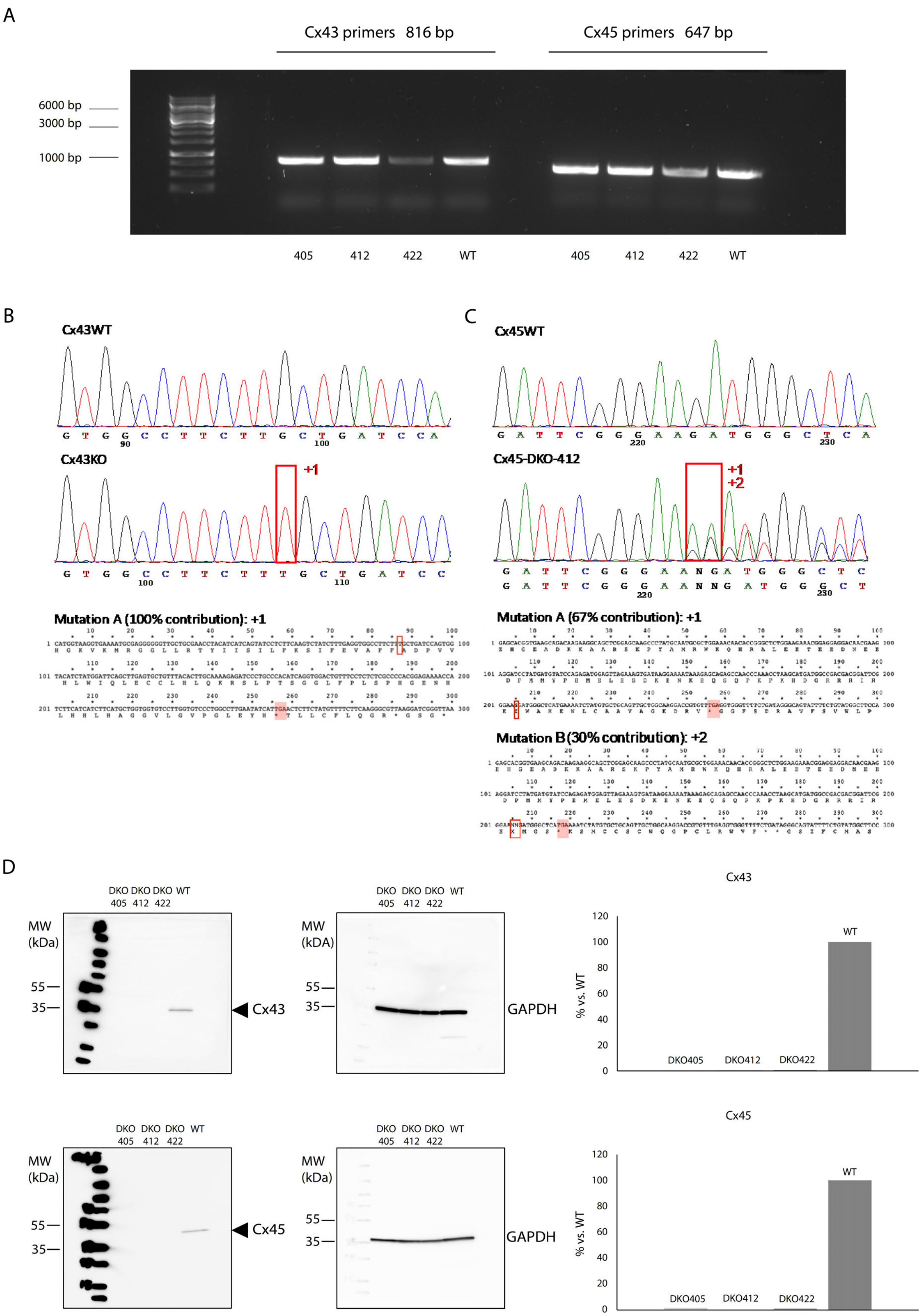
A
Cx43 primers 816 bp
Cx45 primers 647 bp
6000 bp
3000 bp
1000 bp
405
412
422
WT
405
412
422
WT
B
Cx43WT
Cx43KO
+1
Mutation A (100% contribution): +1
C
Cx45WT
Cx45-DKO-412
+1
+2
Mutation A (67% contribution): +1
Mutation B (30% contribution): +2
D
DKO DKO DKO WT
405 412 422
MW
(kDa)
55
35
Cx43
GAPDH
Cx43
% vs. WT
WT
DKO405
DKO412
DKO422
Cx45
Cx45
GAPDH
120
100
80
60
40
20
0

**Supplemental Figure S6. A)** PCR of gDNA from the double knockout (DKO) HEK293FT cells showed one clean band indicating utility of the Cx43 and Cx45 primers (Cx43 expected amplicon size= 816bp; Cx45 expected amplicon size=647bp). Point mutations were introduced via CRISPR-Cas9 in the **B)** Cx43 and **C)** Cx45 sequences, adding respectively 1 specific nucleotide base (T) in Cx43 (Mutation A) and 2 specific nucleotide bases (AA) in Cx45 (Mutation B). Mutations are indicated by the red boxes. Those modifications induced a frameshift of the Cx43 and Cx45 DNA sequence**. D)** Western blotting analysis comparing three double knockout (DKO) HEK293FT cell lines to wild type (WT) HEK293FT cells. Note the absence of Cx43 (top row) and Cx45 bands (bottom row) in the DKO lines, but not in the WT cells. Specifically, Cx43 was decreased by 99.6%, 100%, and 99.4% in the DKO 405, 412 and 422 lines, respectively (top right); Cx45 was decreased by 98.5%, 100%, and 99% in the DKO 405, 412 and 422 lines, respectively (bottom right). Values were determined based on Cx43 or Cx45 / GAPDH. The DKO412 cell line was used for our subsequent recording analysis since it showed the strongest reduction of Cx43 and Cx45.

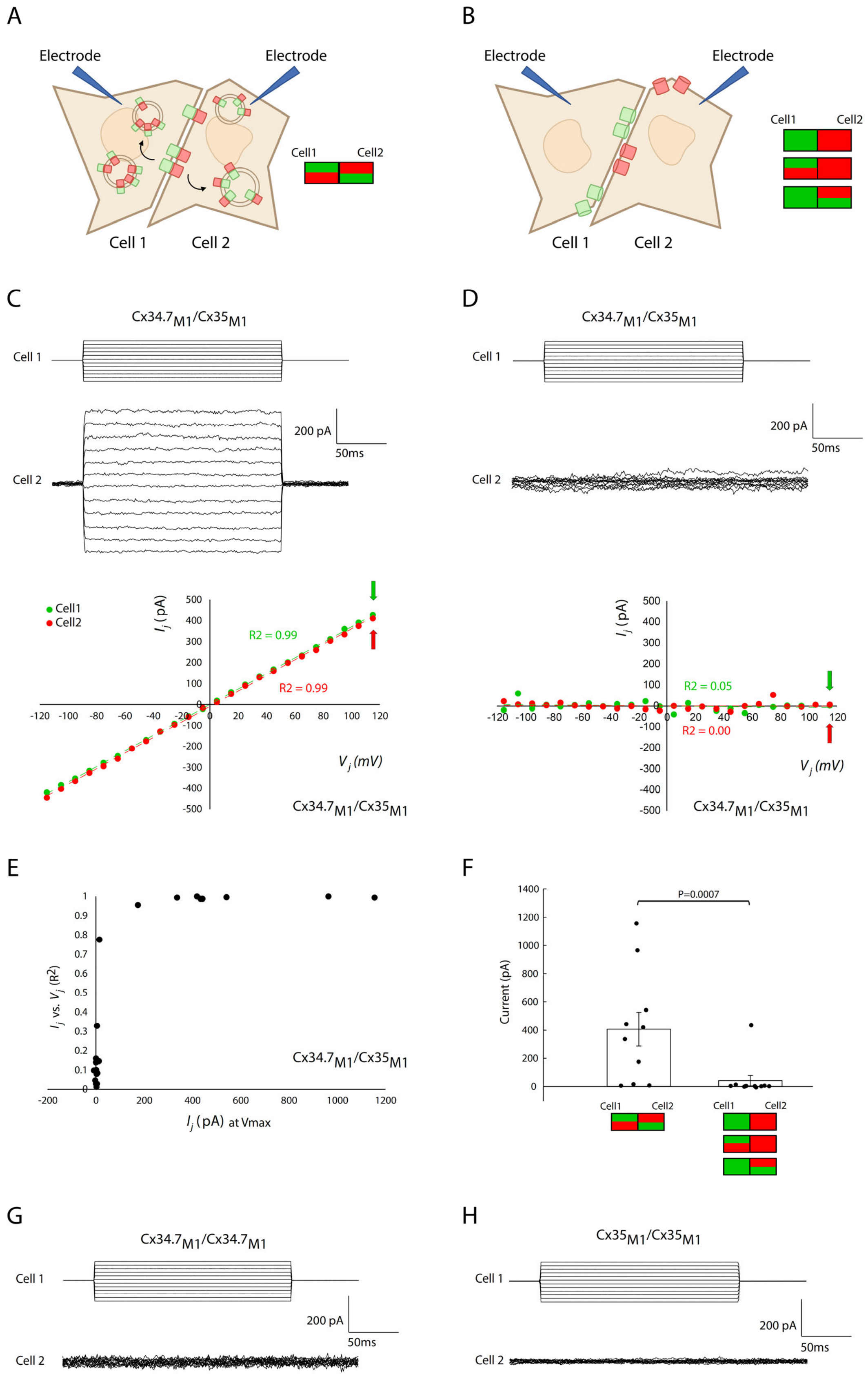
A
Electrode
Electrode
Cell1
Cell2
Cell 1
Cell 2
B
Electrode
Electrode
Cell1
Cell2
Cell 1
Cell 2
C
Cx34.7M1/Cx35M1
Cell 1
200 pA
50ms
Cell 2
Cell1
Cell2
Ij (pA)
R2 = 0.99
R2 = 0.99
Vj (mV)
Cx34.7M1/Cx35M1
D
Cx34.7M1/Cx35M1
Cell 1
200 pA
50ms
Cell 2
Ij (pA)
R2 = 0.05
R2 = 0.00
Vj (mV)
Cx34.7M1/Cx35M1
E
Ij vs. Vj (R2)
Cx34.7M1/Cx35M1
Ij (pA) at Vmax
F
P=0.0007
Current (pA)
Cell1
Cell2
Cell1
Cell2
G
Cx34.7M1/Cx34.7M1
Cell 1
200 pA
50ms
Cell 2
H
Cx35M1/Cx35M1
Cell 1
200 pA
50ms
Cell 2

**Supplemental Figure S7: Functional validation of Cx34.7M1-mEmerald and Cx35M1-mApple using Cx43/Cx45 double knockout HEK293FT cells. A)** We performed dual patch clamp recordings in Cx34.$7_{M1}$-mEmerald and Cx$35_{M1}$-mApple expressing cell pairs that both showed fluorescent patterns indicative of gap junction plaque internalization (red and green labeling in both cells), and **B)** fluorescently labeled HEK293FT double knockout pairs which did not. **C)** Representative injected and recorded current traces (top and middle), and the relationship between transjunctional current ($I_j$) and transjunctional voltage ($V_j$, bottom) for a cell pair with dual fluorescence internalization (same as Fig. 3I) and **D)** a cell pair where one cell expressed only one fluorescent protein (Pearson correlation values are shown for each cell within the pair). **E)** Current at the max transjunctional voltage vs. relationship between current flow and transjunctional voltage is shown for each cell pair (i.e., electrical coupling). Note that a current > 100pA at the max transjunctional voltage was indicative of electrical coupling of the cell pair. **F)** The max transjunctional voltage current was significantly higher for cell pairs that showed dual fluorescence internalization compared to pairs that did not (U =75, P=0.0007 using one tailed Wilcoxon rank sum test). **G-H)** Representative current traces for cell pairs that expressed G) Cx34.$7_{M1}$ or H) Cx$35_{M1}$ in a homotypic configuration. Injected and recorded currents are shown above and below, respectively. All current traces were filtered using Clampfit 11.4. They are depicted at a 20mV step and 1ms resolution.

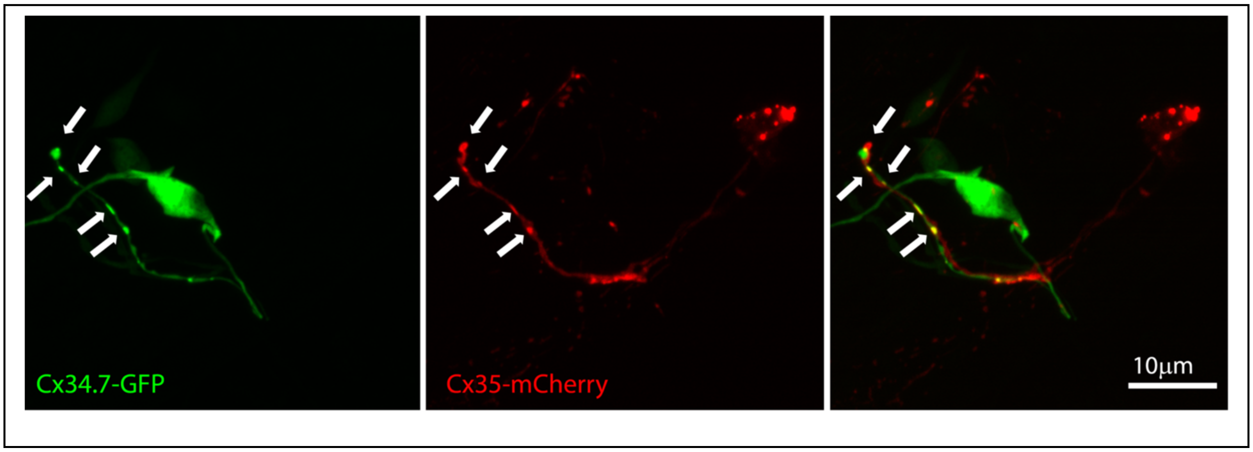


**Supplemental Figure S8: Confocal maximum intensity projections of *C. elegans* expressing the $Cx34.7_{M1}$/$Cx35_{M1}$ pair.** GFP-tagged $Cx34.7_{M1}$ is expressed in the AFD neuron, with puncta along its axon (left). mCherry-tagged $Cx35_{M1}$ is expressed in the AIY neuron, with puncta along its neurite (middle). Composite image showing the colocalizing GFP/mCherry puncta (right). White arrows highlight puncta. The representative image was selected from 30 acquired images.

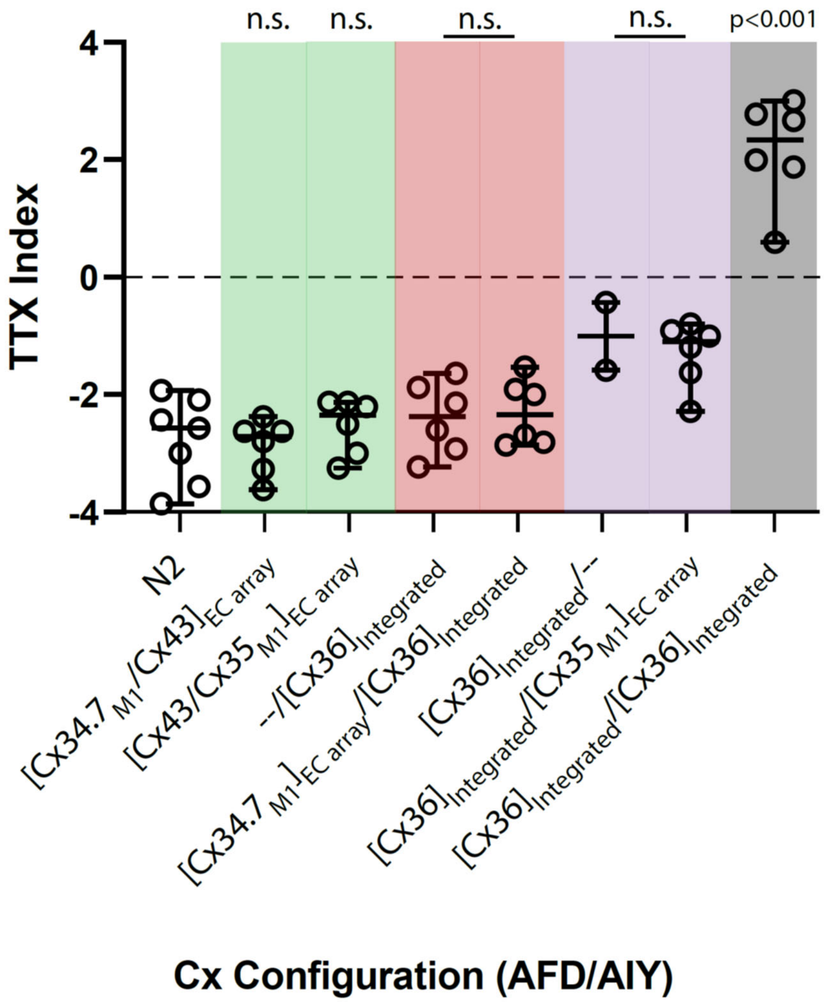


**Supplemental Figure S9: Connexin Cx34.7 and Cx35 do not alter *C. elegans* behavior when expressed as counterparts to connexins endogenous to the human brain.** We tested $Cx34.7_{M1}$ and $Cx35_{M1}$ hemichannels against Cx36 and Cx43. Worms with integrated arrays were used to test the mutant channels against Cx36. TTX Index (Y axis) refers to the thermotaxis index, calculated as previously described [3]. Negative values represent migration towards cold temperatures while positive values indicate migration towards warm temperatures. In the X-axis, N2 refers to the wild type *C. elegans* animals expressing no arrays. The other groups describe the expression of specific arrays in an AFD (presynaptic)/AIY (postsynaptic) configuration. "EC array" refers to extrachromosomal arrays, while "Integrated" refers to integrated arrays. Experimental worms were directly compared against integrated Cx36 controls, where $Cx34.7_{M1}$ or $Cx35_{M1}$ expression patterns mirrored our original experimental configuration. Worms expressing extrachromosomal arrays in AFD or AIY were used to test Cx43. C. elegans expressing $Cx34.7_{M1}$/Cx36, $Cx34.7_{M1}$/Cx43, Cx36/$Cx35_{M1}$ or Cx43/$Cx35_{M1}$ in AFD/AIY all continued to migrate towards cold temperatures ($F_{7,10.67}$= 19.29; P<0.0001 using Welch one-way ANOVA followed by Dunnett's T3 multiple comparisons; p<0.001), and no differences were observed across experimental pairs (P>0.05; highlighted by red or purple), indicating they were not forming functional electrical synapses.

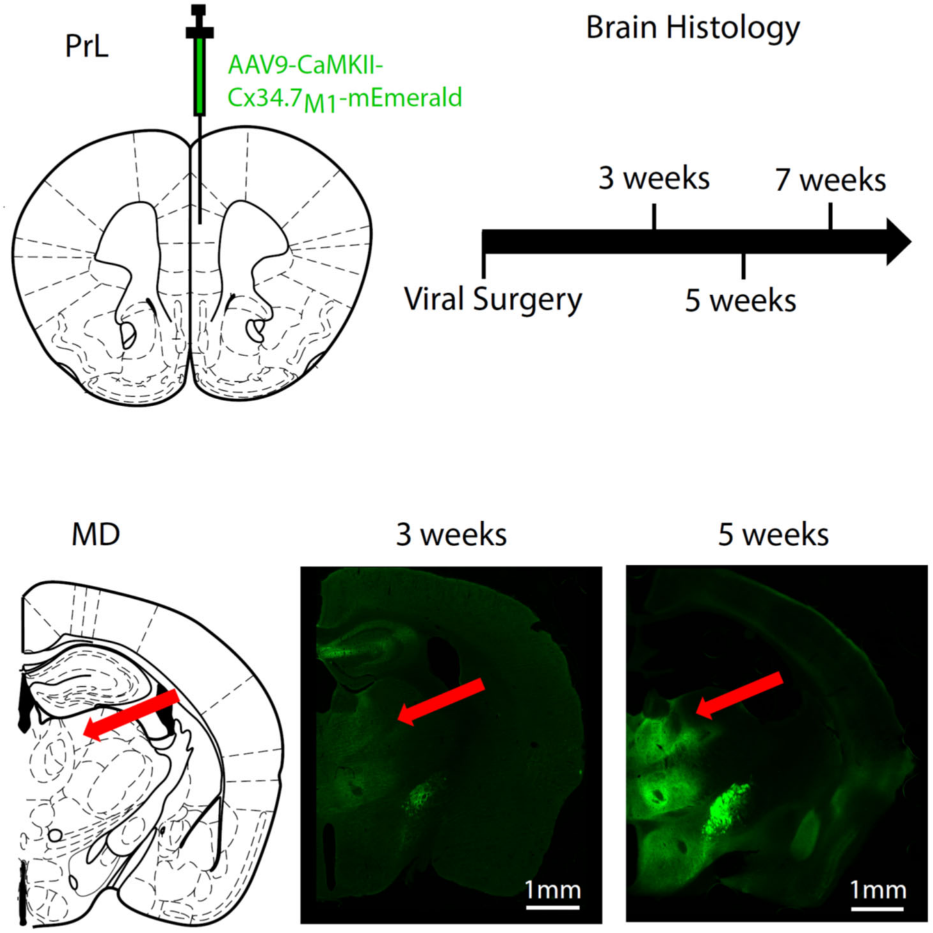


| | 3 weeks | 5 weeks | 7 weeks |
|---|---|---|---|
| Dorsal Striatum | ✓ | ✓ | ✓ |
| Nucleus Accumens | | ✓ | ✓ |
| Basolateral Amygdala | | ✓ | ✓ |
| Medial Dosal Thalamus | | ✓ | ✓ |
| Ventral Tegmental Area | | | ✓ |

**Supplemental Figure S10: Time course of pronounced Cx34.7 expression and trafficking in prelimbic cortex soma and terminals.** Five C57BL/6J mice were infected with AAV9-CaMKII-Cx34.7$_{M1}$-mEmerald in prelimbic cortex (PrL). Histology was performed 3 weeks (N=1 mouse), 5 weeks (N=2 mice), or 7 weeks (N=2 mice) after viral surgeries. Red arrows highlight the medial dorsal thalamus (MD) target for the tail suspension test experiments. Note the prominent trafficking of CaMKII-Cx34.7$_{M1}$-mEmerald apparent at 5 weeks (the time course utilized for our behavioral experiments). The table on the bottom indicates the post-surgical timeline at which prominent CaMKII-Cx34.7$_{M1}$-mEmerald expression was observed in each region.

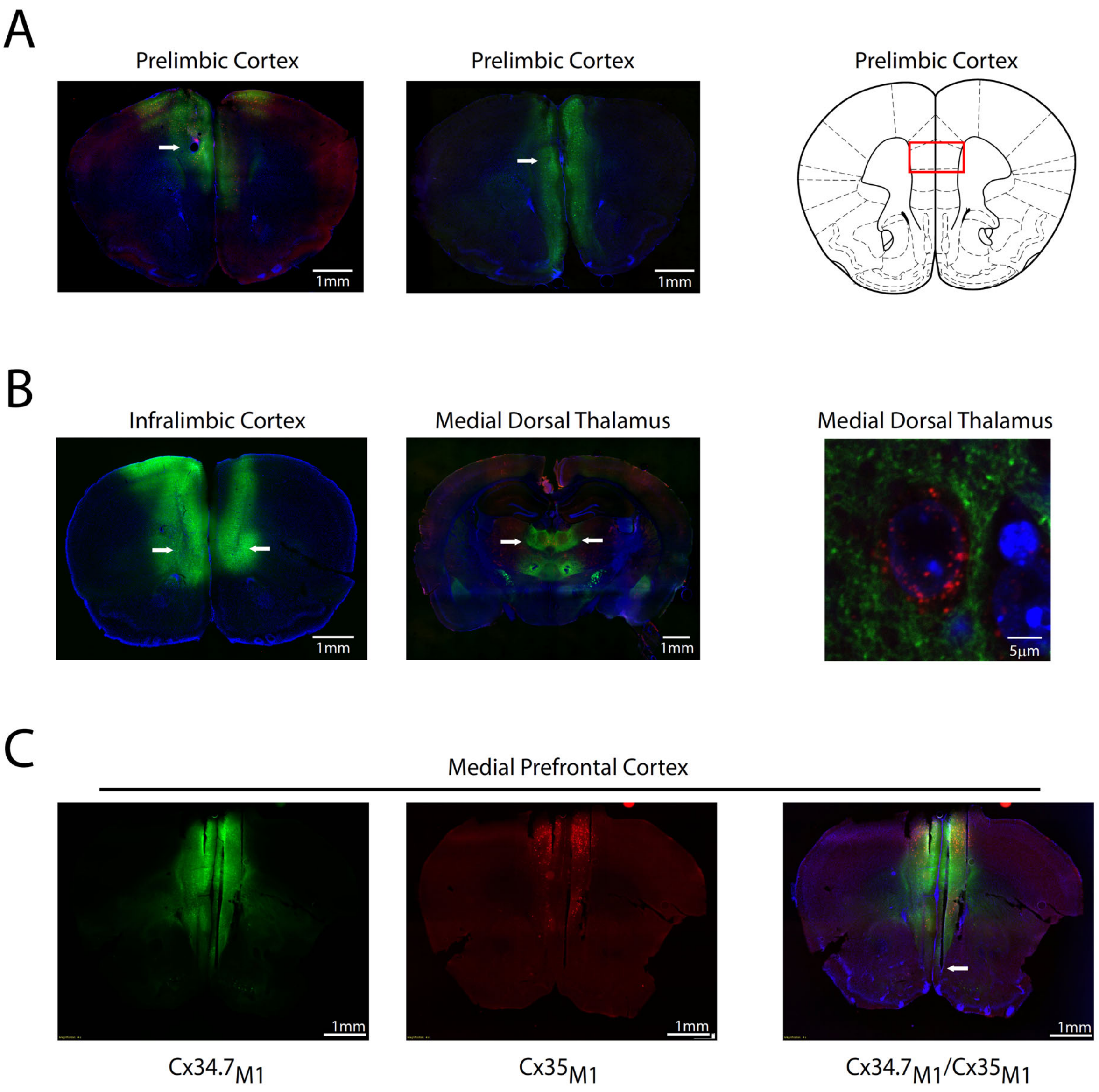


**Supplemental Figure S11: Representative histological images for mouse circuit-editing experiments. A)** Representative histological image of prelimbic cortex in a Cx34.7$_{M1}$/ Cx35$_{M1}$ infected mouse (left) and a Cx34.7$_{M1}$/ Cx34.7$_{M1}$ control (middle). White arrows highlight electrode tracks. For mice included in the prelimbic cortex editing experiment, individual electrode wires were distributed within the area highlighted by the red rectangle (right). Representative of N=26 mice **B)** Histological image of mouse injected with AAV9-CaMKII-Cx34.7$_{M1}$-mEmerald in infralimbic cortex (left). Image showing medial dorsal thalamus in mouse injected with AAV9-CaMKII-Cx34.7$_{M1}$-mEmerald in infralimbic cortex and AAV9-CaMKII-Cx35$_{M1}$-mApple in medial dorsal thalamus (middle). White arrows highlight viral injection tracks. Composite confocal image of medial dorsal thalamus showing expression of Cx35$_{M1}$-mApple (red) at the soma of a cell stained with DAPI (blue) and Cx34.7$_{M1}$-mEmerald (green) at IL nerve terminals (right). Representative of N=26 mice **C)** Histological images of a PV-Cre/Vglut2-flp mouse injected in medial prefrontal cortex with AAV9-CaMKII-Cx34.7$_{M1}$-mEmerald (left) and AAV9-DIO-Cx35$_{M1}$-T2A-mCherry (middle). Overlay is shown to the right. White arrow highlights the tip of one of the two shanks traversing the medial prefrontal cortex. Representative of N=6 mice.

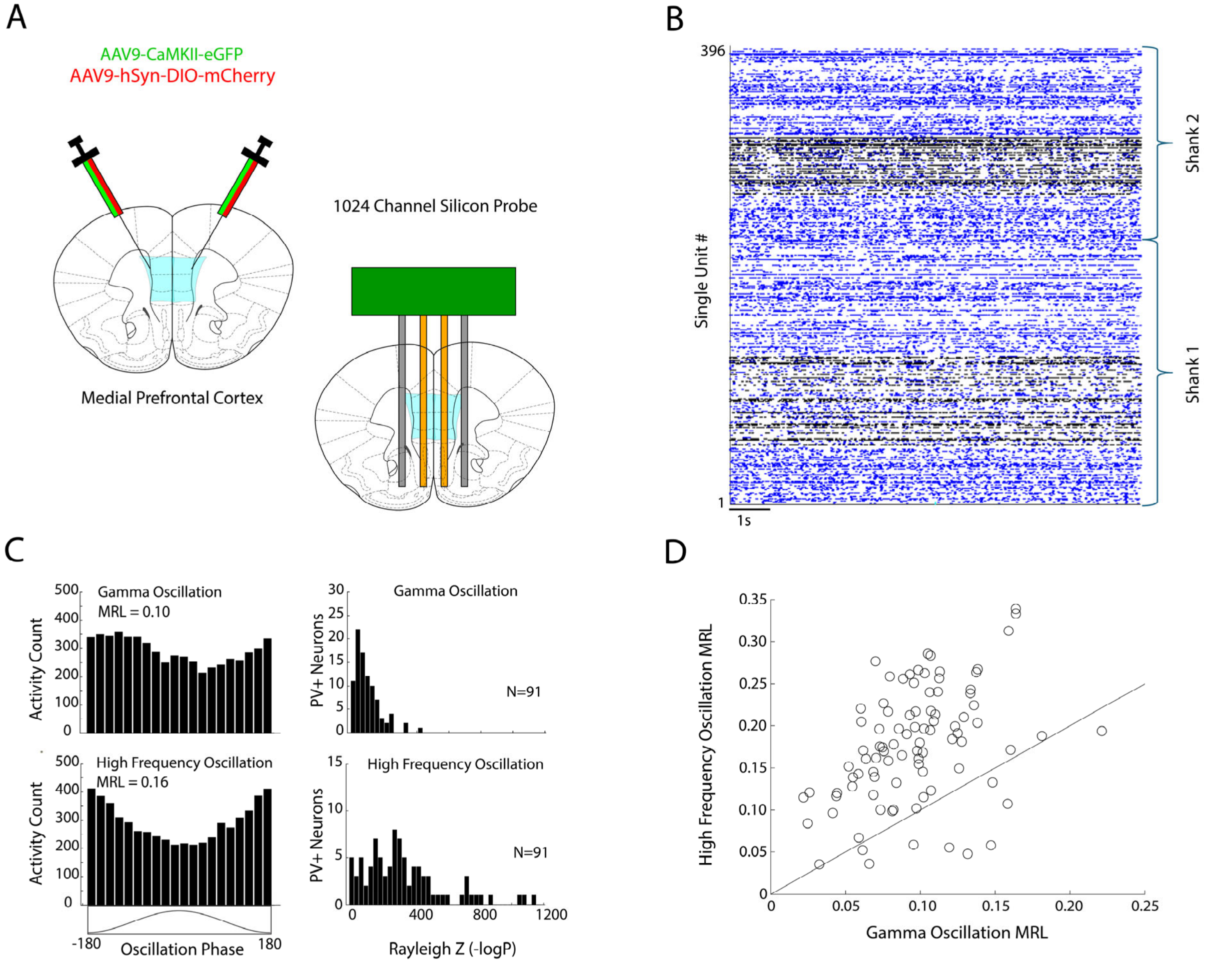


**Supplemental Figure S12: Medial Prefrontal Cortex (mPFC) PV+ interneurons optimally phase lock to high frequency oscillations. A)** Schematic for viral infection and silicon probe implantation targets in control mice used for analysis. **B)** Raster plot showing 396 single units recorded concurrently from two implanted shanks that traversed the mPFC in a female mouse. PV+ neurons from mPFC are depicted in black. **C)** Histograms depicting a mPFC PV+ neuron firing relative to the phase of gamma oscillations (30-80Hz; top left) and high frequency oscillations (80-200Hz; bottom left) recorded from the same channel as the neuron. Histogram showing phase locking of 91 PV+ neurons to gamma oscillations (top right) and high frequency oscillations (bottom right) are shown to the right. Data was acquired from two female mice and one male mouse. All the PV+ neurons showed phase locking to both oscillations. D) Mean resultant length (MRL) for all recorded PV+ neurons showing coupling to gamma and high frequency oscillations. The diagonal line corresponds to equivalent coupling to both oscillations. We observed significantly higher phase coupling to high frequency oscillations ($t_{90}$=12.2, P=$6\times10^{-21}$ for comparison of gamma and high frequency oscillation MRL using two-tailed paired t-test, N=91 neurons).

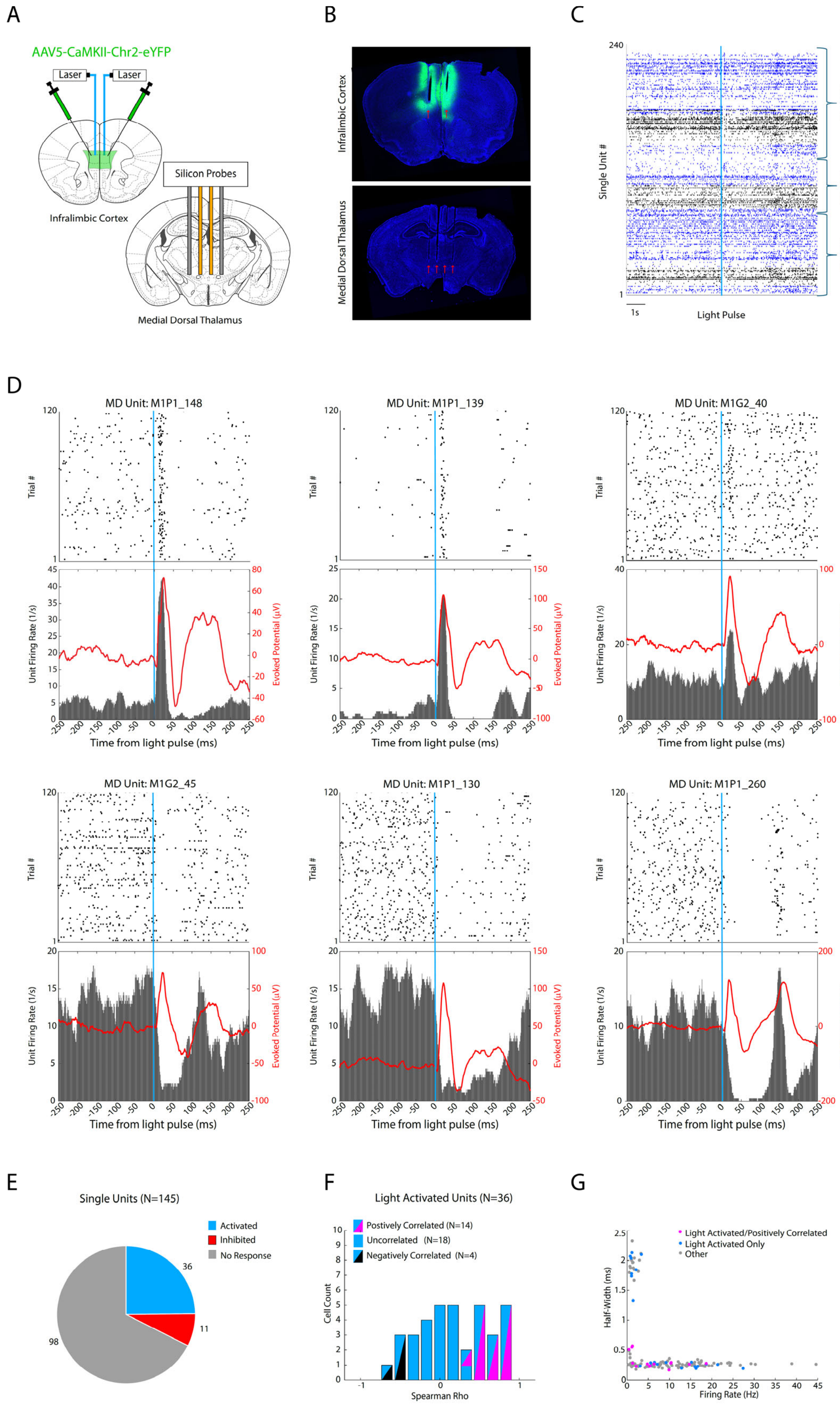

A
AAV5-CaMKII-Chr2-eYFP
Laser
Laser
Infralimbic Cortex
Silicon Probes
Medial Dorsal Thalamus
B
Infralimbic Cortex
Medial Dorsal Thalamus
C
Single Unit #
Shank 2
Shank 1
Shank 0
1s
Light Pulse
D
MD Unit: M1P1_148
MD Unit: M1P1_139
MD Unit: M1G2_40
MD Unit: M1G2_45
MD Unit: M1P1_130
MD Unit: M1P1_260
Trial #
Unit Firing Rate (1/s)
Evoked Potential (μV)
Time from light pulse (ms)
E
Single Units (N=145)
Activated
Inhibited
No Response
36
11
98
F
Light Activated Units (N=36)
Postively Correlated (N=14)
Uncorrelated (N=18)
Negatively Correlated (N=4)
Cell Count
Spearman Rho
G
Light Activated/Positively Correlated
Light Activated Only
Other
Half-Width (ms)
Firing Rate (Hz)

**Supplemental Figure S13: Medial dorsal thalamus (MD) cellular and local field potential responses to optogenetic stimulation of infralimbic cortex (IL). A)** Schematic of viral and electrical recording targeting approach. Two mice were infected with AAV5-ChR2-GFP in IL and implanted with a stimulating fiber in IL (bilaterally) and a 1024-channel silicon probe targeting MD. Activity was recorded from the two medial probes during quiet waking while mice were stimulated with blue light (1mW, 10ms pulse width, 493nm) 120 times with a pseudorandomized intertrial interval ranging from 8-24 seconds. **B)** Histological images showing ChR2-EYFP expression and optic fiber tracks in IL (red arrows, top), and individual electrode shank tracks in MD (red arrows, bottom). **C)** Raster plot showing 240 single units recorded concurrently from three implanted shanks that traversed MD. Cellular firing is shown relative to a light pulse in IL (light blue vertical line), and units from MD neurons are depicted in black. **D)** Examples of single units recorded from MD that showed a change in their activity following a light pulse. A raster plot depicting firing of each neuron relative to the 120 light pulses is shown on top. The average firing rate across all the trials is shown on the bottom for each neuron, and the mean LFP activity recorded from the same electrode channel is overlaid in red. Note that the three neurons in the top row showed an increase in their activity following a light pulse, while the three neurons on the bottom row showed a decrease in their activity. **E)** Light stimulation increased the activity of 36 of the 145 single neurons isolated from MD and inhibited the activity of 11 MD neurons. **F)** Activity of the 36 MD neurons that were induced by light stimulation was compared to the evoked potential measured in the same channel. 14/36 of these neurons showed activity that was directly correlated with the evoked potential, confirming its local relevance. 6/36 neurons showed activity that was inversely correlated with the evoked potential. G) Firing rate and waveform properties for the 145 MD neurons. The 36 neurons that were activated by light stimulation are highlighted in pink (positively correlated with the evoked potential), or blue (not positively correlated with the evoked potential). The remaining neurons in gray were either suppressed or not modulated by light.

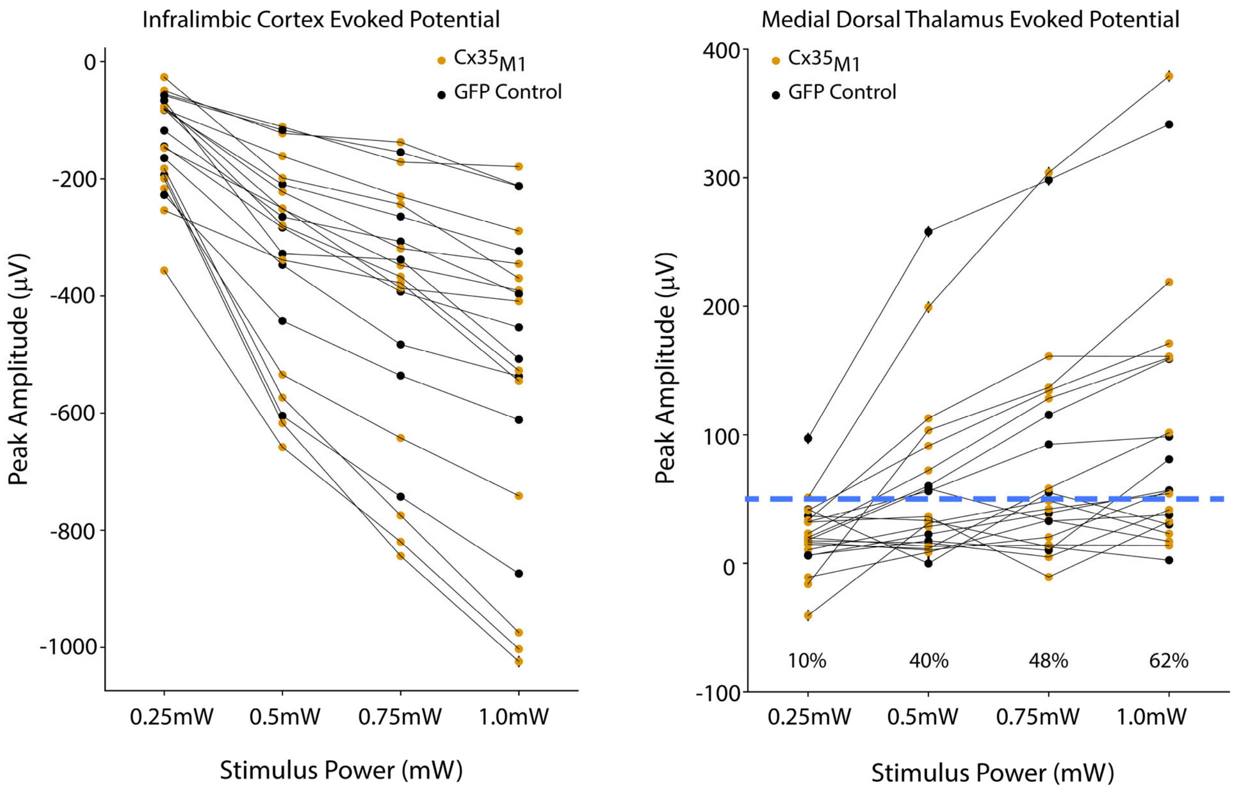


**Supplemental Figure S14: Cortical and thalamic potentials evoked by IL stimulation during first experimental session.** Peak cortical amplitudes within 10ms of stimulation are shown to the left. Responses were averaged across microwires. Peak thalamic amplitudes within 25ms are shown the right. The percentage of mice that showed thalamic evoked responses above 50mV (indicated by dashed blue line) is shown for each light intensity. Each line represents one mouse; N=21 total mice.

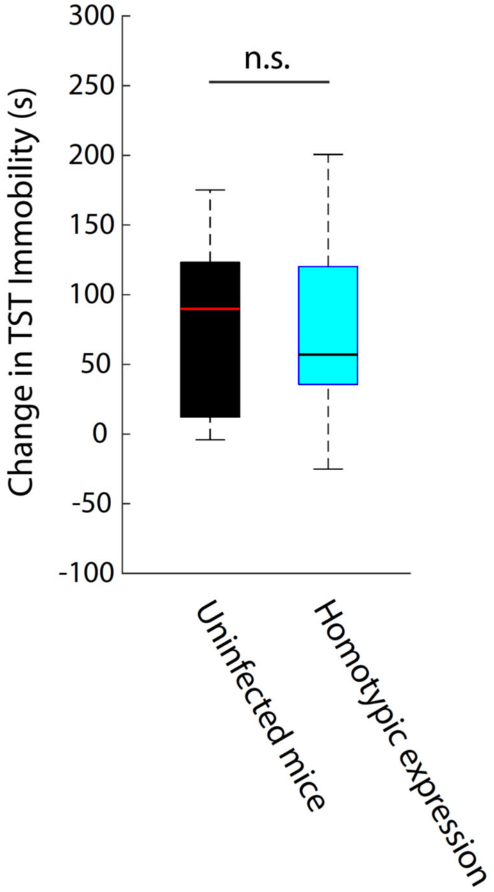


**Supplemental Figure S15: Homotypic expression of $Cx34.7_{M1}$ or $Cx35_{M1}$ across the IL→MD circuit does not impact stress-induced behavioral adaptation in the tail suspension test (TST) relative to uninfected BALB/cJ mice ($T_{29}$=0.16, P=0.87 using two tailed t-test).**

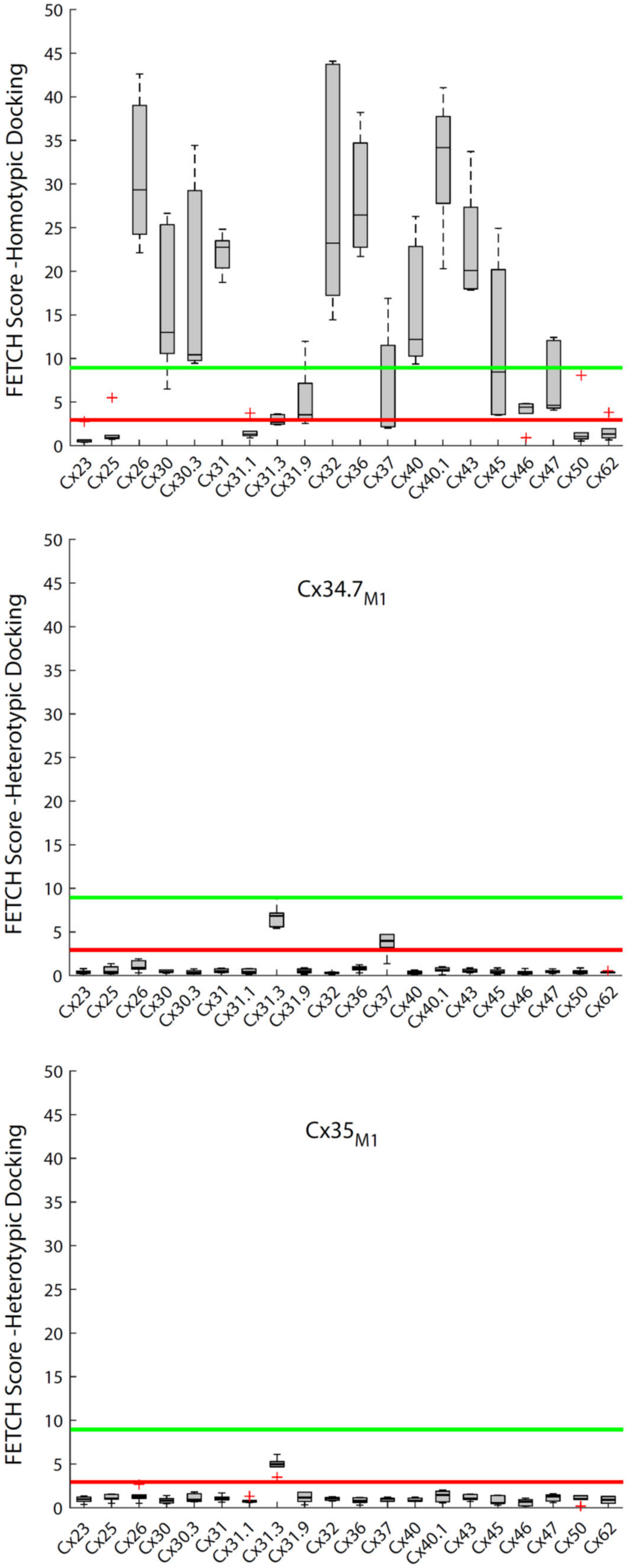

FETCH Score -Homotypic Docking
0
5
10
15
20
25
30
35
40
45
50
Cx23
Cx25
Cx26
Cx30
Cx30.3
Cx31
Cx31.1
Cx31.3
Cx31.9
Cx32
Cx36
Cx37
Cx40
Cx40.1
Cx43
Cx45
Cx46
Cx47
Cx50
Cx62
$Cx34.7_{M1}$
FETCH Score -Heterotypic Docking
$Cx35_{M1}$
FETCH Score -Heterotypic Docking

**Supplemental Figure S16. Docking compatibility of $Cx34.7_{M1}$ and $Cx35_{M1}$ with human connexins.** We used FETCH to quantify the homotypic docking characteristics of 20 human connexin isoforms (top). We then characterized the heterotypic docking characteristic of our mutant proteins with the 20 endogenous isoforms (middle and bottom). The red and green lines correspond to the upper and lower limits of non-docking and docking population shown in Fig. 1C, respectively. Notably, expression of our Cx59 constructs appeared to promote cell death and was thus excluded from our studies. Additionally, we were unable to establish that Cx23, Cx25, Cx31.1, Cx46, Cx50, and Cx62 were amenable to FETCH analysis given very low homotypic FETCH scores. We did not perform statistical analysis across all the pairs shown here. Rather, we provide this data to highlight potential interactions that our engineered proteins may exhibit with other endogenous connexins, and to guide future experiments. For each Cx, the central mark is the median, the edges of the box are the 25th and 75th percentiles, the whiskers extend to the most extreme datapoints the algorithm considers to be not outliers, and the outliers are plotted individually as a "+" (MATLAB, The MathWorks, Inc., Natick, MA).
